# Small subpopulations of β-cells do not drive islet oscillatory [Ca^2+^] dynamics via gap junction communication

**DOI:** 10.1101/2020.10.28.358457

**Authors:** JaeAnn M. Dwulet, Jennifer K. Briggs, Richard K.P. Benninger

## Abstract

The islets of Langerhans exist as a multicellular network that is important for the regulation of blood glucose levels. The majority of cells in the islet are insulin-producing β-cells, which are excitable cells that are electrically coupled via gap junction channels. β-cells have long been known to display heterogeneous functionality. However, due to gap junction electrical coupling, β-cells show coordinated [Ca^2+^] oscillations when stimulated with glucose, and global quiescence when unstimulated. Small subpopulations of highly functional β-cells have been suggested to control the dynamics of [Ca^2+^] and insulin release across the islet. In this study, we investigated the theoretical basis of whether small subpopulations of β-cells can disproportionality control islet [Ca^2+^] dynamics. Using a multicellular model of the islet, we generated continuous or bimodal distributions of β-cell heterogeneity and examined how islet [Ca^2+^] dynamics depended on the presence of cells with increased excitability or increased oscillation frequency. We found that the islet was susceptible to marked suppression of [Ca^2+^] when a ∼10% population of cells with high metabolic activity was hyperpolarized; where hyperpolarizing cells with normal metabolic activity had little effect. However, when these highly metabolic cells were removed from the islet model, near normal [Ca^2+^] remained. Similarly, when ∼10% of cells with either the highest frequency or earliest elevations in [Ca^2+^] were removed from the islet, the [Ca^2+^] oscillation frequency remained largely unchanged. Overall these results indicate that small populations of β-cells with either increased excitability or increased frequency, or signatures of [Ca^2+^] dynamics that suggest such properties, are unable to disproportionately control islet-wide [Ca^2+^] via gap junction coupling. As such, we need to reconsider the physiological basis for such small β-cell populations or the mechanism by which they may be acting to control normal islet function.

**Author summary:** Many biological systems can be studied using network theory. How heterogeneous cell subpopulations come together to create complex multicellular behavior is of great value in understanding function and dysfunction in tissues. The pancreatic islet of Langerhans is a highly coupled structure that is important for maintaining blood glucose homeostasis. β-cell electrical activity is coordinated via gap junction communication. The function of the insulin-producing β-cell within the islet is disrupted in diabetes. As such, to understand the causes of islet dysfunction we need to understand how different cells within the islet contribute to its overall function via gap junction coupling. Using a computational model of β-cell electrophysiology, we investigated how small highly functional β-cell populations within the islet contribute to its function. We found that when small populations with greater functionality were introduced into the islet, they displayed signatures of this enhanced functionality. However, when these cells were removed, the islet, retained near-normal function. Thus, in a highly coupled system, such as an islet, the heterogeneity of cells allows small subpopulations to be dispensable, and thus their absence is unable to disrupt the larger cellular network. These findings can be applied to other electrical systems that have heterogeneous cell populations.

## Introduction

Many tissues exist as multicellular networks that have complex structures and functions. Multicellular networks are generally comprised of heterogenous cell populations, and heterogeneity in cellular function makes it difficult to understand the underlying network behavior. Studying the constituent cells individually is of value. However, understanding how heterogeneous cell populations come together to form a coherent structure with emergent properties is important to understand what leads to dysfunction in these networks [1]. The multicellular pancreatic islet lends itself to network theory with its distinct architecture, cellular heterogeneity, and cell-cell interactions.

The pancreatic islet is a micro-organ that helps maintain blood glucose homeostasis [2]. Death or dysfunction to insulin-secreting β-cells within the islet generally causes diabetes [3]. When blood glucose levels rise, glucose is transported into the β-cell and phosphorylated by glucokinase (GK), the rate limiting step of glycolysis [4-6]. Following glucose metabolism, the ratio of ATP/ADP increases, closing ATP sensitive K^+^ channels (K_ATP_). K_ATP_ channel closure causes membrane depolarization, opening voltage gated Ca^2+^ channels and elevating intra-cellular free-calcium ([Ca^2+^]); which triggers insulin granule fusion and insulin release [7, 8]. Disruptions to this glucose stimulated insulin secretion pathway occur in diabetes [9-11]. β-cells are electrically coupled by connexin36 (Cx36) gap junctions which can transmit depolarizing currents across the islet that synchronize oscillations in [Ca^2+^]. Under low glucose conditions, gap junctions transmit hyperpolarizing currents that suppress islet electrical activity [12-15]. Understanding the role cell-cell communication between β-cells plays can increase our understanding of dysfunction to islet dynamics during the pathogenesis of diabetes.

Despite their robust coordinated behavior within the intact islet, β-cells are functionally heterogeneous [16]. Individual β-cells show heterogeneity in expression of GK [17], glucose metabolism [16], differing levels of insulin production and secretion [18-21], and faster and irregular [Ca^2+^] oscillations when compared to whole islet oscillations [22]. Various cell surface and protein markers have been used to identify subpopulations of β-cells with differences in functionality and proliferative capacity [23-27]. Nevertheless, the importance of β-cell heterogeneity and how these subpopulations contribute to islet function is poorly understood.

While many studies of β-cell heterogeneity have been performed in dissociated cells, a few studies have investigated the role of heterogeneity in the intact islet [28]. In one study, following stimulation via the optogenetic cationic channel channelrhodopsin (ChR2), ∼10% of β-cells were found to be highly excitable and able to recruit [Ca^2+^] elevations in large regions in the islet. These highly excitable cells had higher metabolic activity [29]. In another study, the optogenetic Cl^-^ pump halorhodopsin (eNpHr3) was used to silence β-cells. A population of ∼1-10% “hub” β-cells was discovered that when hyperpolarized by eNpHr3 substantially disrupted coordinated [Ca^2+^] dynamics across the islet. These cells had increased GK expression [30]. In related studies, a small population of cells showed [Ca^2+^] oscillations that consistently preceded the rest of the islet and were suggested to be ‘pacemaker cells’ that drove islet [Ca^2+^] dynamics [31]. Theoretically, how small subpopulations of cells may be capable of driving elevations and oscillatory dynamics of [Ca^2+^] across the islet is not well established, and has been a significant topic of debate [32, 33].

In this study we explore the theoretical basis for whether small β-cell subpopulations can control multicellular islet [Ca^2+^] dynamics. Towards this, we utilize a computational model of the islet that we have previously validated against a wide-range of experimental data [29, 34-36]. This includes understanding how populations of inexcitable cells suppress islet function and the role for electrical coupling. We investigate whether small populations of highly excitable cells or cells with high frequency oscillations can respectively drive the elevations or dynamics of islet [Ca^2+^] oscillations. This includes simulating the removal of specific cell populations within the context of broad continuous distributions or distinct bimodal distributions of heterogeneity.

## Results

### How variation in metabolic activity impacts islet function

Experimental evidence indicates that within the intact islet there exists 10-20% variation in metabolic activity [37]. Previous modelling studies have represented beta cell heterogeneity as a continuous distribution with 10-25% variation in GK activity and metabolic activity, which is sufficient to model the impact of electrical coupling and heterogeneity within the islet [29, 34-36]. However, recent experimental evidence has suggested that small β-cell subpopulations with elevated metabolic activity or GK expression are present within the islet and may disproportionately drive elevated [Ca^2+^] [29, 30]. For example, ‘hub’ β-cells show increased connectivity (synchronized Ca^2+^ oscillations) and increased GK expression compared to the rest of the islet [30]. When these ‘hub’ cells were hyperpolarized, [Ca^2+^] is suppressed in large regions of the islet; whereas hyperpolarizing other cells had little impact.

We first asked whether identification of such a ‘hub’ subpopulations may arise as part of the natural variation within a continuous distribution. We simulated an islet with a continuous distribution in GK activity (Fig 1a), and targeted hyperpolarization to a population of cells based on their GK activity. Simulated islets had normal synchronized [Ca^2+^] oscillations (Fig 1b), comparable to previous studies [29, 34-36, 38, 39]. When hyperpolarization was targeted to a random set of cells across the islet, near-normal [Ca^2+^] activity was maintained until greater than 20% of cells within the islet were targeted (Fig 1b,c). Above this level, the islet lacked significant [Ca^2+^] elevations (Fig 1c), consistent with prior measurements [34, 36]. When hyperpolarization was targeted specifically to cells with either higher GK (GK^Higher^) or lower GK (GK^Lower^), similar changes in [Ca^2+^] activity were observed as with targeting a random subset of cells: the islet retained near-normal [Ca^2+^] activity until greater than 20% of these GK^Higher^ or GK^Lower^ cells were targeted (Fig 1c). Nevertheless when 20% of cells were hyperpolarized, targeting GK^Higher^ cells did result in silencing of significantly more of the islet compared to GK^Lower^ cells. Within the simulated islet we also decoupled and removed the same GK^Higher^ or GK^Lower^ populations. In this case, the remaining islet showed normal elevations in [Ca^2+^], with little to no difference between removing GK^Higher^ or GK^Lower^ cells (Fig 1d).

**Figure 1.**
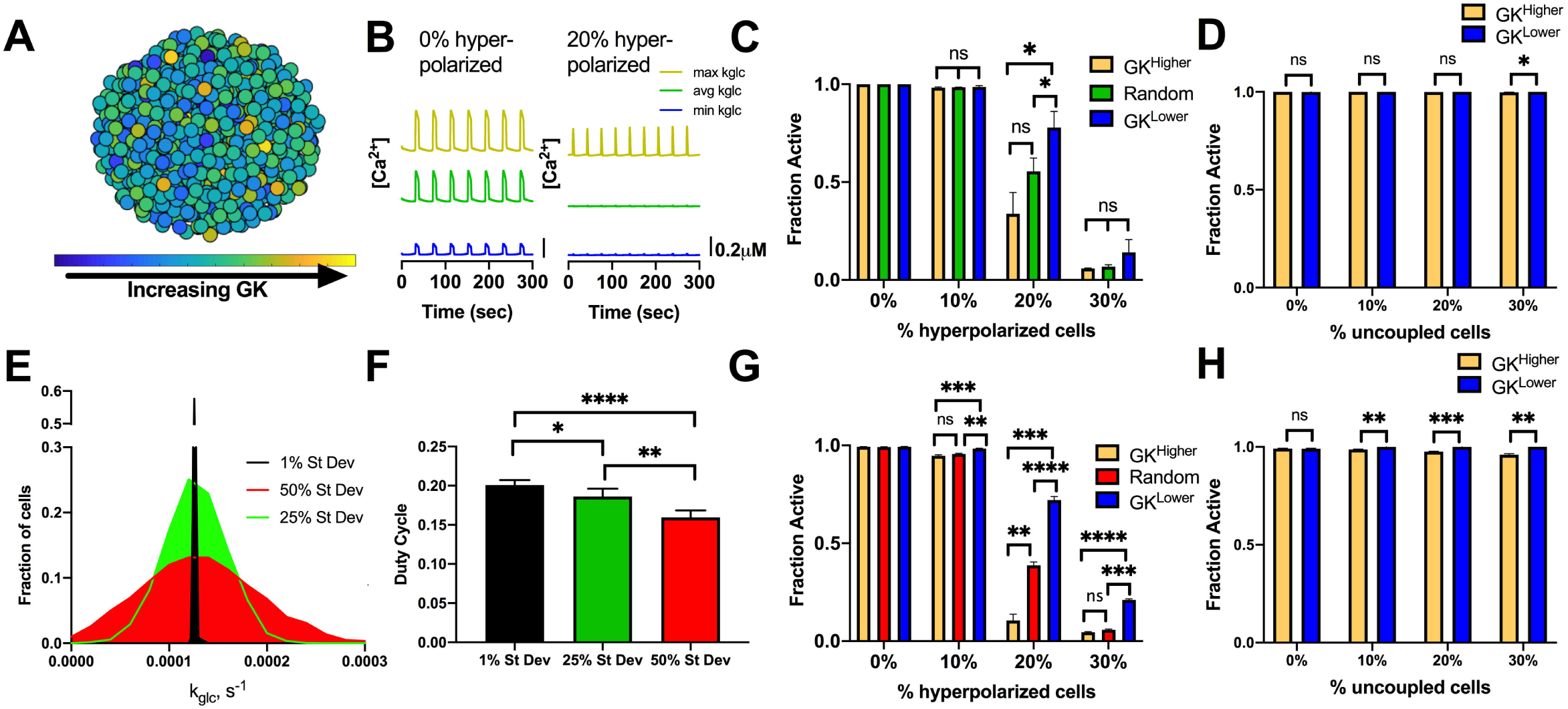
Simulations predicting how variation in GK activity impact islet function. A). Schematic of continuous distribution of heterogeneous GK activity across simulated islet with 25% variation in GK rate (k_glc_). B). Representative time courses of [Ca^2+^] for 3 cells in simulated islet in A. Left is simulation with 0% hyperpolarized cells and right is simulation with a random 20% of cells hyperpolarized. Blue trace is cell with lowest GK rate (k_glc_), Green is cell with the average GK rate, yellow is cell with the highest GK rate. C). Fraction of cells showing elevated [Ca^2+^] activity (active cells) in simulated islets vs. the percentage of cells hyperpolarized in islet. Hyperpolarized cells are chosen based on their GK rate. D). Fraction of active cells in islet when cells are uncoupled from the rest of the cells in the simulation. E). Histogram showing average frequency of cells at varying GK rate (k_glc_) for simulations that have different standard deviation in GK activity. F). Average duty cycle of cells from simulations with different standard variation in GK activity. G). As in C. for simulations with standard deviation in GK activity at 50% of the mean. H). As in D. for simulations with standard deviation in GK activity at 50% of the mean. Error bars are mean ± s.e.m. Repeated measures one-way ANOVA with Tukey post-hoc analysis was performed for simulations in C and G, Student’s paired t-test was performed for D and H, and one-way ANOVA was performed for F to test for significance. Significance values: ns indicates not significant (p>.05), * indicates significant difference (p<.05), ** indicates significant difference (p<.01), *** indicates significant difference (p<.001), **** indicates significant difference (p<.0001). Data representative of 5 simulations with differing random number seeds.

Given uncertainty in the exact level of heterogeneity within the islet, we next tested whether changes to the variability in GK could lead to differences in [Ca^2+^] upon targeting cells with higher GK (GK^Higher^) or lower GK (GK^Lower^) cells. We simulated islets with decreased variation in GK activity (1% variation) or increased variation in GK activity (50% variation) and compared [Ca^2+^] with our previous simulations of 25% variation (Fig 1e and S1a Fig). The duty cycle of the simulated islets slightly decreased as the GK variation increased (Fig 1f), but [Ca^2+^] oscillations remained across the islet that closely matched previous studies. Under 50% variation in GK, when hyperpolarization was targeted to a random set of cells across the islet, the islet retained near-normal [Ca^2+^] activity until greater than 20% of the islet was targeted, as before. In contrast, when hyperpolarization was targeted specifically to cells with higher GK (GK^Higher^), [Ca^2+^] was largely abolished for greater than 10% of cells being targeted (Fig 1g). However, when hyperpolarization was targeted to lower GK (GK^Lower^) cells, [Ca^2+^] was largely unchanged until 30% of cells were targeted (Fig 1g). As such, upon hyperpolarizing 20% of cells, a substantial difference in [Ca^2+^] resulted from targeting GK^Higher^ or GK^Lower^ cells. Nevertheless, when these higher GK or lower GK cells were decoupled and removed from the islet, the impact on [Ca^2+^] elevations was very minor. A minor 2-4% decrease in [Ca^2+^] occurred when removing >10% GK^Higher^ cells, with no impact when removing GK^Lower^ cells (Fig 1h).

We also tested whether changing other properties of cells with higher GK or lower GK would impact the suppression of [Ca^2+^]. When GK activity correlated with gap junction conductance such that higher GK cells also had increased gap junction conductance (GK^Higher^/g _Coup_ ^Higher^), little impact was observed (S2a-c Fig). However, when GK activity negatively correlated with K_ATP_ conductance, such that higher GK cells had increased gap junction conductance, but also had reduced K_ATP_ conductance, no difference occurred when hyperpolarizing higher GK (GK^Higher^/g _Coup_ ^Higher^/g_KATP_ ^Lower^) or lower GK (GK^Lower^/g _Coup_ ^Lower^/g _KATP_ ^Higher^) cells (S2d-f Fig).

Thus, hyperpolarizing a small sub-population of metabolically active cells can disproportionately suppress islet [Ca^2+^], particularly when heterogeneity is very broad. However, when these same cells are removed or absent from the islet, the impact on [Ca^2+^] is minimal under the model assumptions set here.

### Impact of a bimodal distribution of functional β-cell subpopulations

We next examined how imposing a bimodal distribution in GK activity would impact targeting hyperpolarization to a small population of metabolically active cells. We simulated an islet with a population of highly metabolic cells that comprised 10% of the islet (GK^High^) (Fig 2a). To maintain normal average GK activity, the rest of the islet had slightly reduced GK activity (GK^Low^) (Fig 2b). Gap junction coupling conductance of all cells remained unchanged (S1b Fig). Under this bimodal distribution, the islet displayed regular [Ca^2+^] oscillations at high glucose that closely matched previous simulations (Fig 2c). We tested the effect of targeting hyperpolarization to either the GK^High^ or GK^Low^ cell populations. When all GK^High^ cells (10%) were hyperpolarized, [Ca^2+^] was fully suppressed across the islet. Conversely, when GK^Low^ cells (10%) were hyperpolarized, [Ca^2+^] remained largely unchanged (Fig 2d). However, when a greater proportion of GK^Low^ cells (20%) were hyperpolarized, [Ca^2+^] was suppressed, as with a continuous distribution under 50% variation. These results show very good agreement between prior experiments where very different [Ca^2+^] response was observed when hyperpolarizing cells with higher GK and cells with lower GK.

**Figure 2.**
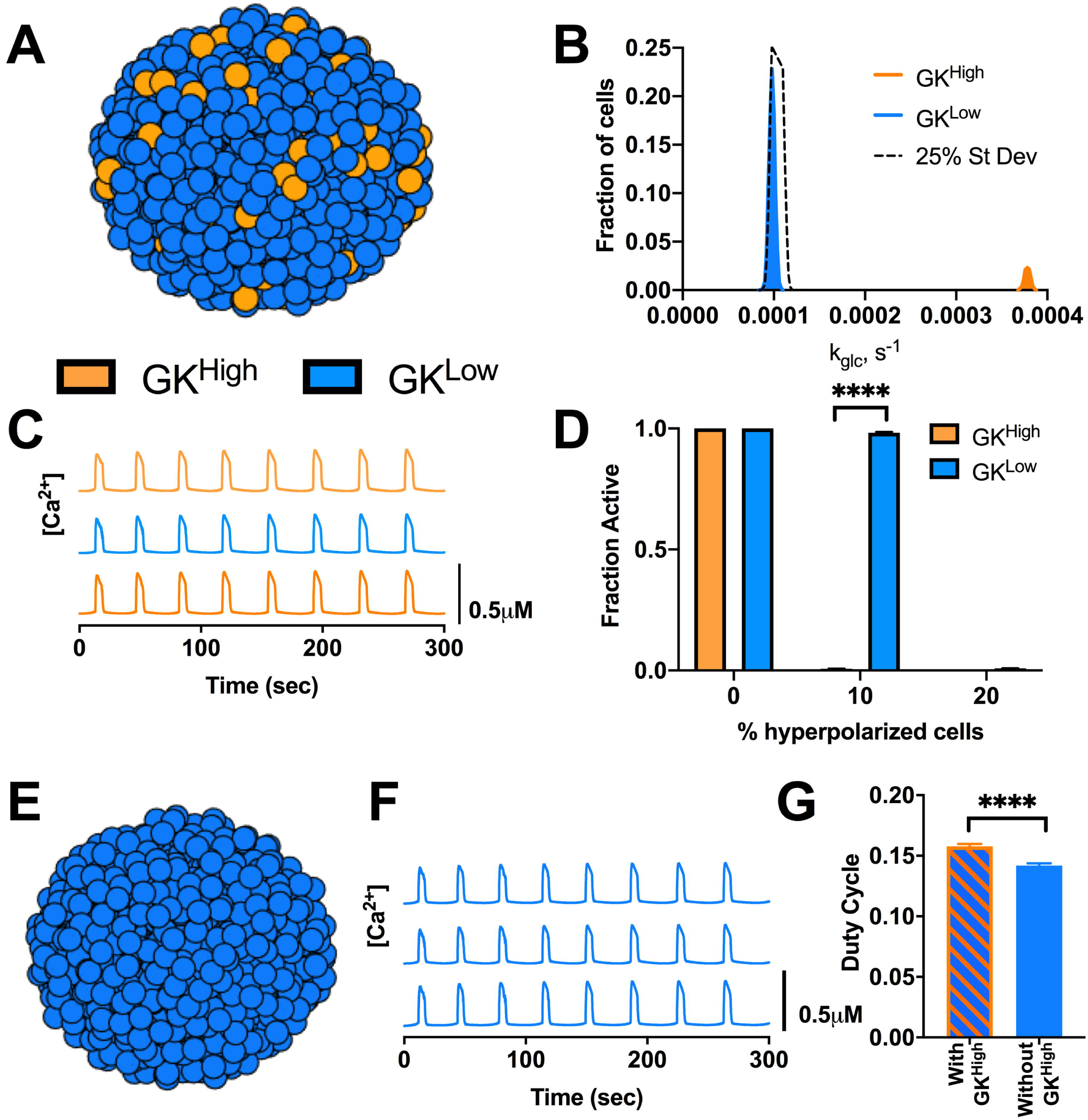
Bimodal distribution in GK activity predicts small highly functional cells are dispensable for islet [Ca^2+^] dynamics. A). Schematic of bimodal distribution of GK activity across simulated islet. B). Histogram showing average frequency of cells at varying GK rate (k_glc_) for bimodal compared with continuous distribution (25% St Dev). C). Representative time courses of [Ca^2+^] for 3 cells in simulated bimodal islet in A. Blue traces are cells from GK^Low^ population and orange traces are cells from GK^High^ population. D). Fraction of cells showing elevated [Ca^2+^] activity (active cells) in bimodal simulations vs. the percentage of cells hyperpolarized in islet. Hyperpolarized cells are chosen either from GK^High^ (orange bars) or GK^Low^ (blue bars) population. E). Schematic of simulation where only GK^Low^ cells are present and no GK^High^ cells are included. F). Representative time courses of [Ca^2+^] for 3 cells in simulated islet in E. G). Average duty cycle of cells from simulations of a bimodal model as in A (With GK^High^) and from simulations as in E (Without GK^High^). Error bars are mean ± s.e.m. Student’s paired t-test was performed to test for significance. Significance values: ns indicates not significant (p>.05), * indicates significant difference (p<.05), ** indicates significant difference (p<.01), *** indicates significant difference (p<.001), **** indicates significant difference (p<.0001). Data representative of 4 simulations with differing random number seeds.

We next tested whether the cells from the highly metabolic population (GK^High^) are important to support islet function, by simulating an islet with only cells from the lower GK population (GK^Low^) (Fig 2e). In this context, the islet retained near-normal [Ca^2+^] activity (Fig 2f), with a minor drop in duty cycle (Fig 2g). As such, the simulated islet was capable of functioning near-normally in the absence of a small (∼10%) highly metabolic subpopulation. Thus, despite showing substantial differences in islet activity when hyperpolarized, a small metabolically active subpopulation is not required to maintain elevations in oscillatory [Ca^2+^] across the islet.

### How variations in gap junction coupling impact functional β-cell subpopulations

Metabolically active subpopulations of cells that disproportionately control the islet have been suggested to have increased connectivity [30]. We next examined how changes in gap junction electrical coupling affect how targeting hyperpolarization to specific cell populations impacts islet [Ca^2+^]. We simulated the islet with the same bimodal distribution in GK activity as in Fig 2, but correlated gap junction coupling conductance (g_Coup_) with GK activity (k_glc_) across the islet (Fig 3a and S1c Fig). As such, more metabolically active GK^High^ cells had ∼2 times higher coupling conductance than that of the population of cells with lower metabolic activity (GK^Low^ cells). Under this model, when the highly metabolic population, GK^High^ cells, were targeted with hyperpolarization, the islet retained some [Ca^2+^] elevations (∼45%) (Fig 3b). When cells with less metabolic activity, GK^Low^, cells were targeted with hyperpolarization, the islet showed little change in [Ca^2+^] activity, as before. As such, the suppression of [Ca^2+^] upon targeting hyperpolarization to highly metabolic cells is reduced by those cells having elevated electrical coupling, (Fig 3c). We further simulated the islet with a bimodal distribution in both GK activity and gap junction conductance, such that more metabolically active GK^High^ cells had increased their coupling conductance by ∼3 times (Fig 3d and S1d Fig). Under these conditions when highly metabolic cells were targeted with hyperpolarization, the islet retained substantial [Ca^2+^] elevations (∼60%) (Fig 3e,f). Thus, increasing gap junction coupling does not enhance the ability of metabolically active cells to maintain oscillatory islet [Ca^2+^] elevations.

**Figure 3.**
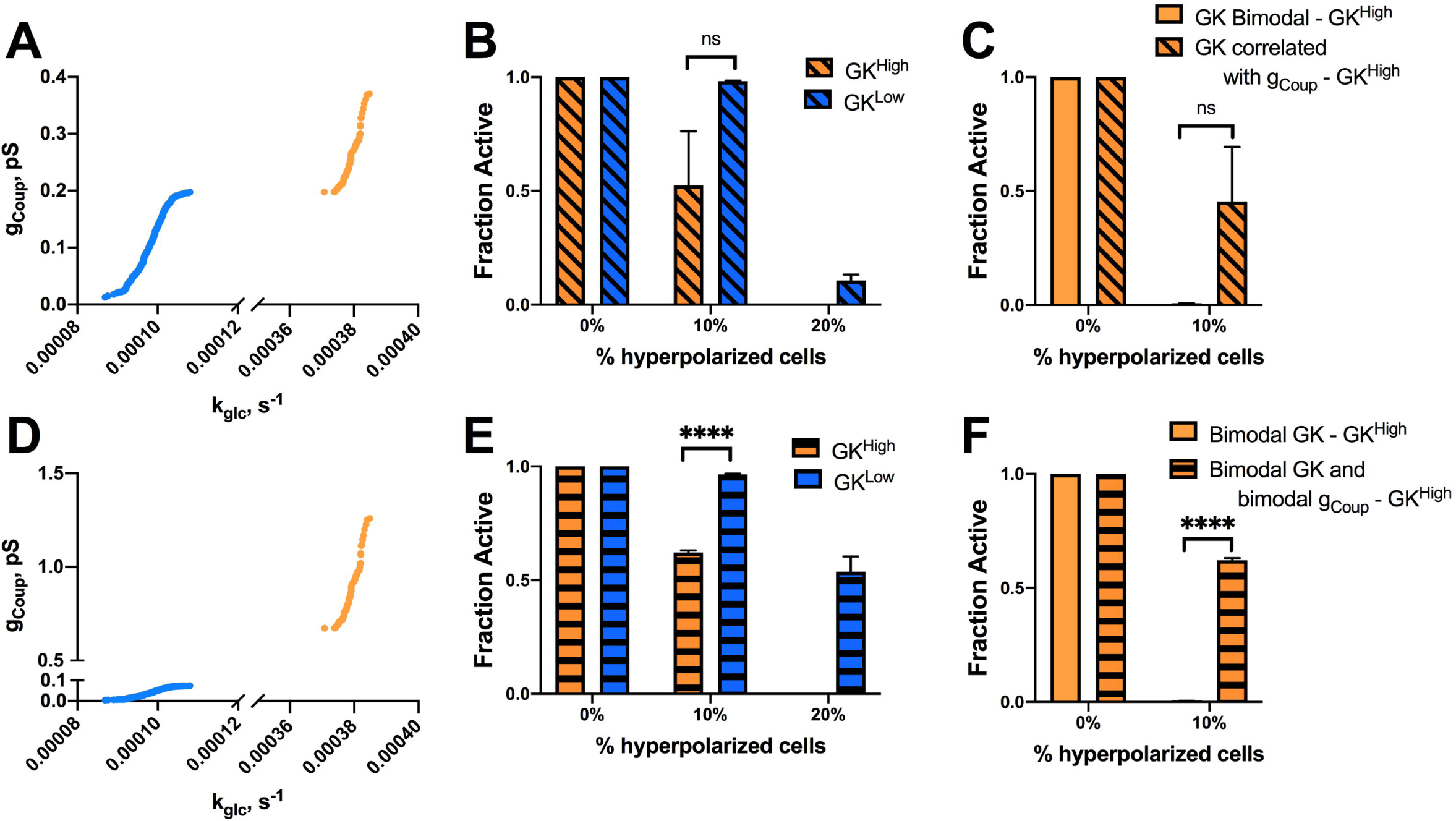
Simulations predicting how changes in coupling impact highly metabolic populations in a bimodal model. A). Scatterplot of g_Coup_ vs. k_glc_ for each cell from a representative simulation where g_Coup_ is correlated with k_glc_. B). Fraction of cells showing elevated [Ca^2+^] activity (active cells) vs. the percentage of cells hyperpolarized in islet from bimodal simulations in k_glc_ with correlated g_Coup_ and k_glc_ as in A. Hyperpolarized cells are chosen either from GK^High^ (orange bars) or GK^Low^ (blue bars) population. C). As in B. but comparing hyperpolarization in GK^High^ cells in the presence and absence of correlations in g_Coup_. D). as in A but from a simulation where g_Coup_ and k_glc_ are correlated AND both g_Coup_ and k_glc_ are bimodal distributions. E). As in B. but for simulations where g_Coup_ and k_glc_ are correlated AND both g_Coup_ and k_glc_ are bimodal distributions. F). As in C. but comparing hyperpolarization in GK^High^ cells in the presence and absence of a bimodal distribution of g_Coup_. Error bars are mean ± s.e.m. Student’s paired t-test was performed for B and E and a Welches t-test for unequal variances was used for C and F to test for significance. Significance values: ns indicates not significant (p>.05), * indicates significant difference (p<.05), ** indicates significant difference (p<.01), *** indicates significant difference (p<.001), **** indicates significant difference (p<.0001). Data representative of 4 simulations with differing random number seeds.

Given this dependence on gap junction coupling, we examined whether decreases in coupling impacted how metabolically active cells controlled islet function. We performed similar simulations as in Fig 1 and Fig 2 for an islet with reduced average gap junction conductance of 50%. In this context, hyperpolarizing highly metabolic populations (GK^Higher^ or GK^High^) or cells with reduced metabolic activity (GK^Lower^ or GK^Low^) reduced islet [Ca^2+^] to a lesser degree than when gap junction conductance was higher (S3 Fig). This applied to simulated islets with either a continuous distribution in GK activity (S3a,b Fig) or a bimodal distribution of GK activity (S3c,d Fig). In each case, a similar difference in islet [Ca^2+^] resulted from hyperpolarizing highly metabolic or low metabolic cells, albeit with greater numbers of cells needing targeting to suppress [Ca^2+^]. Thus, decreasing gap junction coupling does not enhance the ability of small populations of metabolic active cells to maintain islet [Ca^2+^].

### Cells with [Ca^2+^] oscillations that precede the rest of the islet do not drive islet [Ca^2+^] oscillations

Another subpopulation of β-cells that has been associated with islet function are those cells that show [Ca^2+^] oscillations that precede the rest of the islet [29, 31, 40]. These cells have been suggested to have higher intrinsic oscillation frequency [29, 40], which may lend themselves to act as rhythmic pacemakers to drive [Ca^2+^] oscillations across the islet. We next investigated whether a small subpopulation of these cells is able to drive islet [Ca^2+^] oscillatory dynamics. We simulated an islet with a continuous distribution of heterogeneity, as in Fig 1, and identified cells with [Ca^2+^] oscillations that preceded the rest of the islet (low phase) or cells with [Ca^2+^] oscillations that are delayed with respect to the rest of the islet (high phase) (Fig 4a,b). Cells that preceded the rest of the islet (low phase cells) were temporally separated to a greater degree with respect to the rest of the islet compared to cells that were delayed (high phase cells) (Fig 4c). The top 1% and 10% of low phase cells (earlier [Ca^2+^] oscillations) in the islet had higher intrinsic oscillation frequency – the oscillation frequency if the cell is simulated in isolation – and lower GK activity compared to the rest of the islet (Fig 4d,e). This is consistent with prior experimental measurements that demonstrated lower metabolic activity in cells that show earlier [Ca^2+^] oscillations [29]. Conversely, the top 1% and 10% of high phase = cells (delayed [Ca^2+^] oscillations) had lower intrinsic oscillation frequency and high GK activity (Fig 4d,e).

**Figure 4.**
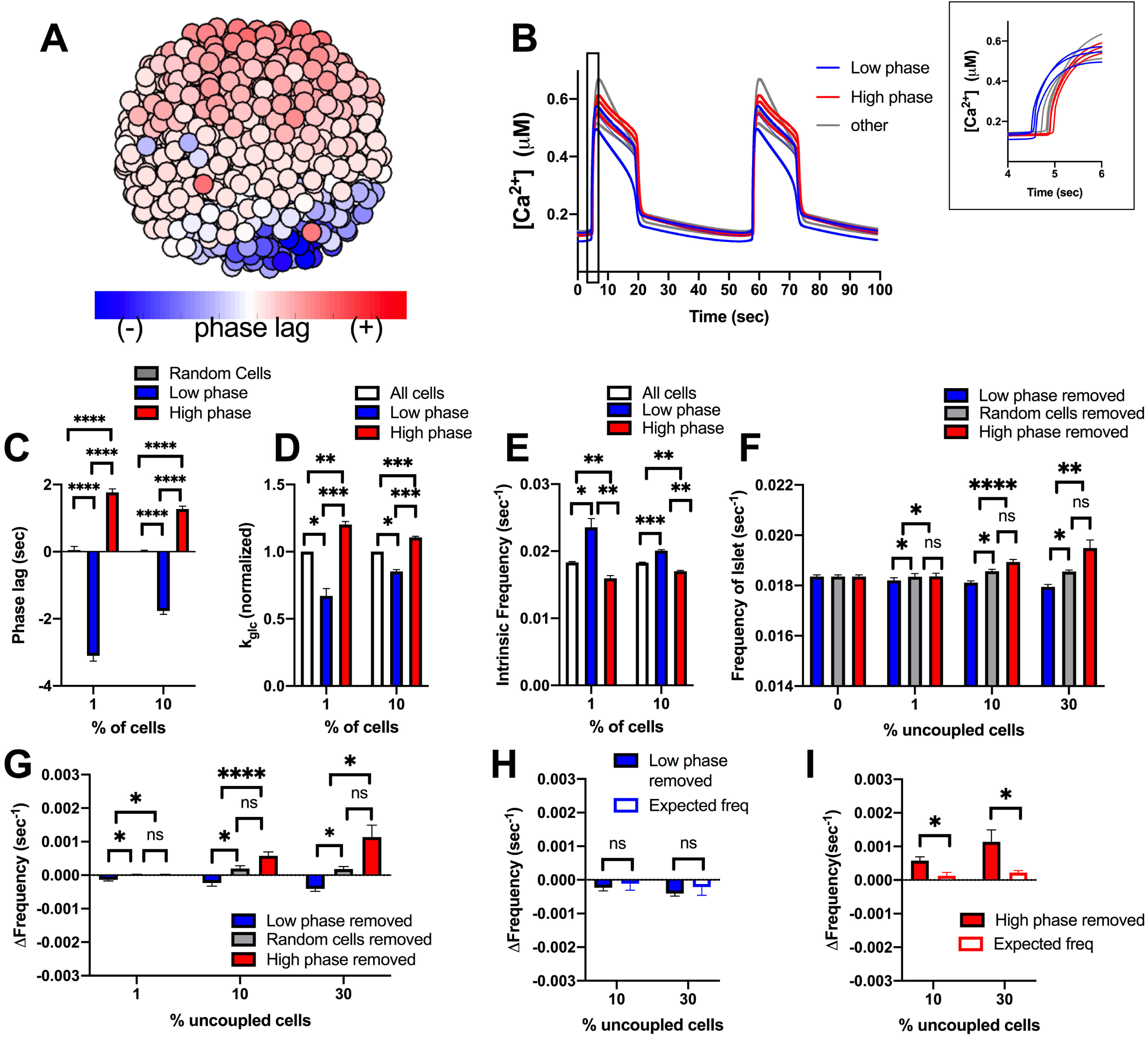
Simulations predicting how small populations of cells contribute to islet frequency. A). Schematic of phase lag across simulated islet with 25% variation in GK activity. B). Representative time courses of [Ca^2+^] for 9 cells in simulated islet at 60pS coupling conductance to determine phase lag of cells in A. Blue traces are low phase cells (negative phase lag), Grey is non low or high phase cells, red is a high phase cells (positive phase lag). Inset: Close up of rise of [Ca2+] oscillation showing phase lags. C). Phase lag from islet average of top 1% or 10% of low phase, high phase cells, or random cells. D). Average k_glc_ from all cells, low phase cells or high phase cells across simulated islet. E). Average intrinsic oscillation frequencies of all cells, top 1% and 10% of low phase cells, or top 1% or 10% of high phase cells when re-simulated in the absence of gap junction coupling (0pS). F). Average frequency of islet when indicated populations of cells are removed from the simulated islet. G). Change in frequency of islet with indicated populations removed with respect to control islet with all cells present. H). Change in frequency when low phase cells are removed compared to average oscillation frequency of remaining cells that indicates the expected oscillation frequency. I). Same as H. but for simulations where high phase cells are removed. Error bars are mean ± s.e.m. Repeated measures one-way ANOVA with Tukey post-hoc analysis was performed for simulations in C-G (if there were any missing values a mixed effects model was used), Student’s paired t-test was performed for H and I to test for significance. Significance values: ns indicates not significant (p>.05), * indicates significant difference (p<.05), ** indicates significant difference (p<.01), *** indicates significant difference (p<.001), **** indicates significant difference (p<.0001). Data representative of 4-9 simulations with differing random number seeds. Random regions were removed for 10% and 30% simulations, but random removal of cells was used for 1% simulations.

To determine the role these cells may play in islet function, we re-simulated the islet with populations of low phase and high phase cells removed from the islet. When populations (1%, 10%, 30%) of low or high phase cells were removed, the elevation of [Ca^2+^] was unchanged (S4a Fig). Similarly, the frequency of the islet did not differ significantly from control islets when up to 10% of low or high phase cells were removed (Fig 4f,g). Low or high phase cells usually exist within a compact region, rather than being distributed randomly across the islet. Removing random cells within a similar sized region impacts frequency of the remaining islet to a lesser degree than removing randomly positioned cells across the islet (S5 Fig). Removal of up to 10% of low phase or high phase cells also showed no change in frequency compared to removal of random cells within a similar sized region (Fig 4f, g). When 30% of low phase cells (earlier [Ca^2+^] oscillations) were removed from the islet, frequency decreased slightly, by ∼2% (Fig 4g). This minor decrease in frequency was equivalent to the average frequency of the remaining cells in the islet, indicating no disproportionate effect of the low phase cells on oscillation frequency (Fig 4h). In contrast, when 30% of high phase cells (delayed [Ca^2+^] oscillations) were removed, the islet frequency increased, by ∼8% (Fig 4g). This increase in frequency upon removing the phase high cells was significantly greater than the average frequency of the remaining cells in the islet, indicating a disproportionate effect of high phase (delayed) cells on oscillation frequency (Fig 4i). When these manipulations were performed in the presence of reduced (50%) gap junction conductance, the changes in frequency were exacerbated: no change in frequency when removing low phase (early) cells and a greater increase in frequency (∼15%) when removing high phase (delayed) cells (S6 Fig). Thus, low phase cells that show earlier [Ca^2+^] oscillations do not drive the [Ca^2+^] oscillation frequency of the islet, when considering a continuous distribution of cell heterogeneity. However, unexpectedly, high phase cells that show delayed [Ca^2+^] oscillations appear to drive a slower [Ca^2+^] oscillation frequency; but only in proportions of at least 30% of the islet.

Low phase and high phase cells that show different timings in their [Ca^2+^] oscillations on average have higher or lower intrinsic [Ca^2+^] oscillation frequency respectively. However, other factors such as gap junction coupling or position within the cluster may also determine their relative timing. We next examined the role of cells that intrinsically have the highest or lowest [Ca^2+^] oscillation frequency (Fig 5a-c). The top 1% or 10% of cells with highest or lowest intrinsic oscillation frequency, showed a frequency substantially different than the islet average (Fig 5d). On average, cells with a higher intrinsic oscillation frequency showed earlier [Ca^2+^] oscillations compared with the average of the islet (Fig 5e) and had lower metabolic activity (Fig 5f). In contrast, cells with the lowest frequency showed delayed [Ca^2+^] oscillations compared with the average of the islet and had higher metabolic activity (Fig 5e,f). This is consistent with previous experimental measurements that demonstrated a negative correlation between oscillation frequency and metabolic activity [29]. We do note that a small (∼0.5%) of cells with low metabolic activity lacked [Ca^2+^] elevations and were excluded from frequency measurements.

**Figure 5.**
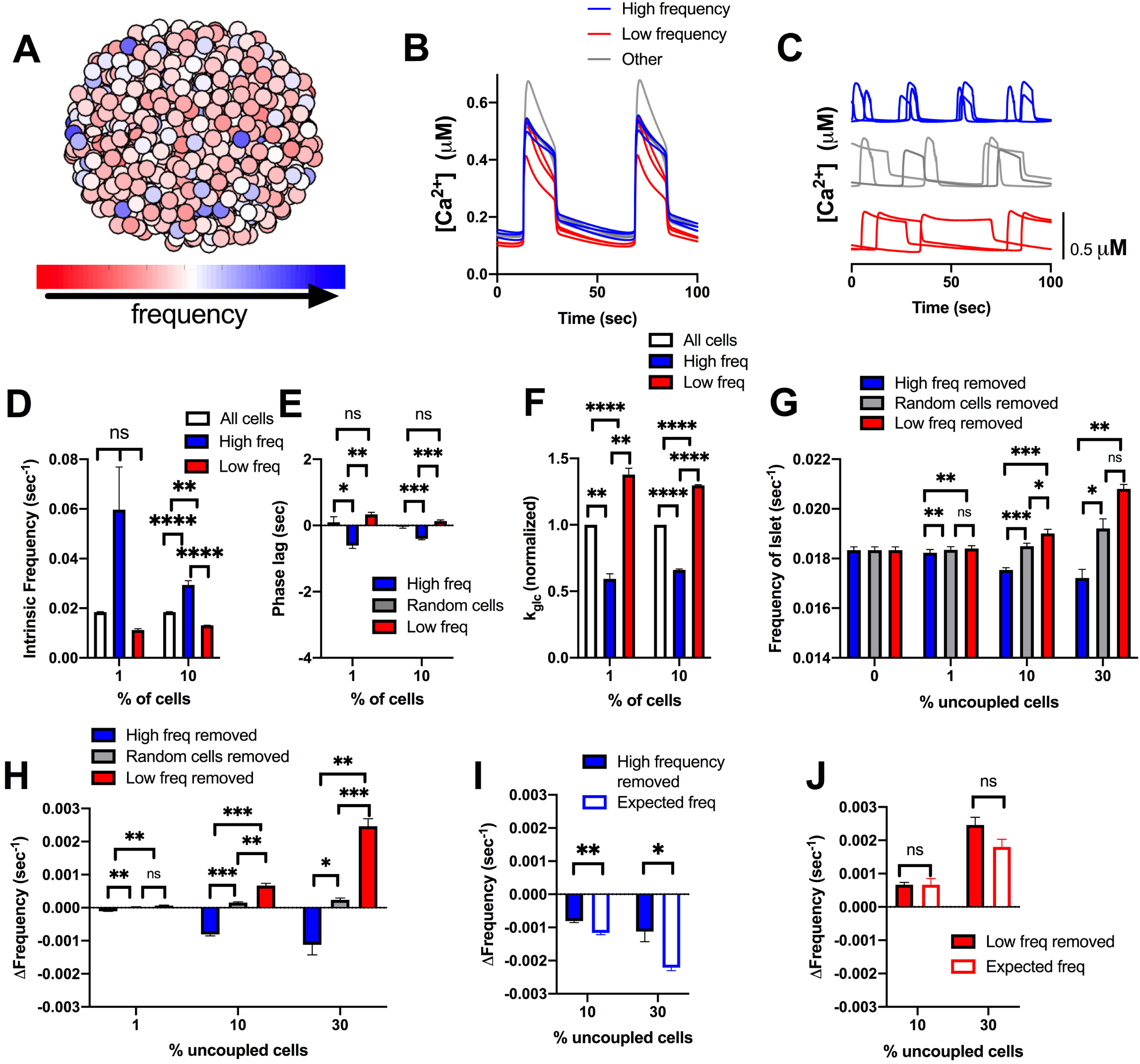
Simulations predicting how intrinsic frequency of cells contributes to islet frequency. A). Schematic of frequency across simulated islet with 25% variation in GK activity. B). Representative time courses of [Ca^2+^] for 9 cells in simulated islet in A in a simulation with full (120pS) coupling conductance. Blue traces are high frequency cells, Grey are cells with frequency near average frequency, red traces are low frequency cells. C). Same cells as in B but showing [Ca^2+^] time courses from an uncoupled simulation (0pS coupling conductance). D). Average intrinsic oscillation frequencies of all cells, top 1% or 10% of high frequency cells, or low frequency cells when re-simulated in the absence of gap junction coupling. E). Phase lag from islet average of top 1% or 10% of low phase, high phase cells, or random cells. F). Average k_glc_ from all cells, high frequency cells, or low frequency cells across simulated islet. G). Average frequency of islet when indicated populations of cells are removed from the simulated islet. H). Change in frequency of islet with indicated populations removed with respect to control islet with all cells present. I). Change in frequency when high frequency cells are removed compared to average oscillation frequency of remaining cells that indicates the expected oscillation frequency. J). Same as I. but for simulations where low frequency cells are removed. Error bars are mean ± s.e.m. Repeated measures one-way ANOVA with Tukey post-hoc analysis was performed for simulations in D-H and a Student’s paired t-test was performed for I and J to test for significance. Significance values: ns indicates not significant (p>.05), * indicates significant difference (p<.05), ** indicates significant difference (p<.01), *** indicates significant difference (p<.001), **** indicates significant difference (p<.0001). Data representative of 5 simulations with differing random number seeds. Random removal of cells across the islet was used for all simulations where random cells removed is indicated.

When greater than 10% or 30% of high frequency cells were removed from the islet, the frequency of the islet decreased, whereas when 10% or 30% of lower frequency cells were removed from the islet, the frequency of the islet increased (Fig 5g,h). However, in each case the change in frequency upon removing high or low frequency cells was not significantly greater than the change when considering the average frequency of the remaining cells (Fig 5i,j). In fact, the decrease in frequency upon removing high frequency cells was significantly less than that considering the frequency of remaining cells (Fig 5i). In each case, the elevation of [Ca^2+^] was unchanged (S4b Fig). These results again suggest that small numbers of cells with differing oscillation frequency do not disproportionately affect islet [Ca^2+^] oscillations.

### A bimodal distribution in frequency lessens the effect of high phase cells

Earlier we considered a bimodal distribution in metabolic activity that better described experimental data (Fig 2)[30]. We next investigated whether high phase and low phase cells may influence the islet to a greater degree when described by a bimodal distribution. From the continuous distribution we previously modelled (Fig 4), we generated a population of cells that incorporated the average properties of low phase cells that showed earlier oscillations in [Ca^2+^] (see methods). This population (10%), which showed a faster oscillation frequency (Fig 6a and S7a Fig) was combined with a population of cells that were similar to the average properties of an islet. The resultant simulated islet showed cells with earlier and delayed [Ca^2+^] oscillations, as before (Fig 6b,c), albeit with a slight reduction in the time between the early and delayed oscillations (Fig 6d). On average, the low phase cells that showed earlier [Ca^2+^] oscillations had higher intrinsic oscillation frequencies (Fig 6e) and lower metabolic activity (Fig 6f), as before. However, the difference between low and high phase cells was not a large as with the continuous distribution. When low phase cells or high phase cells were removed from the islet, the frequency was not significantly different than when random cells were removed (Fig 6g). However, when 10% of low phase cells were removed, the change in frequency was significantly different, albeit small, compared to the expected frequency of the remaining cells in the distribution (Fig 6h). On the other hand, the removal of high phase cells was not significantly different than the expected frequency of the remaining cells (Fig 6i).

**Figure 6.**
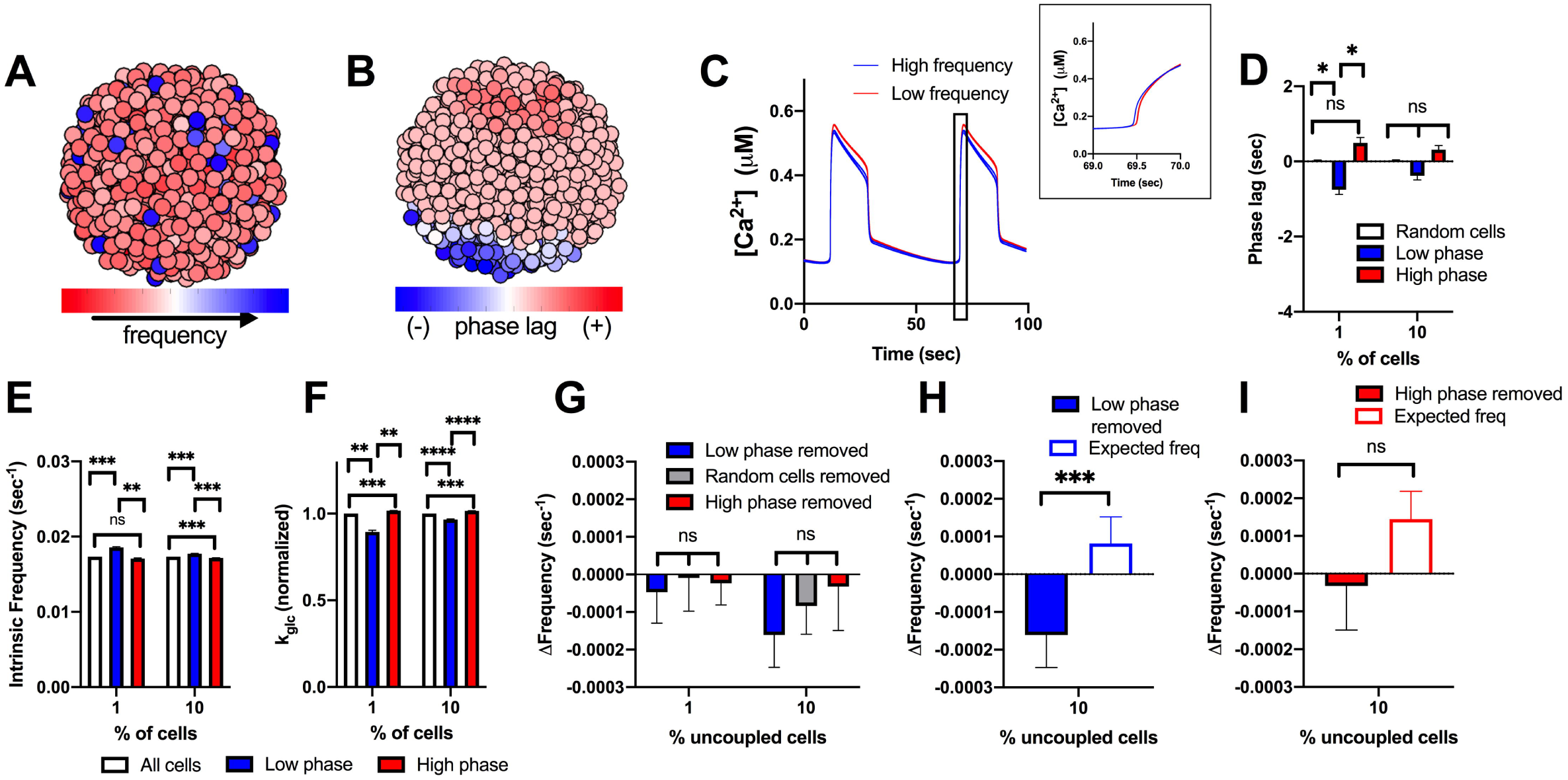
Simulations predicting how cells from a bimodal distribution characterized by low phase cells contribute to islet frequency. A). Schematic of frequency across simulated islet with a bimodal distribution in GK activity. B). Schematic of phase lag across simulated islet with a bimodal distribution in GK activity. C). Representative time courses of [Ca^2+^] for 6 cells in simulated islet in A (and B) in a simulation with full (120pS) coupling conductance. Blue traces are high frequency cells, red traces are low frequency cells. Inset: Close up of rise of [Ca2+] oscillation showing phase lags. D). Phase lag from islet average of top 1% or 10% of low phase, high phase cells, or random cells. E). Average intrinsic oscillation frequencies of all cells and 1% or 10% of low phase cells, or 1% or 10% of high phase cells when re-simulated in the absence of gap junction coupling (0pS). F). Average k_glc_ from all cells and top 1% or 10% of low phase cells or high phase cells across simulated islet. G). Change in frequency of islet with indicated populations removed with respect to control islet with all cells present. H). Change in frequency when low phase cells are removed compared to average oscillation frequency of remaining cells that indicates the expected oscillation frequency. I). Same as H. but for simulations where high phase cells are removed. Repeated measures one-way ANOVA with Tukey post-hoc analysis was performed for simulations in D-G and a Student’s paired t-test was performed for H and I to test for significance. Error bars are mean ± s.e.m. Significance values: ns indicates not significant (p>.05), * indicates significant difference (p<.05), ** indicates significant difference (p<.01), *** indicates significant difference (p<.001), **** indicates significant difference (p<.0001). Data representative of 5 simulations with differing random number seeds. Random removal of cells across the islet was used where random cells removed is indicated.

We further examined how the islet behaved when the high frequency population of cells were removed. These high frequency cells showed only slightly earlier [Ca^2+^] oscillations compared to the rest of the islet on average (S7b Fig) but did show lower metabolic activity (S7c Fig). Upon removal of these high frequency cells, the islet showed significantly slower oscillations (S7d Fig), that were slower than expected given the average frequency of the remaining cells (S7d,e Fig). However, the change in frequency was still low (∼2%). When these high frequency cells were positioned with the same spatial distribution as low phase cells, the change in frequency upon their removal was significantly greater but was still relatively small and similar to the change seen when high frequency cells were removed from the continuous distribution model (∼5%) (S8 Fig). In conclusion, within a bimodal distribution, a small population of cells with higher frequencies does not substantially impact the frequency of the islet.

### Limited excitatory gap junction current can explain lack of action of small sub-populations

To understand the basis by which cells with differing metabolic activity and oscillatory frequency interact, we examined the gap junction currents for cell populations within the islet (Fig 7a). As expected, the total membrane current was highest in magnitude during the upstroke and downstroke of the [Ca^2+^] oscillation, and low in magnitude during the active and silent phase (Fig 7b-d). Conversely, the gap junction current was highest during the active and silent phase of the [Ca^2+^] oscillation but was minimal during the upstroke and downstroke of [Ca^2+^] (Fig 7b-c,e). Thus, there is less communication between cells during the upstroke and downstroke of [Ca^2+^] oscillations compared to the stable active and silent phases.

**Figure 7.**
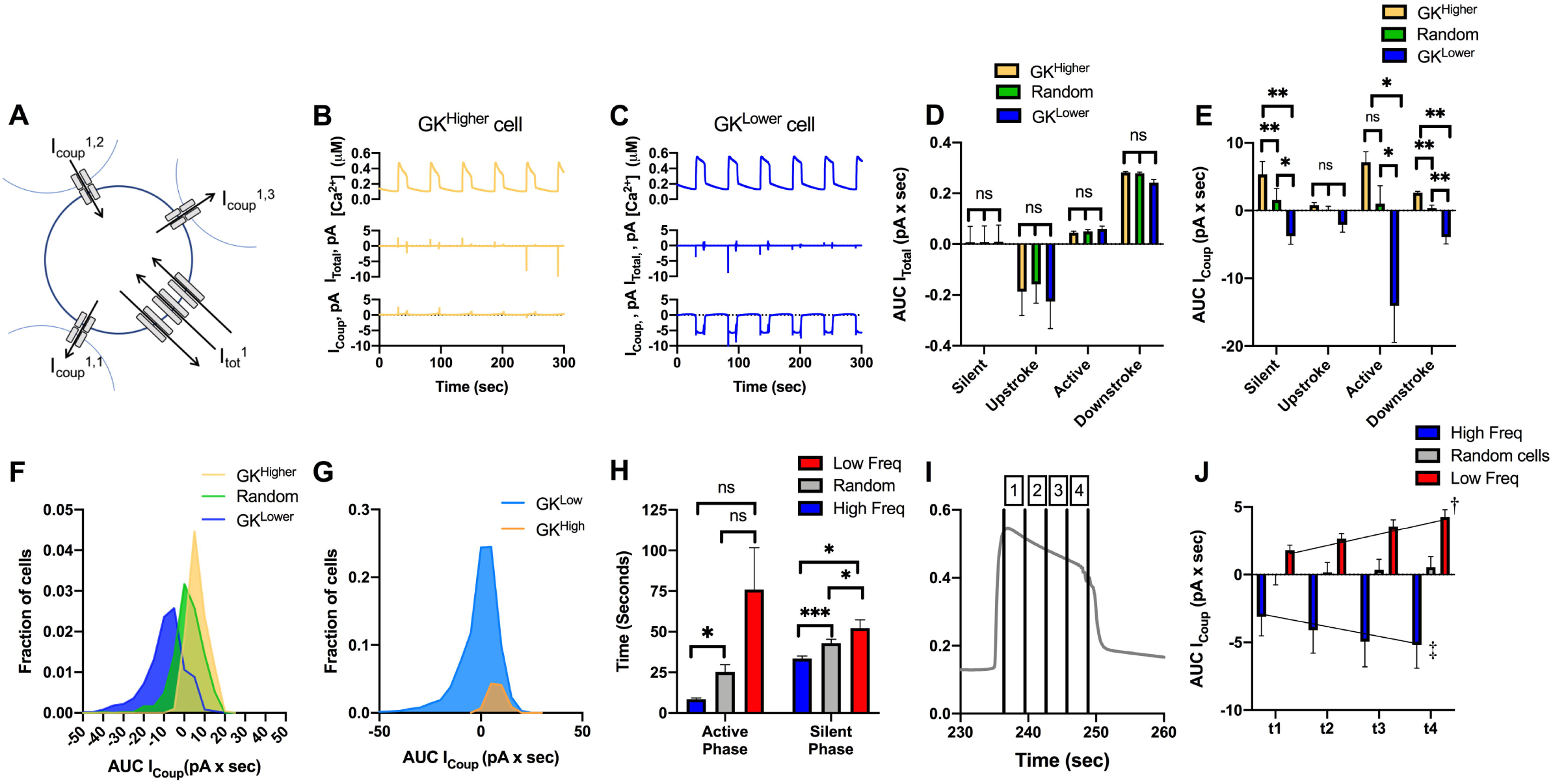
Gap junction current in cells with high/low metabolic activity or oscillation frequency. A). Schematic of cell within the simulated islet, showing 3 gap junction currents that contribute to the total gap junction current, together with the total membrane current. B). Time course of [Ca^2+^] from a cell, together with the total membrane current and total gap junction current for a representative cell with higher metabolic activity (k_glc_). C). As in B for a representative cell with lower metabolic activity. D). Total membrane current, as expressed by an area under the curve (AUC), for each phase of the [Ca^2+^] oscillation averaged over the 10% of cells with highest or lowest k_glc_ or a random 10% of cells. E). As in D for total gap junction current. F). Distribution of total gap junction current, as expressed by AUC, for the 10% of cells with highest or lowest k_glc_ or a random 10% of cells. G). As in E for a bimodal distribution in k_glc_. H). Mean duration of active phase and silent phase averaged over the 10% of cells with highest or lowest oscillation frequency, or a random 10% of cells. I). Mean islet [Ca^2+^] time course showing different portions of the active phase (1-4). J). Mean islet gap junction current during different portions of the active phase, as indicated in I for the 10% of cells with highest or lowest oscillation frequency, or a random 10% of cells. Black lines are fitted regression lines. Error bars are mean ± s.e.m. Repeated measures one-way ANOVA was performed for data in D, E, H to test for significance. Linear regression was performed on data in J. Significance values: ns indicates not significant (p>.05), * indicates significant difference (p<.05), ** indicates significant difference (p<.01), *** indicates significant difference (p<.001), **** indicates significant difference (p<.0001), † indicates significant linear regression (p<.05), ‡ indicated significant linear regression (p<.01). Data representative of 5 simulations with differing random number seeds.

The total membrane current did not differ significantly between cells with high or low metabolic activity (Fig 7d). However, there was a substantial difference in gap junction current between cells with high or low metabolic activity (Fig 7e). Cells with high metabolic activity showed a positive (outward, hyperpolarizing) gap junction current, whereas cells with low metabolic activity showed a negative (inward, depolarizing) gap junction current, across all phases of the [Ca^2+^] oscillation (Fig 7e). The magnitude of the gap junction current for less metabolically active cells was also greater. This larger gap junction-mediated current would be expected to hyperpolarize neighboring cells to a greater degree than metabolically active cells depolarizing neighboring cells. Nevertheless, there was significant variability, such that some cells with low metabolic activity had little gap junction current and some cells with high metabolic activity had a positive current that would depolarize neighbors (Fig 7f). When examining the bimodal simulation (Fig 2), we observed broadly similar findings where cells with high metabolic activity depolarize their neighbors whereas cells with low metabolic activity hyperpolarize their neighbors (Fig 7g).

Finally, given the stronger gap junction current associated with the active and silent phases, we analyzed the relationship between the duration of these phases for cells with high and low frequency. Cells with a higher intrinsic oscillation frequency showed both a shorter active phase and shorter silent phase compared to cells with a slower intrinsic oscillation frequency, with there being a greater difference in the active phase (Fig 7h). Interestingly, the whole islet active and silent phase times were similar to those of cells with a higher oscillation frequency (which on average have lower metabolic activity). During the active phase, the gap junction current was lowest at the beginning of the active phase and greatest just before the downstroke (Fig 7i,j). We measured changes to the duration of the active and silent phases after removal of low/high phase cells and low/high frequency cells from Fig 4 and 5. When either high phase cells or low frequency cells were removed, the active phase and duty cycle duration decreased compared to when either low phase cells or high frequency cells were removed, respectively (S9 Fig). Thus gap junction coupling contributes more to sustaining the active phase compared to initiating the active phase. While slower oscillating cells contribute significantly to setting the islet frequency, given the greater gap junction current, faster oscillating cells may limit the duration of the active phase by terminating the oscillation.

## Discussion

β-cell heterogeneity has largely been studied in single cells. However, recent studies have demonstrated that heterogeneity plays a physiological role in regulating insulin release within the islet [29, 30, 35]. Previously, using computational models and experimental systems, we demonstrated that a large minority (close to 50%) of metabolically active β-cells were necessary to maintain the activity of the islet [35]. In contrast to this, experimental and theoretical studies have suggested that small (∼10%) highly functional subpopulations may be required to maintain whole islet electrical dynamics [30, 41]. Here, we investigated the theoretical basis by which small populations of cells may impact islet electrical dynamics.

### Small populations of metabolically active cells are not required to drive elevations in [Ca^2+^]

To determine whether small populations of metabolically active β-cells could drive elevations in [Ca^2+^], we constructed two types of islet simulations: showing either a continuous distribution in metabolic activity or a bimodal distribution in metabolic activity. In each case, we either hyperpolarized the most metabolically active cells or removed them from the simulation. These manipulations are equivalent to those applied in the literature. For example, one study used optical stimulation of eNpHr3.0 to induce a hyperpolarizing Cl^-^ current in 1-10% of cells that showed high levels of [Ca^2+^] coordination and elevated GK [30]. Another study used optical stimulation of ChR2 to induce a depolarizing cation current, with the ∼10% of cells activating large parts of the islet showing higher NAD(P)H [29]. In our simulations, we found hyperpolarizing those cells with increased metabolic activity generated similar findings: hyperpolarizing more metabolically active cells silenced the islet to a much greater degree than hyperpolarizing less metabolically active cells. Thus, hyperpolarization or depolarization of metabolically active β-cells can disproportionately suppress or activate islet function, via gap junction coupling.

Importantly, the effects of this targeted silencing were found for both a broad continuous distribution (Fig 1) and for a bimodal distribution (Fig 2). Within the literature there is not exact consistency in the level of metabolic heterogeneity present. Within dissociated β-cells, a variation of 20-30% in NAD(P)H responses has been observed experimentally [19, 37], and in intact islets a variation of 10-20% has been observed [37]. Instead, ∼50% variation is needed to describe experimental observations here. However, early analysis of GK heterogeneity via immunohistochemistry observed substantial variations, which while not quantified would be equivalent to >50% [17]. Similarly, in isolated β-cells the glucose threshold for elevated NAD(P)H varies by ∼50% (3-10mM) [16, 42]. This latter study also found a non-normal distribution with ∼20% of highly metabolically active β-cells. Thus, the distributions required in our model to generate results equivalent to experimental observations are broadly feasible. Furthermore, we do note the process of removing β-cells from the islet via dissociation causes cell stress and could disrupt metabolic signatures. Highly metabolically active cells may also be more susceptible to environmental stress [30, 43]. Therefore, further analysis, in situ, is needed to precisely quantify the level of heterogeneity present.

Interestingly, we observed very different results when comparing the effect of targeted hyperpolarization of a set of cells and targeted removal of a set of cells. Hyperpolarizing a small population of metabolically active cells silenced the islet, whereas removal of this same cell population had little impact. Upon removal, we did observe a small reduction in the duty cycle (% of time the islet resides in the active phase) of ∼10%. Duty cycle is a large determinant of insulin release, thus a ∼10% reduction would not be expected to impact insulin release substantially. However, experimental measurements would be needed to exclude whether exocytosis varies across the pulse duration. As such, the manipulations involving hyperpolarization and cell removal, theoretically, assess the importance a cell has on islet function in different ways. Thus, care must be taken when interpreting the results of optogenetic stimulation-based analysis.

Cell removal from the simulation may be considered similar to the experimental ablation of that cell. Ablation of small populations of cells that show earlier [Ca^2+^] oscillations (see below), but which overlaps with those cells that show increased [Ca^2+^] coordination, has experimentally been demonstrated to reduce the elevation in [Ca^2+^] across zebrafish islets. These studies showed a substantial reduction in [Ca^2+^] amplitude, whereas our theoretical findings showed no apparent differences in amount of active cells. Little change in [Ca^2+^] activity is observed in the model when removing either those cells with earlier Ca^2+^ oscillations (S4 Fig) or those cells with elevated metabolic activity that when hyperpolarized silences islet [Ca^2+^] (Fig 1,2). However, differences do exist between zebrafish islets and mouse islets which our model is based upon and has been validated against, including islet size, gap junction protein isoform and Ca^2+^ dynamics [44, 45]. Thus, species differences may account for these observations.

The way cells interact within our simulated islet is restricted to gap junction electrical coupling. As such, we conclude that gap junction communication is unlikely to be able to explain the role small cell subpopulations play in islet function, under the model assumptions presented here. These conclusions are also consistent with elevated oscillatory [Ca^2+^] being maintained upon a loss of Cx36 gap junction coupling [15], albeit with a lack of synchronization. However, we do note that first phase insulin release is diminished upon a loss of Cx36 gap junction coupling [46]. Therefore, we cannot exclude that small cell subpopulations can drive [Ca^2+^] elevations via gap junction coupling during the initial first phase response.

β-cells can communicate across the islet via paracrine communication. This includes inhibitory factors such as GABA, 5-HT, dopamine and Ucn3 (via δ-cell somatostatin release) and stimulatory factors such as ATP [47-49]. Thus, it is conceivable, small sub-populations of metabolic active cells are secreting increased levels of stimulatory paracrine factors. Alternatively, small sub-populations may be acting via other endocrine cells, such as glucagon-secreting α-cells, to stimulate other β-cells within the islet [50]. Removal of immature cell populations can also disrupt islet function, suggesting a broader remodeling of the islet can be induced by small cell sub-populations [51]. Therefore, analyzing whether subpopulations show differential release of paracrine factors will be important to better elucidate their function within the islet.

### Gap junction coupling homogenizes subpopulations and reduces their impact

Gap junction coupling allows for heterogeneous populations of β-cells to act in a cohesive manner. For example, when populations of normally excitable and inexcitable cells combine within an islet, gap junction coupling ensures that a uniform response occurs, whether this be suppressed [Ca^2+^] or coordinated elevated [Ca^2+^] [34]. Some cell populations have been suggested to have elevated connectivity with other cells in the islet, as measured by correlated [Ca^2+^] oscillations [30, 31], and could result from an increase in gap junction coupling.

When more metabolically active cells had increased coupling conductance, hyperpolarizing those cells had less impact on suppressing islet function (Fig 3). If gap junction coupling is elevated in metabolically active cells, it is reduced in less metabolically active cells. A decrease in coupling lessens how the islet is suppressed in the presence of inexcitable cells that transmit hyperpolarizing current across the islet. Thus, hyperpolarizing a population of metabolically active cells would transmit less hyperpolarizing current beyond the nearest neighbor cells.

We also observed that less metabolically active cells show a greater gap junction current that hyperpolarizes neighboring cells. Thus, there is an asymmetry by which metabolically active and inactive cells within the islet act (Fig 7). As such, increases in coupling are not beneficial for highly metabolic cells to control the islet. Rather distributing the coupling more uniformly allows all cells within the islet to coordinate their activity.

### Small subpopulations cannot efficiently act as rhythmic pacemakers

Multiple studies have identified cells that consistently show earlier [Ca^2+^] oscillations that may drive the dynamics of [Ca^2+^] across the islet [29, 31, 40]. These populations have been suggested to have a higher intrinsic oscillation frequency and thus act as a rhythmic pacemaker [29], in the same manner as the cardiac SA node. Here, we investigated whether a small subpopulation of cells with increased oscillation frequency could act as such a pacemaker. We found that cells that show earlier [Ca^2+^] oscillations do have a higher intrinsic oscillation frequency. However, upon removal of these cells, the islet [Ca^2+^] oscillations changed little, suggesting that small populations of these cells are unable to pace islet [Ca^2+^] oscillations. This initially is surprising as with all cells capable of firing, the cell with the highest frequency will depolarize first and stimulate neighbors to fire. However, at least ∼30% of high frequency cells are required to even slightly impact islet oscillation frequency. These findings are consistent with prior modelling studies where cells with fast and slow oscillation frequencies, when combined within an islet, led to an overall oscillation midway between the intrinsic cell oscillations [52]. This suggests the oscillation frequency is not per se determined by a small pacemaker population but rather is formed by a weighted combination of all cells across the islet. Thus, the islet also shows significant redundancy where only loss of large populations of cells impacts the activity or dynamics of [Ca^2+^] (Fig 8). Further, the introduction of a small population (∼10% cells) with a defined high intrinsic oscillation frequency has little impact on islet [Ca^2+^] oscillations frequency and wave propagation.

**Figure 8.**
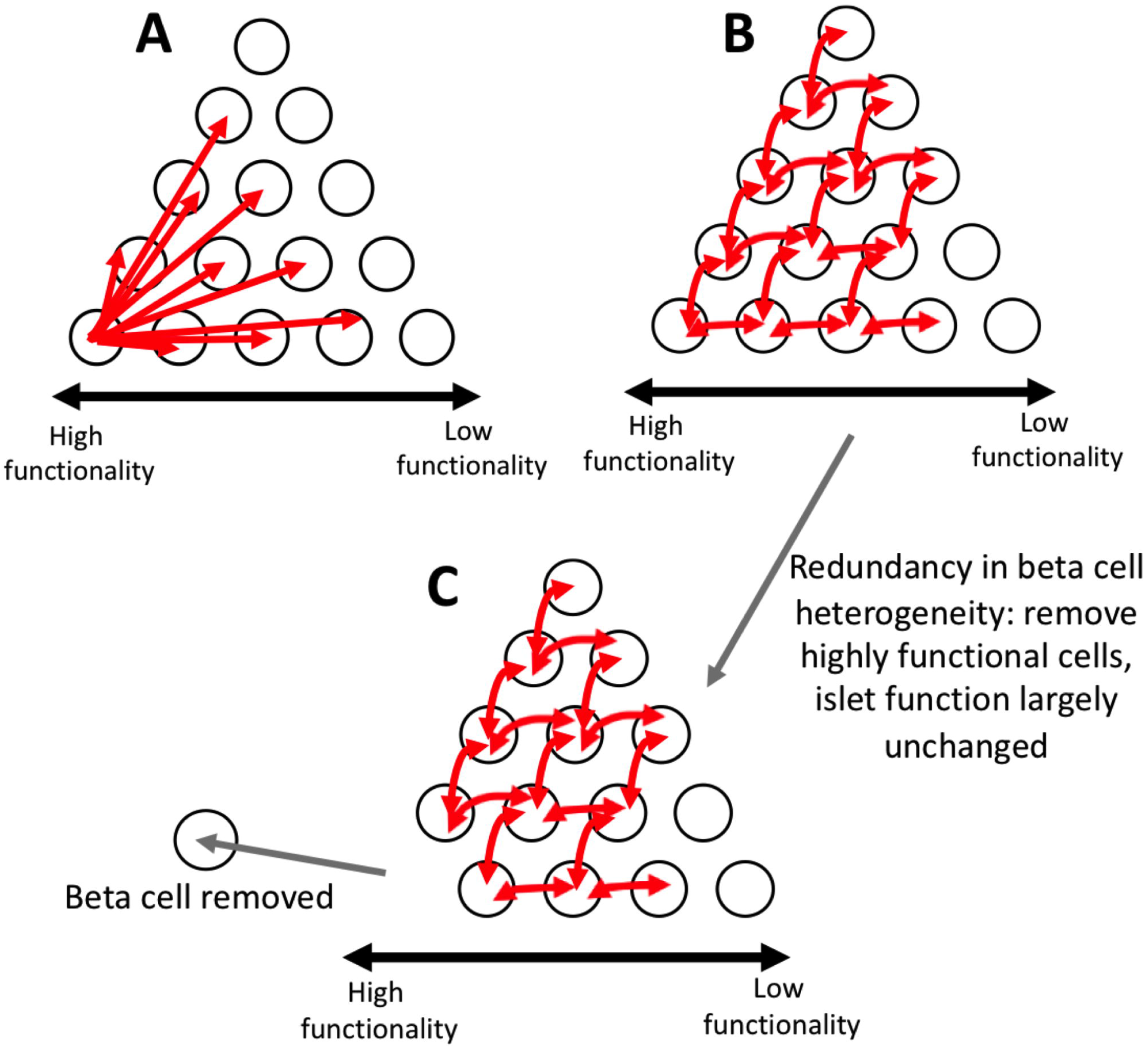
Schematic of multicellular dynamics of the islet. A). Schematic of suggestion that small subpopulations of highly functional cells can control whole islet dynamics. White circles represent β-cells. Red arrows represent which cells can be controlled by individual cell where the arrow begins. Increasing functionality in cells is from right to left B). Same as A, but a schematic of how our simulations predict islet [Ca^2+^] dynamics are controlled. Our simulations predict that control is redundant, and many cells can control many other cells. Our simulations predict there is not one small subpopulation that controls the entire islet. C). Same as B, but schematic of how our simulations predict the islet responds when highly functional subpopulations are removed. When highly functional subpopulations are removed, the remaining cells are able to maintain the function of the islet due to the redundancy in control.

In contrast to removal of cells that show earlier [Ca^2+^] oscillations, removal of those cells that show delayed [Ca^2+^] oscillations increased the frequency of islet [Ca^2+^] oscillations (Fig 4). These cells on average showed slower oscillations. Therefore, slow [Ca^2+^] oscillations contribute to setting the islet [Ca^2+^] oscillation frequency to a greater degree. This also suggests that slow metabolic oscillations will better coordinate [Ca^2+^] dynamics across the islet, rather than purely a faster-oscillating electrical subsystem. Nevertheless, at least 30% of these slow oscillators are needed to have a substantial impact on the islet dynamics, which is consistent with the oscillation frequency again being formed by a weighted combination of all cells across the islet.

We did not observe a complete overlap between cells that show earlier/delayed [Ca^2+^] oscillations and cells with faster/slower intrinsic [Ca^2+^] oscillations, respectively. Similarly, while removal of the highest and lowest frequency cells changes the overall islet frequency to a greater degree, only removal of cells with delayed [Ca^2+^] oscillations showed a change in frequency above that expected, given the frequency of the remaining cells. Thus, other properties of the islet also contribute to setting the islet oscillation frequency, and determining these properties remains to be determined.

Therefore, our simulations indicate that there is not a small population of rhythmic pacemaker cells within the islet. Rather, a large number of cells are needed to impact islet frequency. Of interest, the distribution of cells with faster or slower intrinsic [Ca^2+^] oscillations in our simulation is distributed across the islet, whereas cells that show earlier or delayed [Ca^2+^] oscillations exist within a specific region within the islet, often at the islet edge. While having only a minor impact, the spatial distribution of higher frequency cells was important in affecting islet [Ca^2+^] oscillations. Whether intrinsically fast or slow oscillating cells show some spatially restricted distribution is unknown. A different spatial organization could potentially contribute to a greater control over islet frequency, especially if slow oscillators overlap with other properties of the islet that confer greater control over islet oscillation frequency.

We also speculate that the level of gap junction coupling for cells with slower or faster oscillations may be important: the time course of gap junction current indicates that faster oscillating cells transmit a greater hyperpolarizing current to neighboring cells earlier, as compared to slower oscillating cells. This may explain why the islet oscillation active phase duration trends closer to those cells with a higher frequency and thus shorter active phase duration. However, given the lower gap junction current in the silent phase, this appears not to be sufficient to disproportionately impact the oscillation frequency.

## Summary

Overall, the results from this study show how small populations of highly functional cells impact islet function via gap junction electrical coupling. Our simulations suggest that both a small subpopulation of metabolically active cells or the most metabolically active subset of cells within a continuous distribution are unable to maintain elevated [Ca^2+^] across the islet via gap junction coupling. Further, a small population or subset of cells that shows early [Ca^2+^] elevations or that have a higher oscillation frequency are also unable to act as rhythmic pacemakers to drive oscillatory [Ca^2+^] dynamics. As such the mechanism(s) by which these cells may act to impact islet function should be further investigated.

## Methods

### Coupled β-cell electrical activity model

The coupled β-cell model was described previously [35] and adapted from the published Cha-Noma single cell model [53, 54]. All code was written in C++ and run on the SUMMIT supercomputer (University of Colorado Boulder). Model code is included in supplemental information (Files S1). All simulations are run at 8mM glucose unless otherwise noted.

The membrane potential (V_i_) for each β-cell i is related to the sum of individual ion currents as described by [53]:

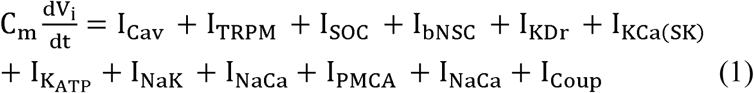

Where the gap junction mediated current I_Coup_ [34] is:

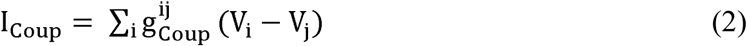

### Modelling GK activity

The flux of glycolysis J_g1c_, which is limited by the rate of GK activity in the β-cell, is described as:

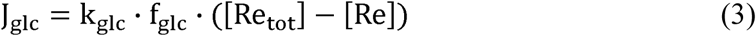

Where k_g1c_ is the maximum rate of glycolysis (equivalent to GK activity), which was simulated as a continuous Gaussian distribution with a mean of 0.000126 ms^-1^ and standard deviation of 25% of the mean (unless indicated). [Re_tot_] = 10mM, the total amount of pyrimidine nucleotides. The ATP and glucose dependence of glycolysis (GK activity) is:

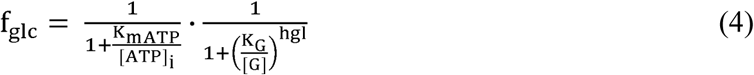

Where [G] is the extracellular concentration of glucose, hgl is the hill coefficient, K_G_ is the half maximal concentration of glucose, and K_mATP_ is the half maximal concentration of ATP.

For simulations with changes in variation in GK, the mean remained the same at 0.000126 ms^-1^, but standard deviation of 1% or 50% of the mean was used.

### Hyperpolarizing cell populations

Hyperpolarization of cells was induced by including a V-independent leak current I_hyper_ that hyperpolarizes the cell [36], described as:

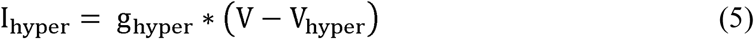

Where g_hyper_ is the hyperpolarizing conductance which is zero in the absence of the applied hyperpolarizing current and is g_hyper_’ (1-p_0KATP_) during applied hyperpolarization. The number of cells that were hyperpolarized were defined as the fraction P_hyp_ multiped by the number of cells, N (1000 in all simulations).

### For bimodal distribution of GK

The bimodal distribution of GK can be described by the following 2 equations.

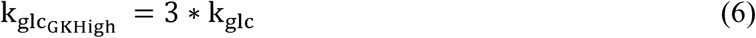

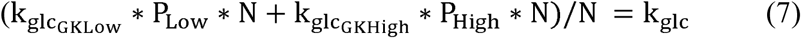

Where 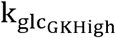 is the mean rate of glycolysis for the GK_High_ population and is 3 times the islet mean, k_g1c_. The mean rate of glycolysis for the GK_Low_ population, 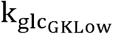, is scaled so that the islet mean, k_g1c_ of the whole simulated islet remains the same at 0.000126 ms^-1^ (7). P_Low_, the percent of GK_Low_ cells in the simulation, is 90%, and P_High_, the percent of GK_High_ cells, is 10% in all bimodal simulations. N is the number of cells in the simulation (1000). The standard deviation of 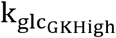 and 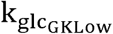 is 1% of the mean for bimodal simulations.

### Modelling changes in coupling

Heterogeneity in Cx36 gap junctions is modeled as a γ-distribution with parameters k=θ=4 as described previously [29] and scaled to an average g_Coup_ between cells = 120pS.

Simulations where k_glc_ and g_Coup_ are correlated, the values for k_glc_ and g_Coup_ for the 1000 cells are randomly calculated, then these values are ordered for both k_glc_ and g_Coup_ and paired together so that the highest k_glc_ and highest g_Coup_ are pairs. The paired k_glc_ and g_Coup_ values are then randomly distributed to the cells in the simulation.

For simulations where g_Coup_ is described as a bimodal distribution, the GK_High_ cells are given 3x the average coupling and then the distribution is scaled so that mean g_Coup_ remains 120pS similarly to equations (6) and (7).

For simulations where cells are removed, the conductance, g_coup_, of the cells to be removed is set to 0 pS. Removed cells are excluded from subsequent islet analysis.

### Determining high and low phase cells

To determine low phase and high phase cells, one full [Ca^2+^] oscillation is taken between time points 300 sec to 400 sec. This time point ensures the model and frequencies are stable and in the second phase of [Ca^2+^] oscillations. A cross correlation is used to determine the time delay of each cell time course compared to the mean [Ca^2+^] across the islet, using xcorr() in MATLAB. A negative delay is therefore equivalent to an earlier oscillation. The low phase cells are determined as the cells with the most negative time delay and the high phase cells are determined as the cells with the most positive time delay. If the cutoff occurs where multiple cells have the same delay, then a random cell is chosen from the cells with the same lag.

### Modelling bimodal for low phase cells

The mean values of the low phase and non-low phase cells in the continuous distribution was used to define a new bimodal distribution as described in Table S1. All standard deviations are 1% of the mean. For more information on parameters see [55]. The ‘low phase’ cell population, N_lowphase_ comprised 10% of the islet (left column), and the N_non-low_ comprised the other 90% (right column).

### Simulation data analysis

All simulation data analysis was performed using custom MATLAB scripts. The first 1500 time points (150 sec) were excluded to allow the model to reach a stable state.

<u>Fraction Active</u> was determined by calculating the fraction of cells that were active relative to the total number of simulated cells (1000). Cells were considered active if membrane potential, [Ca^2+^] exceeded 0.165µM at any point in the time course.

<u>Duty Cycle</u> was determined as the fraction of the [Ca^2+^] oscillations spent above a threshold value during the time course analyzed. This threshold value was determined as 50% of the average amplitude of [Ca^2+^] in an islet simulated at 8mM glucose with 25% variation in GK activity or time above 70% of the maximum [Ca^2+^] (S9 Fig). Duty cycle was reported as the mean across all cells in the simulated islet.

<u>Frequency of a cell in the islet</u> was determined by taking the [Ca^2+^] time-course between times 150 sec and 400 sec and identifying the first 2 peaks. The peak to peak time was determined, and this oscillation period was inverted to calculate the frequency. For whole islet frequency calculations, the coupling in the islet is g_Coup_=120pS and the mean islet frequency is calculated over all cells in the simulation.

<u>Intrinsic frequencies</u> were determined using simulations where the mean coupling conductance of all cells in the simulation is g_Coup_=0pS so that all cells oscillate on their own without influence from other cells within the simulation. When determining low and high frequency cells in the simulation, only active cells were used.

<u>Expected Frequency</u> was determined by finding the average intrinsic frequencies of the cells (g_Coup_=0pS) that are included in the simulation. These values are then compared to the simulation where g_Coup_=120pS.

<u>Total gap junction current</u> for a cell was calculated by summing the gap junction current over each connection between the cell and all of its neighbors, as in equation (2). The total membrane current was calculated as the sum over each current for that cell, as in equation (1).

<u>Active, silent, upstroke and downstroke phases</u>were chosen manually. The Area Under the Curve (AUC) was calculated using the trapz() function in MATLAB, which calculates trapezoidal integration over the time period. AUC was calculated for each cell in the given decile and then averaged over those cells.

<u>Active phase duration</u> for one oscillation was determined for each cell, as the total time [Ca^2+^] was above 70% of the maximum value, divided by the number of oscillations over the duration assessed. The silent phase duration was similarly calculated as the total of time [Ca^2+^] was below 40% of maximum value.

### Statistical analysis

All statistical analysis was performed in Prism (GraphPad). Either a Student’s t-test (or Welch’s t-test for significantly difference variances) or a one-way ANOVA with Tukey post-hoc analysis was utilized to test for significant differences for simulation results. Paired t-test or repeated measures ANOVA was used anywhere the results were compared with a simulated matching control islet or groups within the same islet, e.g. before a population was either hyperpolarized or uncoupled. Data is reported as mean ± s.e.m. (standard error in the mean) unless otherwise indicated.

## Acknowledgements

The authors thank Dr David J Hodson (University of Birmingham, UK) and Dr Victoria Salem (Imperial College London, UK) for reviewing this manuscript and for providing helpful comments and suggestions. The authors are also grateful for utilization of the SUMMIT supercomputer from the University of Colorado Boulder Research Computing Group, which is supported by the National Science Foundation (awards ACI-1532235 and ACI-1532236), the University of Colorado Boulder, and Colorado State University.

## Author contributions

Conceptualization and data curation: JMD, RKPB. Funding acquisition, project administration and supervision: RKPB. Formal analysis and investigation: JMD, JKB. Methodology: JMD, JKB, RKPB. Resources, software, validation and visualization: JMD. Writing, review and editing: JMD, RKPB.

## Supporting information captions

**S1 Figure. Histograms of GK activity (k**_**glc**_**) and g**_**Coup**_ **for all continuous and bimodal distributions in GK activity for Fig 1-3**. A). All continuous distributions’ histograms. Left: Average frequency of cells at varying GK rate (k_glc_) for simulations that have different standard deviations in GK activity from Fig 1. Right: Corresponding histogram of average frequency of cells at varying coupling conductance (g_Coup_). B). As in A but for simulations with a bimodal distribution of GK activity from Fig 2. C). As in A for simulations with bimodal distribution of GK activity and correlated GK and g_Coup_ from Fig 3 top panel. D). As in A for simulations with bimodal distribution of GK activity and bimodal g_Coup_ from Fig 3 bottom panel. Data representative of 4-5 simulations with differing random number seeds.

**S2 Figure. Additional simulations with continuous distribution in GK activity with correlated g**_**Coup**_ **and g**_**KATP**_. A). Scatterplot of g_Coup_ vs. k_glc_ for each cell from a representative simulation where g_Coup_ is correlated with k_glc_ for simulation where GK activity is a modeled as a continuous distribution. B). Fraction of cells showing elevated [Ca^2+^] activity (active cells) vs. the percentage of cells hyperpolarized in islet from simulations with a continuous distribution in k_glc_ with correlated g_Coup_ and k_glc_ as in A. Hyperpolarized cells are chosen based on their GK rate which is correlated to g_Coup_. C). As in B. but comparing hyperpolarization in high GK cells in the presence (B) and absence (Fig 1c) of correlations in g_Coup_. D). as in A but from a simulation where g_Coup_ and k_glc_ and g_KATP_ (K_ATP_ channel conductance) are correlated. E). As in B. but for simulations where g_Coup_ and k_glc_ and g_KATP_ are correlated. F). As in C. but comparing high GK cells hyperpolarization from Fig 1c to high GK hyperpolarization from simulations where g_Coup_ and k_glc_ and g_KATP_ are correlated (E). Error bars are mean + s.e.m. Repeated measures one-way ANOVA with Tukey post-hoc analysis was performed for simulations in B and C (if there were any missing values a mixed effects model was used) and a Student’s t-test was performed for C and F (Welches t-test for unequal variances was used when variances were determined to be statistically different using an F-test) to test for significance. Significance values: ns indicates not significant (p>.05), * indicates significant difference (p<.05), ** indicates significant difference (p<.01), *** indicates significant difference (p<.001), **** indicates significant difference (p<.0001). Data representative of 5 simulations with differing random number seeds.

**S3 Figure. Simulations predicting effect of 50% reduction in coupling in simulations with continuous and bimodal distributions in GK activity**. A). Fraction of cells showing elevated [Ca^2+^] activity (active cells) vs. the percentage of cells hyperpolarized in islet from simulations with a continuous distribution as in Fig 1c but with 50% reduction in average coupling conductance (60pS) for all cells. Hyperpolarized cells are chosen based on their GK rate. B). As in A. but comparing hyperpolarization in high GK cells in simulations with full coupling (120pS – Fig 1c) and reduced coupling (60pS) from A. C). as in A but for bimodal simulations with reduced coupling (60pS). D). As in B but comparing bimodal distributions in GK with full coupling (120pS) from Fig 2d to bimodal simulations with reduced coupling (60pS) from C. Error bars are mean ± s.e.m. Student’s paired t-test was performed to test for significance for all simulations. Significance values: ns indicates not significant (p>.05), * indicates significant difference (p<.05), ** indicates significant difference (p<.01), *** indicates significant difference (p<.001), **** indicates significant difference. Data representative of 4-5 simulations with differing random number seeds.

**S4 Figure. Fraction of active cells in simulations where cells are uncoupled from the rest of the cells in the islet from Fig 4-6**. A). Fraction of cells showing elevated [Ca^2+^] activity (active cells) in simulated islets vs. the percentage of cells uncoupled in islet from simulations in Fig 4. B). As in A but for simulations in Fig 5. C). As in A but for simulations in Fig 6. D). As in A but for simulations in S6 Fig. Error bars are mean ± s.e.m. Repeated measures one-way ANOVA was performed for simulations in A and B and a Student’s paired t-test was performed for C and D to test for significance. Significance values: ns indicates not significant (p>.05), * indicates significant difference (p<.05), ** indicates significant difference (p<.01), *** indicates significant difference (p<.001), **** indicates significant difference. Data representative of 5 simulations with differing random number seeds.

**S5 Figure. Random removal of cells vs. random removal of a region of cells**. A). Schematic showing which cells are chosen to be removed when a random selection of cells is chosen across the islet. B). Schematic showing which cells are chosen to be removed when a *random region* of cells is chosen. C). The frequency of the islet after removal of 0%, 10%, or 30% of randomly chosen cells or from a random region. Error bars are mean + s.e.m. Student’s t-test was performed for 10% and a Welch’s t-test for unequal variances was used to test for significance at 30% of cells removed. Significance values: ns indicates not significant (p>.05), * indicates significant difference (p<.05), ** indicates significant difference (p<.01), *** indicates significant difference (p<.001), **** indicates significant difference. Data representative of 4-9 simulations with differing random number seeds.

**S6 Figure. Simulations predicting the effect of 50% reduction in coupling in simulations where high and low phase cells are removed under a continuous model**. A). Average frequency of islet when indicated populations of cells are removed from the simulated islet with 50% reduction in coupling conductance (60pS). B). Change in frequency of islet with indicated populations removed with respect to control islet with all cells present. C). Change in frequency when low phase cells are removed compared to average oscillation frequency of remaining cells that indicates the expected oscillation frequency. D). Same as C. but for simulations where high phase cells are removed. Error bars are mean ± s.e.m. Repeated measures one-way ANOVA with Tukey post-hoc analysis was performed for simulations in A-B and a Student’s paired t-test was performed for C and D to test for significance. Significance values: ns indicates not significant (p>.05), * indicates significant difference (p<.05), ** indicates significant difference (p<.01), *** indicates significant difference (p<.001), **** indicates significant difference (p<.0001). Data representative of 4 simulations with differing random number seeds.

**S7 Figure. Simulations predicting the effect of removing cells from individual populations of the bimodal model of low phase cells**. A). Average intrinsic oscillation frequencies of all cells, top 1% or 10% of high frequency cells, or low frequency cells when re-simulated in the absence of gap junction coupling from bimodal model of phase. B). Phase lag from islet average of top 1% or 10% of high frequency, low frequency cells, or random cells. C). Average k_glc_ from all cells, high frequency cells, or low frequency cells across simulated islet. D). Change in frequency of islet with indicated populations removed with respect to control islet with all cells present. E). Change in frequency when high frequency cells are removed compared to average oscillation frequency of remaining cells that indicates the expected oscillation frequency. F). Same as E. but for simulations where low frequency cells are removed. Error bars are mean ± s.e.m. Repeated measures one-way ANOVA with Tukey post-hoc analysis was performed for simulations in A-D and a Student’s paired t-test was performed for E and F to test for significance. Significance values: ns indicates not significant (p>.05), * indicates significant difference (p<.05), ** indicates significant difference (p<.01), *** indicates significant difference (p<.001), **** indicates significant difference (p<.0001). Data representative of 5 simulations with differing random number seeds.

**S8 Figure. Simulations predicting the effect of removing a region of high frequency cells from a bimodal model of low phase cells**. A). Schematic of frequency across simulated islet with a bimodal distribution in GK activity and a region of high frequency cells. B). Schematic of phase lag across simulated islet with a bimodal distribution in GK activity and a region of high frequency cells. C). Change in frequency of islet with indicated populations removed with respect to control islet with all cells present comparing bimodal model with a region of high frequency cells to a bimodal model with randomly distributed high frequency cells as in Fig 6. D). Change in frequency when high frequency region is removed compared to average oscillation frequency of remaining cells that indicates the expected oscillation frequency. Error bars represent mean ± s.e.m. Student’s t-test was performed for C and D (paired test) to test for significance. Significance values: ns indicates not significant (p>.05), * indicates significant difference (p<.05), ** indicates significant difference (p<.01), *** indicates significant difference (p<.001), **** indicates significant difference (p<.0001). Data representative of 5 simulations with differing random number seeds.

**S9 Figure. Analysis of changes in [Ca**^**2+**^**] wave dynamics when high/low phase or high/low frequency cells are removed from islet**. A). Change in mean duration of active phase when top 1%, 10% or 30% low/high phase cells are removed from simulations in Fig 4. B). Change in mean duration of silent phase when top 1%, 10% or 30% low/high phase cells are removed from simulations in Fig 4. C). Change in mean duty cycle when top 1%, 10% or 30% low/high phase cells are removed from simulations in 4. D). As in A for simulations when high/low frequency cells are removed from Fig 5. E). As in B for simulations when high/low frequency cells are removed from Fig 5. F). As in C for simulations when high/low frequency cells are removed from Fig 5. Error bars are mean ± s.e.m. Paired Student’s t-test was used to test for significance. Significance values: ns indicates not significant (p>.05), * indicates significant difference (p<.05), ** indicates significant difference (p<.01), *** indicates significant difference (p<.001), **** indicates significant difference (p<.0001). Data representative of 5 simulations with differing random number seeds.

**S1 Table: Parameters for bimodal phase low cell simulations**. Table describes the parameters that have heterogeneous populations in computational model. The mean of each population is determined from the mean parameter value from continuous simulations (See methods).

**S1 Files: Model code used in this study, in zip file**. Files include those used to generate data in figure 1, figure 2, figure 4 and figure 6.

