## Supplementary material for "Small subpopulations of β-cells do not drive islet oscillatory [Ca^2+^] dynamics via gap junction communication": S1 File: BetaCell.h

```
#ifndef BETACELL_H
#define BETACELL_H
#endif // BETACELL_H

#include <stdio.h>
#include <iostream>
#include <boost/numeric/odeint.hpp>
#include <boost/multi_array.hpp>
#include <boost/numeric/ublas/matrix.hpp>
#include <boost/numeric/ublas/io.hpp>
#include <cassert>
#include <cmath>
#include <ostream>
#include <fstream>
#include <vector>
#include <boost/random/detail/config.hpp>
#include <boost/random.hpp>
#include <boost/random/normal_distribution.hpp>
#include <boost/fusion/container/vector.hpp>
#include <boost/fusion/include/vector.hpp>
#include <boost/fusion/container/vector/vector_fwd.hpp>
#include <boost/fusion/include/vector_fwd.hpp>
#include <omp.h>
#include <string>
#include <algorithm>
#include <boost/math/distributions/skew_normal.hpp>
//#include "ChR2CurrentPulseV3.h"

#include "ChR2Class.h" 
#include <ctime>
using namespace std;
using namespace boost::numeric::odeint;
using namespace boost::math;

typedef boost::numeric::ublas::matrix<double> matrix_type;
//typedef boost::fusion::vector2<double,double> state_type;
 //boost::array<state_type::index,2> shape={{18,2}};
const double R = 8.3143;
const double Tem = 310.15;
const double F = 96.4867;
const double RTF = R*Tem / F;
const double RTF2 = RTF / 2;
//const double Glucose=8;
const double yini0 = -69.8663703359279;
const double yini1 = 7.92502913389466;
const double yini2 = 125.248586232226;
const double yini3 = 7.56138347594955E-05;
const double yini4 = 0.0047417301517704;

vector<double> potential;
vector<double> TimeLoc;
double pMutant;
int totalnodes;
using namespace boost;

double pKATP[2000];
double tOld = 0;
const int cellNumber = 1000;
const double hubMutant = 0.0; //specified number of hubs change here and will update for randomVars etc....
bool mutationInFollowers = true; //1 for mutation in followers and 0 for muation in hubs

typedef boost::array< double, 30 * cellNumber > state_type;
double gKATPar[2000];
double IKATPvec[2000];
double gCoup[2000];
double gKtoar[2000];
double PCaERar[2000];
double gKCaBKar[2000];
double ATPar[2000];
double PNACAar[2000];
double Prelar[2000];
double Popar[2000];
double KRev[2000];
double gChR2[2000];
double RandomSeed[2000];
vector<int> randomCellVector(cellNumber);
double alphaPos[390];
int* randPos = new int[cellNumber];
double data[100][100];
int NumCols = 100;
int NumRows = 100;
double tMax;
double cellClass[1000][2];
double cellPos[cellNumber][3];
int testBool = 1;
char* potentialname = new char;
char* timename = new char;
char* cellinfo = new char;
char* calciuminfo = new char;
char* sodiuminfo = new char;
char* potassiuminfo = new char;
char* caerinfo = new char;
char* atpinfo = new char;
char* adpinfo = new char;
char* ikatpinfo = new char;
const char* O1info = new char;
const char* O2info = new char;
const char* C1info = new char;
const char* C2info = new char;
//exocytosis variables
char* IRPinfo = new char;
char* PPinfo = new char;
char* DPinfo = new char;
char* FIPinfo = new char;
char* RIPinfo = new char;
char* Capinfo = new char;
char* Noiseinfo = new char;

//
double NN[cellNumber][15];
int CBool[cellNumber] = {};
double Frequency = 10;
double delt;
double period1 = 1000 * 1 / Frequency;

//[ stiff_system_definition
typedef boost::numeric::ublas::vector< double> vector_type;
typedef boost::numeric::ublas::matrix< double > matrix_type;
int numCores = 4;

void BetaSolver(vector_type x, vector_type dxdt, double  dt)
{

	double t;

	double Glucose = 8.0;
	double tStep = dt;

	for (t = 0; t < tMax; t = t + dt)
	{
		srand(1);
		int randNum;
		randNum = rand() % 100;

#pragma omp parallel num_threads(numCores)
#pragma omp for
		for (int j = 0; j < cellNumber; j++)
		{

			double hPos;

			ChR2Current ChR2Info;

			hPos = randPos[j] / (10 * 10);

			int	zPos = floor(hPos);

			int xyPos = randPos[j] % (10 * 10);

			double Vm = x[0 + j * 30];
			double Nai = x[1 + j * 30];
			double Ki = x[2 + j * 30];
			double Cai = x[3 + j * 30];
			double Caer = x[4 + j * 30];
			double ATP = x[5 + j * 30];
			double MgADP = x[6 + j * 30];
			double Re = x[7 + j * 30];
			double q_KDr = x[8 + j * 30];
			double d_CaL = x[9 + j * 30];
			double U_CaL = x[10 + j * 30];
			double fus = x[11 + j * 30];
			double p_KDr = x[12 + j * 30];
			double m_Kto = x[13 + j * 30];
			double h_Kto = x[14 + j * 30];
			double E1_tota = x[15 + j * 30];
			double I1 = x[16 + j * 30];
			double I2 = x[17 + j * 30];
			double O1 = x[18 + j * 30];
			double O2 = x[19 + j * 30];
			double C1 = x[20 + j * 30];
			double C2 = x[21 + j * 30];
			// exocytosis variables
			double IRP = x[22 + j * 30];
			double PP = x[23 + j * 30];
			double DP = x[24 + j * 30];
			double RES = x[25 + j * 30];
			double FIP = x[26 + j * 30];
			double RIP = x[27 + j * 30];
			double Cap = x[28 + j * 30];
			//noise
			double Pns = x[29 + j * 30];

			double KRe = 0.000126;
			double Kfa = 0.0000063;
			double Stoichi = 2.5;
			double Rvol = 2.5;
			double kATPCa = 0.187;
			double kATP = 0.000062;
			double kADPf = 0.0002;
			double kADPb = 0.00002;

			double Naout = 140;
			double Kout = 5.4;
			double Caout = 2.6;
			double Cm = 6.158;
			double voli = 764;
			double volER = 280;
			double fi = 0.01;
			double fer = 0.025;
			double totalATP = ATPar[j];

			double ADPb = totalATP - ATP - MgADP / 0.55;

			int Test;
			int hereV;

			double Vsum = 0;
			double currCellNum = j;
			double gCoupEQ;
			double Icoup = 0;

			int NNPos;
			int count = 0;
			for (int f = 0; f < 15; f++)
			{
				if (NN[j][f] != -1)
				{
					count = count + 1;
					NNPos = NN[j][f];


					//gCoupEQ=abs(gCoup[j]+gCoup[NNPos]);
					if (RandomSeed[j] > RandomSeed[NNPos])
					{
						gCoupEQ = gCoup[j];
					}
					else
					{
						gCoupEQ = gCoup[NNPos];
					}

					//Icoup=Icoup+((gCoupEQ-0.5*gCoupEQ)/(1+exp(0.07*((abs(Vm-x[22*NNPos]))-78)))+0.5*gCoupEQ)*(Vm-x[22*NNPos]);
				   // Icoup=Icoup+(((abs(gCoup[j]+gCoup[NNPos]))/2)+(1/(1+exp(abs(Vm-x[18*NNPos])/25)))*(Vm-x[18*NNPos]);
					Icoup = Icoup + ((gCoup[j] + gCoup[NNPos]) / 2)*(Vm - x[30 * NNPos]);
					//Icoup=Icoup+gCoupEQ*(Vm-x[22*NNPos]);    
					//Icoup = Icoup + max(gCoup[j],gCoup[NNPos]) * (Vm - x[30 * NNPos]);
													
				}
			}

			voli = 1.049*exp(0.456*count) + 738.7;


			Caer = yini4 + (fer*voli / 2 / volER)*(Cm / F / voli*(Vm - yini0) - (Nai - yini1) - (Ki - yini2) - 2 / fi*(Cai - yini3));

			// Icav

			double RCaLNa = 0.0000185;
			double RCaLK = 0.000367;
			double PCaL = 48.9;

			// Ikslow

			double PKslow = 0.2;
			double nKslow = 2.2;
			double KdKslow = 0.00074;

			// Ikdr

			double pKDr = 2.1;

			double P_PMCA = 1.56;
			double  K_PMCA = 0.00014;

			// ICRAN params

			double PCRAN = 0.00764;
			double  KCaer = 0.003;
			double RNa_K_CRAN = 0.8;
			// double pIbNSC=0.00396 ;
			//double pIbNSC=0.00115;
			double pIbNSC = 0.00396;

			double KTRPM = 0.00076;
			double pTRPM = 0.0234;
			double  RNa_K_TRPM = 0.8;


			//Ikatp params

			double kdd = 0.01;
			double ktt = 0.05;
			double ktd = 0.026;
			double gKATP = 2.31;

			//IKto params

			double GKto = gKtoar[j];


			// NA/Ca exchange params

			double KdNao = 87.5;
			double  KdCao = 1.38;
			double  KdNai = 20.75;
			double  KdCai = 0.0184;
			double  k3 = 1;
			double  k4 = 1;
			double AmpINaCa = PNACAar[j];


			// Na/K pump
			double Pii = 1.9;
			double Proton = 0.0001;
			//double Kd_MgATP=0.06 ;
			double Kd_MgATP = 0.6;
			double Kd_Nao0 = 26.8;
			double Kd_Nai0 = 5.0;
			double Kd_Ko0 = 0.8;
			double Kd_Ki0 = 18.8;
			double delta_Nao = 0.44;
			double delta_Nai = -0.14;
			double delta_Ko = 0.23;
			double delta_Ki = -0.14;
			double k1_plus = 1.253;
			double k2_plus = 0.139;
			double k3_plus = 6.96;
			double k4_plus = 0.52;
			double k1_minus = 0.139;
			double k2_minus = 0.0139;
			double k3_minus = 13900;
			double k4_minus = 0.348;
			double PNaK = 350;

			//Metabolism

			double Nt = 10;

			//ER dynamics

			double PCaER = PCaERar[j];
			double KCarp = 0.0005;
			double Jserca = PCaER*Cai*Cai / (Cai*Cai + KCarp*KCarp);
			double Pleak = Prelar[j];
			double Jout = Pleak*(Caer - Cai);

			//Glycolysis and oxidative phosph;
			double KmATP = 0.5;
			double hgl = 2.5;
			double Kg = 13;
			double fGlu = ATP / (KmATP + ATP)*pow(Glucose, hgl) / (pow(Kg, hgl) + pow(Glucose, hgl));
			//Check the Pop value
			double Pop = Popar[j];
			double Kop = 0.02;
			double JOP = Pop*Re*pow(MgADP, 2) / (pow(MgADP, 2) + pow(Kop, 2));


			//Constant field equations

			double Denom1 = (1 - exp(-Vm / RTF));
			double NaCF = (Vm / RTF) / Denom1*(Nai - Naout*exp(-Vm / RTF));
			double KCF = (Vm / RTF) / Denom1*(Ki - Kout*exp(-Vm / RTF));
			double Denom2 = (1 - exp(-Vm / RTF2));
			double CaCF = (Vm / RTF2) / Denom2*(Cai - Caout*exp(-Vm / RTF2));


			//reverse potentials

			double EK = RTF*log(Kout / Ki);
			double ENa = RTF*log(Naout / Nai);
			double ECa = RTF*log(Caout / Cai) / 2;


			double IbNSC1 = pIbNSC*NaCF;
			double IbNSC2 = 0.01*KCF;
			double IbNSC0 = IbNSC1 + IbNSC2;

			double dalpha = 1 / (0.88*exp(-(Vm - 3) / 50) + 0.09*exp(-(Vm - 3) / 600));
			double dbeta = 1 / (5.48*exp((Vm - 3) / 12) + 1.245*exp((Vm - 3) / 30));
			double VpOpen = pow(d_CaL, 2);


			//Calcium gating function

			double SingleiCaL = 0.0676*CaCF;
			double Ualpha = 0.0042 * 2;
			double Ubeta = 0.1159*(-1.15*SingleiCaL*VpOpen + Cai) * 2;
			//G
			//Ultraslow gate
			double usalpha = 1 / (75000 * exp(Vm / 34));
			double usbeta = 1 / (5000 * exp(-Vm / 19) + 500 * exp(-Vm / 100));

			//ICaL

			//double RundownATP=0.3+0.7/(1+pow((0.7/ATP),3));
			double RundownATP = 1 / (1 + pow((1.4 / ATP), 3));
			double pO = (VpOpen*U_CaL*(0.4 + 0.6*fus))*RundownATP;
			double ICaL1 = RCaLNa*PCaL*pO*NaCF;
			double ICaL2 = RCaLK*PCaL*pO*KCF;
			double ICaL3 = PCaL*pO*CaCF;
			double ICaL0 = ICaL1 + ICaL2 + ICaL3;

			//Ikdr

			double alphap = 1.1 / (25 * exp(-(Vm - 3) / 8) + 1 * exp(-(Vm - 3) / 100));
			double betap = 1.1 / (25 * exp(Vm / 100));
			double alphaq = 1 / 800;
			double betaq = 1 / (1000 * exp(-Vm / 8) + 100 * exp(-Vm / 100));
			double IKDr2 = pKDr*p_KDr*(0.6*q_KDr + 0.4)*KCF;
			double IKDr0 = IKDr2;

			//Ikto

			double alpham = 0.4 / (5.46*exp(-Vm / 20));
			double betam = 0.4 / (2.48*exp(Vm / 60));
			double alphah = 1.7 / (969 * exp(Vm / 500));
			double betah = 1.7 / (13.2*exp(-Vm / 9) + 6.93*exp(-Vm / 1000));
			double IKto2 = GKto*m_Kto*h_Kto*(Vm - EK);
			double IKto0 = IKto2;

			//ITRPM

			double PoTRPM = 1 / (1 + pow((KTRPM / Cai), 1.7));
			double ITRPM1 = pTRPM*RNa_K_TRPM*NaCF*PoTRPM;
			double ITRPM2 = pTRPM*KCF*PoTRPM;
			double ITRPM0 = ITRPM1 + ITRPM2;


			//IKATP
			double residual = 0.0;
			double kPrime = 1.0;
			double poPrime = 1.0;

			double pOatp = (residual)+((1 - residual)*(poPrime*0.08*(1 + 2 * MgADP / kdd) + 0.89*pow((MgADP / kdd), 2)) / pow((1 + MgADP / kdd), 2) / (1 + 0.45*MgADP / ktd + ATP / (kPrime*ktt)));

			double IChR2 = 0;

			double dO1 = 0;
			double dO2 = 0;
			double dC1 = 0;
			double dC2 = 0;
			//IChR2=ChR2Current(10000,dt,15000);
			//double IChR2=gChR2[j]*Vm*0.3*(O1+0.04*O2);


			//ADD MUTATIONS HERE//////////////////////////////////////////////////////////////////////////
			//double numMuts = 100.0; //use this if not using the 0B file to specify a percentage of mutation
			//Glucokinase mutation - make the rate of glycolysis 0
			//if (t == 10000)
			//{
				//check that pMutant is correct
			//	cout<<"pMutant"<<pMutant<<endl;
			//	cout<<"pMut times 1000"<<pMutant*1000.0<<endl;
			//}	
			//
			double mutationstart = 0;
			if (mutationInFollowers)
			{
				mutationstart = mutationstart + hubMutant*cellNumber;
			}
			if (t > 10000)
			{
				if (j >= mutationstart & j < mutationstart+pMutant*cellNumber) //use this for a random mutation		
				//if (KRev[j] > pMutant) //use pMutant to set a threshold for max glycolysis mutant
				//if (KRev[j] < pMutant) //use pMutant to set a threshold for min glycolysis mutant
				//if (gKATPar[j] > pMutant) //use pMutant to set a threshold for KATP max conductance
				//if (gKATPar[j] < pMutant) //use pMutant to set a threshold for KATP min conductnce

				{
					//use this for glycolysis mutant
					//KRev[j] = 0.0; 

					//use this for KATP Kir6.2 Mutant equation
					pOatp=0.5*pOatp+(1-0.5)*1.0;
					if (pOatp<0.0)
           				{
						pOatp=0.0;
            				}
				}
			}
			//////////////////////////////////////////////////////////////////////////////////

			
/*			double KRevInt; 
			 			
			if (j < hubMutant * cellNumber) //use this for a random mutation		
			{	//if (KRev[j] > pMutant) //use pMutant to set a threshold for max glycolysis mutant

					//use this for glycolysis mutant
				KRevInt = 2.0 * 5.5 * KRev[j];//used 9/8 because took 8/9 of all values in randomvars.cpp 
			} else {
				KRevInt = KRev[j];
			}*/
			
			//ikatp with noise
			// double IKATP2=gKATPar[j]*pOatp*(Vm-EK)*(1+Pns);
			double IKATP2 = gKATPar[j] * pOatp*(Vm - EK)*(1);
			double IKATP0 = IKATP2;
			IKATPvec[j] = IKATP2;
			//INaK

			double fVm = F*Vm / (R*Tem);
			double Kd_Nao = Kd_Nao0*exp(delta_Nao*fVm);
			double Kd_Nai = Kd_Nai0*exp(delta_Nai*fVm);
			double Kd_Ko = Kd_Ko0*exp(delta_Ko*fVm);
			double Kd_Ki = Kd_Ki0*exp(delta_Ki*fVm);
			double Nai_ = Nai / Kd_Nai;
			double Naout_ = Naout / Kd_Nao;
			double Ki_ = Ki / Kd_Ki;
			double Kout_ = Kout / Kd_Ko;
			double MgATP_ = ATP / Kd_MgATP;

			double a1_plus = (k1_plus*pow(Nai_, 3.0)) / (pow((1 + Nai_), 3) + pow((1 + Ki_), 2) - 1);
			double a2_plus = k2_plus;
			double a3_plus = k3_plus*pow(Kout_, 2) / (pow((1 + Naout_), 2) + pow((1 + Kout_), 2) - 1);
			double a4_plus = k4_plus*MgATP_ / (1 + MgATP_);
			double a1_minus = k1_minus*MgADP;
			double a2_minus = k2_minus*pow(Naout_, 3) / (pow((1 + Naout_), 3) + pow((1 + Kout_), 2) - 1);
			double a3_minus = k3_minus*Pii*Proton / (1 + MgATP_);
			double a4_minus = k4_minus*pow(Ki_, 2) / (pow((1 + Nai_), 3) + pow((1 + Ki_), 2) - 1);

			double denomi = (a1_minus + a1_plus)*a2_minus*a3_minus + a1_plus*a2_plus*(a3_plus + a3_minus) + a2_plus*a3_plus*(a4_plus + a4_minus) + (a2_plus + a2_minus)*a3_minus*a4_minus + (a1_minus + a1_plus)*a3_plus*a4_plus + a1_minus*(a3_plus + a3_minus)*a4_minus + a1_plus*(a2_plus + a2_minus)*a4_plus + a1_minus*a2_minus*(a4_plus + a4_minus);

			double numer = a1_plus*a2_plus*a3_plus*a4_plus - a1_minus*a2_minus*a3_minus*a4_minus;

			double iglc = (0.4 + 0.6*exp(-Glucose / 5.84));

			double vcyc = (numer / denomi)*iglc;

			double INaK0 = PNaK*vcyc;
			double INaK1 = 3 * INaK0;
			double INaK2 = -2 * INaK0;

			// INaCa slow

			double pE1Na = 1 / (1 + pow((KdNai / Nai), 3)*(1 + Cai / KdCai));
			double pE1Ca = 1 / (1 + (KdCai / Cai)*(1 + pow((Nai / KdNai), 3)));
			double pE2Na = 1 / (1 + pow((KdNao / Naout), 3)*(1 + Caout / KdCao));
			double pE2Ca = 1 / (1 + (KdCao / Caout)*(1 + pow((Naout / KdNao), 3)));
			double k1 = exp(0.32*Vm / RTF);
			double k2 = exp((0.32 - 1)*Vm / RTF);
			double fCa = Cai / (Cai + 0.004);
			double alpha1 = pE1Na*(fCa*0.002 + (1 - fCa)*0.0015);
			double beta1 = fCa*0.0012 + (1 - fCa)*0.0000005;
			double alpha2 = fCa*0.00003 + (1 - fCa)*0.01;
			double beta2 = fCa*0.09 + (1 - fCa)*0.0001;

			// IPMCA

			double IPMCA0 = P_PMCA*pow(Cai, 2) / (pow(Cai, 2) + pow(K_PMCA, 2));
			double IPMCA1 = -IPMCA0;
			double IPMCA3 = 2 * IPMCA0;

			double kf = k2*pE2Na + k4*pE2Ca;
			double kb = k1*pE1Na + k3*pE1Ca;
			double E2_tot = 1 - E1_tota - I1 - I2;
			double INaCa0 = AmpINaCa*(k1*pE1Na*E1_tota - k2*pE2Na*E2_tot);
			double INaCa1 = 3 * INaCa0;
			double INaCa3 = -2 * INaCa0;

			// ICRAN

			//IKSLOW

			double PoKslow = 1 / (1 + pow((KdKslow / Cai), nKslow));
			double IKslow2 = PKslow*PoKslow*KCF;
			double IKslow0 = IKslow2;

			double PoCRAN = 1 / (1 + exp((Caer - KCaer) / 0.003));
			double ICRAN1 = PCRAN*RNa_K_CRAN*PoCRAN*NaCF;
			double ICRAN2 = PCRAN*PoCRAN*KCF;
			double ICRAN3 = PCRAN*PoCRAN*CaCF * 20;
			double ICRAN0 = ICRAN1 + ICRAN2 + ICRAN3;


			double Itot = IbNSC0 + IKDr0 + IKto0 + IKATP0 + ITRPM0 + ICaL0 + INaK0 + INaCa0 + IPMCA0 + IKslow0 + ICRAN0 + Icoup + IChR2;
			double INatot = IbNSC1 + ITRPM1 + ICaL1 + INaK1 + INaCa1 + IPMCA1 + ICRAN1 + Icoup / 3 + IChR2 / 2;
			double IKtot = IbNSC2 + IKDr2 + IKto2 + IKATP2 + ITRPM2 + ICaL2 + INaK2 + IKslow2 + ICRAN2 + Icoup / 3;
			double ICatot = ICaL3 + INaCa3 + IPMCA3 + ICRAN3 + Icoup / 3 + IChR2 / 2;

			double JGlyc = KRev[j] * fGlu*(Nt - Re);
			double JBox = Kfa*(Nt - Re);

			// noise parameters
			double noiseRand = (rand() % 3) - 1;
			double noisey = noiseRand / 80;
			double taup = 500;

			//exocytosis rates between pools

				// FUSION POOL EXOCYTOSIS % %
			long double fusionMax = .030;//30 gran/sec
			long double nFuse = 4;
			long double K_I = 0.0022;//2.2uM
			//Hill equation for fusion with plasma membrane
			long double fusion_I = fusionMax * (pow(Cai, nFuse)) / (pow(Cai, nFuse) + pow(K_I, nFuse));

			//%exocytosis rates between pools%%
			//% r2max = .00014/1000; cN=2; cKi = 2;
			//rate into immediately releasable pool
			long double r1 = 0.020 / 1000;
			//rate out of IRP
			long double r_1 = 0.025 / 1000;
			//rates into/out of primed and docked pools, and from reserve
			long double r2 = 0.00012 / 1000;
			long double r_2 = 0.0012 / 1000;
			long double Rres = 0.00005 / 1000;
			long double R_res = 0.00004 / 1000;

			// for cAMP later
			long double CaN = 4;
			long double Kp = 2.3E-4;
			// %r2 = r2max * Cai^CaN/(Cai^CaN + Kp^CaN);
			long double cN = 4;
			long double cKi = 2.3E-3;
			// %R_res = R_resmax * ((cAMP^cN)/(cAMP^cN + cKi^cN));

			//rate of movement from fusion to release pool (ms)
			long double u2 = 0.003;
			//rate of release (ms)
			long double u3 = 0.00004;

			//
			long double Lflux = 5.18E-15*ICaL3 / (.00383E-3);
			long double F_md = .01;

			dxdt[0 + j * 30] = -Itot / Cm;
			dxdt[1 + j * 30] = -INatot / (F*voli);
			dxdt[2 + j * 30] = -IKtot / (F*voli);
			dxdt[3 + j * 30] = fi*(-ICatot / (2 * F) - Jserca + Jout) / voli;
			dxdt[4 + j * 30] = fer*(Jserca - Jout) / volER;
			dxdt[5 + j * 30] = JOP - ((INaK0 + IPMCA0) / F + Jserca / 2) / voli - (kATP + kATPCa*Cai)*ATP;
			dxdt[6 + j * 30] = -0.55*(JOP - ((INaK0 + IPMCA0) / F + Jserca / 2) / voli - (kATP + kATPCa*Cai)*ATP) + 0.55*kADPb*ADPb - kADPf*MgADP;
			dxdt[7 + j * 30] = JGlyc + JBox - JOP*Rvol / Stoichi;
			dxdt[8 + j * 30] = alphaq*(1 - q_KDr) - betaq*q_KDr;
			dxdt[9 + j * 30] = dalpha*(1 - d_CaL) - dbeta*d_CaL;
			dxdt[10 + j * 30] = Ualpha*(1 - U_CaL) - Ubeta*U_CaL;
			dxdt[11 + j * 30] = usalpha*(1 - fus) - usbeta*fus;
			dxdt[12 + j * 30] = alphap*(1 - p_KDr) - betap*p_KDr;
			dxdt[13 + j * 30] = alpham*(1 - m_Kto) - betam*m_Kto;
			dxdt[14 + j * 30] = alphah*(1 - h_Kto) - betah*h_Kto;
			dxdt[15 + j * 30] = E2_tot*kf + I1*beta1 + I2*beta2 - E1_tota*(kb + alpha1 + alpha2);
			dxdt[16 + j * 30] = E1_tota*alpha1 - I1*beta1;
			dxdt[17 + j * 30] = E1_tota*alpha2 - I2*beta2;
			dxdt[18 + j * 30] = dO1;
			dxdt[19 + j * 30] = dO2;
			dxdt[20 + j * 30] = dC1;
			dxdt[21 + j * 30] = dC2;
			//exocytosis ODEs
			dxdt[22 + j * 30] = r1 * PP - r_1 * IRP - fusion_I * IRP;
			dxdt[23 + j * 30] = r_1* IRP - (r1 + r_2)*PP + r2*DP;
			dxdt[24 + j * 30] = r_2* PP - r2*DP + Rres*RES - R_res*RES;
			dxdt[25 + j * 30] = Rres*RES - R_res*RES;
			dxdt[26 + j * 30] = fusion_I*IRP - u2*FIP;
			dxdt[27 + j * 30] = u2*FIP - u3*RIP;
			dxdt[28 + j * 30] = .0035*fusion_I*IRP;
			dxdt[29 + j * 30] = (-Pns) / taup - (Pns / taup) + noisey;
		}

#pragma omp parallel num_threads(numCores)
#pragma omp for
		for (int j = 0; j < cellNumber; j++)
		{
			x[0 + j * 30] = x[0 + j * 30] + dxdt[0 + j * 30] * tStep;
			x[1 + j * 30] = x[1 + j * 30] + dxdt[1 + j * 30] * tStep;
			x[2 + j * 30] = x[2 + j * 30] + dxdt[2 + j * 30] * tStep;
			x[3 + j * 30] = x[3 + j * 30] + dxdt[3 + j * 30] * tStep;
			x[4 + j * 30] = x[4 + j * 30] + dxdt[4 + j * 30] * tStep;
			x[5 + j * 30] = x[5 + j * 30] + dxdt[5 + j * 30] * tStep;
			x[6 + j * 30] = x[6 + j * 30] + dxdt[6 + j * 30] * tStep;
			x[7 + j * 30] = x[7 + j * 30] + dxdt[7 + j * 30] * tStep;
			x[8 + j * 30] = x[8 + j * 30] + dxdt[8 + j * 30] * tStep;
			x[9 + j * 30] = x[9 + j * 30] + dxdt[9 + j * 30] * tStep;
			x[10 + j * 30] = x[10 + j * 30] + dxdt[10 + j * 30] * tStep;
			x[11 + j * 30] = x[11 + j * 30] + dxdt[11 + j * 30] * tStep;
			x[12 + j * 30] = x[12 + j * 30] + dxdt[12 + j * 30] * tStep;
			x[13 + j * 30] = x[13 + j * 30] + dxdt[13 + j * 30] * tStep;
			x[14 + j * 30] = x[14 + j * 30] + dxdt[14 + j * 30] * tStep;
			x[15 + j * 30] = x[15 + j * 30] + dxdt[15 + j * 30] * tStep;
			x[16 + j * 30] = x[16 + j * 30] + dxdt[16 + j * 30] * tStep;
			x[17 + j * 30] = x[17 + j * 30] + dxdt[17 + j * 30] * tStep;
			x[18 + j * 30] = x[18 + j * 30] + dxdt[18 + j * 30] * tStep;
			x[19 + j * 30] = x[19 + j * 30] + dxdt[19 + j * 30] * tStep;
			x[20 + j * 30] = x[20 + j * 30] + dxdt[20 + j * 30] * tStep;
			x[21 + j * 30] = x[21 + j * 30] + dxdt[21 + j * 30] * tStep;
			x[22 + j * 30] = x[22 + j * 30] + dxdt[22 + j * 30] * tStep;
			x[23 + j * 30] = x[23 + j * 30] + dxdt[23 + j * 30] * tStep;
			x[24 + j * 30] = x[24 + j * 30] + dxdt[24 + j * 30] * tStep;
			x[25 + j * 30] = x[25 + j * 30] + dxdt[25 + j * 30] * tStep;
			x[26 + j * 30] = x[26 + j * 30] + dxdt[26 + j * 30] * tStep;
			x[27 + j * 30] = x[27 + j * 30] + dxdt[27 + j * 30] * tStep;
			x[28 + j * 30] = x[28 + j * 30] + dxdt[28 + j * 30] * tStep;
			x[29 + j * 30] = x[29 + j * 30] + dxdt[29 + j * 30] * tStep;
		}


		if ((t - tOld) > 100)
		{
			tOld = t;
			std::ofstream outfile;
			outfile.open(potentialname, ios::app);
			std::ofstream outfileCalcium;
			outfileCalcium.open(calciuminfo, ios::app);
			ofstream outfileSodium;
			outfileSodium.open(sodiuminfo, ios::app);
			ofstream outfilePotassium;
			outfilePotassium.open(potassiuminfo, ios::app);
			ofstream outfileCaer;
			outfileCaer.open(caerinfo, ios::app);
			ofstream outfileATP;
			outfileATP.open(atpinfo, ios::app);
			ofstream outfileADP;
			outfileADP.open(adpinfo, ios::app);
			ofstream outfileO1;
			ofstream outfileO2;
			ofstream outfileC1;
			ofstream outfileC2;
			outfileO1.open(O1info, ios::app);
			outfileO2.open(O2info, ios::app);
			outfileC1.open(C1info, ios::app);
			outfileC2.open(C2info, ios::app);
			//exocytosis files
			ofstream outfileIRP;
			outfileIRP.open(IRPinfo, ios::app);
			ofstream outfilePP;
			outfilePP.open(PPinfo, ios::app);
			ofstream outfileDP;
			outfileDP.open(DPinfo, ios::app);
			ofstream outfileFIP;
			outfileFIP.open(FIPinfo, ios::app);
			ofstream outfileRIP;
			outfileRIP.open(RIPinfo, ios::app);
			ofstream outfileCap;
			outfileCap.open(Capinfo, ios::app);
			ofstream outfileNoise;
			outfileNoise.open(Noiseinfo, ios::app);

			for (int k = 0; k < 30 * cellNumber; k = k + 30)
			{
				outfile << x[k] << ' ';
				outfileSodium << x[k + 1] << ' ';
				outfilePotassium << x[k + 2] << ' ';
				outfileCaer << x[k + 4] << ' ';
				outfileCalcium << x[k + 3] << ' ';
				outfileATP << x[k + 5] << ' ';
				outfileADP << x[k + 6] << ' ';
				outfileO1 << x[k + 18] << ' ';
				outfileO2 << x[k + 19] << ' ';
				outfileC1 << x[k + 20] << ' ';
				outfileC2 << x[k + 21] << ' ';
				outfileIRP << x[k + 22] << ' ';
				outfilePP << x[k + 23] << ' ';
				outfileDP << x[k + 24] << ' ';
				outfileFIP << x[k + 26] << ' ';
				outfileRIP << x[k + 27] << ' ';
				outfileCap << x[k + 28] << ' ';
				outfileNoise << x[k + 29] << ' ';
			}
			outfile << ' ' << endl;
			outfileCalcium << ' ' << endl;
			outfileSodium << ' ' << endl;
			outfilePotassium << ' ' << endl;
			outfileCaer << ' ' << endl;
			outfileATP << ' ' << endl;
			outfileADP << ' ' << endl;
			outfileO1 << ' ' << endl;
			outfileO2 << ' ' << endl;
			outfileC1 << ' ' << endl;
			outfileC2 << ' ' << endl;
			outfileIRP << ' ' << endl;
			outfilePP << ' ' << endl;
			outfileDP << ' ' << endl;
			outfileFIP << ' ' << endl;
			outfileRIP << ' ' << endl;
			outfileCap << ' ' << endl;
			outfileNoise << ' ' << endl;
			outfile.close();
			outfileATP.close();
			outfileADP.close();
			outfileO1.close();
			outfileO2.close();
			outfileC1.close();
			outfileC2.close();
			outfileCalcium.close();
			outfileSodium.close();
			outfilePotassium.close();
			outfileCaer.close();
			outfileIRP.close();
			outfilePP.close();
			outfileDP.close();
			outfileFIP.close();
			outfileRIP.close();
			outfileCap.close();
			outfileNoise.close();

		}
		std::ofstream outfile2;
		outfile2.open(timename, ios::app);
		outfile2 << pMutant << endl;
		outfile2.close();

	}

#pragma omp barrier
};
```
