## Supplementary material for "Small subpopulations of β-cells do not drive islet oscillatory [Ca^2+^] dynamics via gap junction communication": S1 File: box.C

```
/* 
   Updated July 24, 2009 to include hardwall boundary condition option, as 
   well as polydisperse spheres.  

   Packing of hard spheres via molecular dynamics
   Developed by Monica Skoge, 2006, Princeton University
   Contact: Monica Skoge (mskoge.princeton.edu) with questions
   This code may be used, modified and distributed freely.
   Please cite:
   
   "Packing Hyperspheres in High-Dimensional Euclidean Spaces"
   	M. Skoge, A. Donev, F. H. Stillinger and S. Torquato, 2006
	
   if you use these codes.	
*/

#include "box.h"
#include <iostream>
#include <math.h>
#include <stdlib.h>
#include <time.h>
#include <iomanip>
#include <cstdlib>

//==============================================================
//==============================================================
//  Class Box: Fills box with hardspheres to given packing fraction
//  and evolves spheres using molecular dynamics!
//==============================================================
//==============================================================


//==============================================================
// Constructor
//==============================================================
box::box(int N_i, double r_i, double growthrate_i, double maxpf_i, 
	 double bidispersityratio_i, double bidispersityfraction_i, 
	 double massratio_i, int hardwallBC_i):
  r(r_i),          
  N(N_i),
  growthrate(growthrate_i),
  h(N_i+1),
  maxpf(maxpf_i),
  bidispersityratio(bidispersityratio_i),
  bidispersityfraction(bidispersityfraction_i),
  massratio(massratio_i),
  hardwallBC(hardwallBC_i)
{
  ngrids = Optimalngrids2(r);
  cells.set_size(ngrids);
 
  s = new sphere[N];
  binlist = new int[N];
  x = new vector<DIM>[N];        
  h.s = s;

  gtime = 0.;
  rtime = 0.;
  ncollisions = 0;
  ntransfers = 0;
  nchecks = 0;
  neventstot = 0;
  ncycles = 0;
  xmomentum = 0.; 
  pressure = 0.;
  collisionrate = 0.;

  cells.set_size(ngrids);
  cells.initialize(-1);      // initialize cells to -1
  srandom(1);        // initialize the random number generator
  for (int i=0; i<N; i++)    // initialize binlist to -1
    binlist[i] = -1;
  
  time(&start);
}


//==============================================================
// Destructor
//==============================================================
box::~box() 
{
  delete[] s;
  delete[] binlist;
  delete[] x;
}


//==============================================================
// ReadFile
//==============================================================
void box::ReadPositions(const char* filename)
{  
  // open file to read in arrays
  std::ifstream infile(filename);
  
  infile.ignore(256, '\n');  // ignore the dim line
  infile.ignore(256, '\n');  // ignore the #sphere 1 line
  infile.ignore(256, '\n');  // ignore the #sphere line
  infile.ignore(256, '\n');  // ignore the diameter line
  infile.ignore(1000, '\n'); // ignore the 100 010 001 line
  infile.ignore(256, '\n');  // ignore the T T T line

  for (int i=0; i<N; i++)
    {
      infile >> s[i].r;      // read in radius    
      infile >> s[i].gr;     // read in growth rate
      infile >> s[i].m;      // read in mass
      for (int k=0; k<DIM; k++)  
	infile >> s[i].x[k]; // read in position 
    }
  infile.close();
}


//==============================================================
// Recreates all N spheres at random positions
//==============================================================
void box::RecreateSpheres(const char* filename, double temp)
{
  ReadPositions(filename);  // reads in positions of spheres
  VelocityGiver(temp);      // gives spheres initial velocities
  AssignCells();            // assigns spheres to cells
  SetInitialEvents();
}


//==============================================================
// Creates all N spheres at random positions
//==============================================================
void box::CreateSpheres(double temp)
{
  int Ncurrent = 0;
  for(int i=0; i<N; i++)
    {
      CreateSphere(Ncurrent);
      Ncurrent++;
    }
  if (Ncurrent != N)
    std::cout << "problem! only made " << Ncurrent << " out of " << N << " desired spheres" << std::endl;
  
  VelocityGiver(temp);
  SetInitialEvents();
}

   
//==============================================================
// Creates a sphere of random radius r at a random unoccupied position 
//==============================================================
void box::CreateSphere(int Ncurrent)
{
  int keeper;    // boolean variable: 1 means ok, 0 means sphere already there
  int counter = 0;   // counts how many times sphere already exists
  vector<DIM> xrand;  // random new position vector
  double d = 0.;
  double radius;
  double growth_rate;
  double mass;
  int species;
  
  if (Ncurrent < bidispersityfraction*N)
    {
      radius = r;
      growth_rate = growthrate;
      mass = 1.;
      species = 1;
    }
  else
    {
      radius = r*bidispersityratio;
      growth_rate = growthrate*bidispersityratio;
      mass = massratio;
      species = 2;
    }
  
  while (counter<1000)
    {
      keeper = 1;
      
      for(int k=0; k<DIM; k++) 
	xrand[k] = ((double)random()/(double)RAND_MAX)*SIZE;
      
      for (int i=0; i<Ncurrent; i++)  // check if overlapping other spheres
	{
	  d=0.;
	  if (hardwallBC)
	    {
	      for (int k=0; k<DIM; k++) 
		{     
		  d += (xrand[k] - s[i].x[k])*(xrand[k] - s[i].x[k]);
		}
	    }
	  else
	    {
	      for (int k=0; k<DIM; k++)
		{
		  if ((xrand[k] - s[i].x[k])*(xrand[k] - s[i].x[k]) > SIZE*SIZE/4.)
		    {
		      if (xrand[k] > SIZE/2.)  // look at right image
			d += (xrand[k] - (s[i].x[k]+SIZE))*
			  (xrand[k] - (s[i].x[k]+SIZE));
		      else                     // look at left image
			d += (xrand[k] - (s[i].x[k]-SIZE))*
			  (xrand[k] - (s[i].x[k]-SIZE));
		    }
		  else
		    d += (xrand[k] - s[i].x[k])*(xrand[k] - s[i].x[k]);
		}
	    }
	  if (d <= (radius + s[i].r)*(radius + s[i].r)) // overlapping!
	    {
	      keeper = 0;
	      counter++;
	      break;
	    }
	}
      
      if (hardwallBC)
	{
	  for (int k=0; k<DIM; k++)    // check if overlapping wall
	    {     
	      if ((xrand[k] <= radius) || (SIZE - xrand[k] <= radius)) // touching wall
		{
		  keeper = 0;
		  counter++; 
		  break;
		}
	    }
	}
      
      if (keeper == 1)
	break;   
      
      if (counter >= 1000)
	{
	  std::cout << "counter >= 1000" << std::endl;
	  exit(-1);
	}
    }

  // now convert xrand into index vector for cells
  vector<DIM,int> cell;
  cell = vector<DIM>::integer(xrand*((double)(ngrids))/SIZE);
  
  s[Ncurrent] = sphere(Ncurrent, xrand, cell, gtime, radius, growth_rate, 
		       mass, species);
  
  //first check to see if entry at cell
  if (cells.get(cell) == -1) //if yes, add Ncurrent to cells gridfield
    cells.get(cell) = Ncurrent;

  else  // if no, add i to right place in binlist
    {  
      int iterater = cells.get(cell); // now iterate through to end and add Ncurrent
      int pointer = iterater;
      while (iterater != -1)
	{
	  pointer = iterater;
	  iterater = binlist[iterater];
	}
      binlist[pointer] = Ncurrent;
    }

}


//==============================================================
// Assign cells to spheres read in from existing configuration
//==============================================================
void box::AssignCells()
{
  for (int i=0; i<N; i++)
    {
      // now convert x into index vector for cells
      vector<DIM,int> cell;
      cell = vector<DIM>::integer(s[i].x*((double)(ngrids))/SIZE);
      s[i].cell = cell;
      
      //first check to see if entry at cell
      if (cells.get(cell) == -1) //if yes, add Ncurrent to cells gridfield
	cells.get(cell) = i;
      
      else  // if no, add i to right place in binlist
	{  
	  int iterater = cells.get(cell); // now iterate through to end and add Ncurrent
	  int pointer = iterater;
	  while (iterater != -1)
	    {
	      pointer = iterater;
	      iterater = binlist[iterater];
	    }
	  binlist[pointer] = i;
	}
    }
}

	
//==============================================================
// Velocity Giver, assigns initial velocities from Max/Boltz dist.
//==============================================================
void box::VelocityGiver(double T)
{
  for (int i=0; i<N; i++)
    {
      for (int k=0; k<DIM; k++)
	{
	  if (T==0.)
	    s[i].v[k] = 0.;
	  else
	    s[i].v[k] = Velocity(T);
	}
    }
}


//==============================================================
// Velocity, gives a single velocity from Max/Boltz dist.
//==============================================================
double box::Velocity(double T)
{
  double rand;                       // random number between -0.5 and 0.5
  double sigmasquared = T;    // Assumes M = mass of sphere = 1
  double sigma = sqrt(sigmasquared); // variance of Gaussian
  double stepsize = 1000.;           // stepsize for discretization of integral
  double vel = 0.0;                  // velocity
  double dv=sigma/stepsize;
  double p=0.0;
  
  rand = (double)random() / (double)RAND_MAX - 0.5;
  if(rand < 0) 
    {
      rand = -rand;
      dv = -dv;
    }
  
  while(fabs(p) < rand) // integrate until the integral equals rand
    {
      p += dv * 0.39894228 * exp(-vel*vel/(2.*sigmasquared))/sigma;
      vel += dv;
    }
  return vel;
}


//==============================================================
// Finds next events for all spheres..do this once at beginning
//==============================================================
void box::SetInitialEvents()
{
  for (int i=0; i<N; i++)  // set all events to checks
    {
      event e(gtime, i, INF); 
      s[i].nextevent = e;
      h.insert(i);
    }
}


//==============================================================
// Finds next event for sphere i 
//==============================================================
event box::FindNextEvent(int i)
{
  double outgrowtime;
  outgrowtime = (.5/ngrids - (s[i].r+s[i].gr*gtime))/s[i].gr + gtime;

  event t = FindNextTransfer(i);
  event c = FindNextCollision(i);

  if ((outgrowtime < c.time)&&(outgrowtime < t.time)&&(ngrids>1))
    {
      event o = event(outgrowtime,i,INF-1);
      return o;
    }

  if ((c.time < t.time)&&(c.j == INF)) // next event is check at DBL infinity
    return c;
  else if (c.time < t.time) // next event is collision!
    {
      CollisionChecker(c); 
      return c; 
    }
  else // next event is transfer!
    return t;  
} 


//==============================================================
// Checks events of predicted collision partner to keep collisions
// symmetric
//==============================================================
void box::CollisionChecker(event c)
{
  int i = c.i;
  int j = c.j;
  event cj(c.time,j,i,c.v*(-1));

  // j should have NO event before collision with i!
  if (!(c.time  < s[j].nextevent.time))
    std::cout << i << " " <<  j << " error collchecker, s[j].nextevent.time= " << s[j].nextevent.time << " " << s[j].nextevent.j << ", c.time= " << c.time << std::endl;
  
  int k = s[j].nextevent.j; 
  if ((k < N) && (k!=i)) // j's next event was collision so give k a check
    s[k].nextevent.j = INF;
  
  // give collision cj to j
  s[j].nextevent = cj;
  h.upheap(h.index[j]);
}


//==============================================================
// Find next transfer for sphere i 
//==============================================================
event box::FindNextTransfer(int i)
{
  double ttime = dblINF;  
  int wallindex = INF;   // -(k+1) left wall, (k+1) right wall

  vector<DIM> xi = s[i].x + s[i].v*(gtime - s[i].lutime);
  vector<DIM> vi = s[i].v;

  for (int k=0; k<DIM; k++)
    {
      double newtime;
      if (vi[k]==0.) 
	newtime= dblINF;
      else if (vi[k]>0)  // will hit right wall, need to check if last wall
	{
	  if ((hardwallBC)&&(s[i].cell[k] == ngrids - 1))
	    newtime = ((double)(s[i].cell[k]+1)*SIZE/((double)(ngrids))
		       - (xi[k]+s[i].r+s[i].gr*gtime))/(vi[k]+s[i].gr);
	  else
	    newtime = ((double)(s[i].cell[k]+1)*SIZE/((double)(ngrids))
		       - xi[k])/(vi[k]);
	  
	  if (newtime<ttime)
	    {
	      wallindex = k+1;
	      ttime = newtime;
	    }
	}
      else if (vi[k]<0)  // will hit left wall
	{
	  if ((hardwallBC)&&(s[i].cell[k] == 0))
	    newtime = ((double)(s[i].cell[k])*SIZE/((double)(ngrids)) 
		       - (xi[k]-(s[i].r+s[i].gr*gtime)))/(vi[k]-s[i].gr);
	  else
	    newtime = ((double)(s[i].cell[k])*SIZE/((double)(ngrids)) 
		       - xi[k])/(vi[k]);
	  
	  if (newtime<ttime)
	    {
	      wallindex = -(k+1);
	      ttime = newtime;
	    }
	}
    }
  

  if (ttime < 0)
    ttime = 0;
  // make the event and return it
  event e = event(ttime+gtime,i,wallindex+DIM+N+1);
  return e;
}


//==============================================================
// Check all nearest neighbor cells for collision partners
//==============================================================
void box::ForAllNeighbors(int i, vector<DIM,int> vl, vector<DIM,int> vr,
			  neighbor& operation)
{
  vector<DIM,int> cell = s[i].cell;

  // now iterate through nearest neighbors
  vector<DIM, int> offset;          // nonnegative neighbor offset
  vector<DIM, int> pboffset;        // nearest image offset

   vector<DIM,int> grid;

   int ii ;

   grid=vl;
   while(1)
   {
     //if (vr[0] > 1)
     //std::cout << grid << "..." << cell+grid << "\n";
     for(int k=0; k<DIM; k++)
     {
        offset[k]=grid[k]+ngrids;  // do this so no negatives 	
        if (cell[k]+grid[k]<0) //out of bounds to left
          pboffset[k] = -1;
	else if (cell[k]+grid[k]>=ngrids) // out of bounds to right
	  pboffset[k] = 1;
        else
          pboffset[k] = 0;
     }     
     int j = cells.get((cell+offset)%ngrids);
     while(j!=-1)
       {
	 operation.Operation(j,pboffset);
	 j = binlist[j];
       }

     // A. Donev:     
     // This code makes this loop dimension-independent
     // It is basically a flattened-out loop nest of depth DIM
     for(ii=0;ii<DIM;ii++)
     {
       grid[ii] += 1;
       if(grid[ii]<=vr[ii]) break;
       grid[ii]=vl[ii];
     }
     if(ii>=DIM) break;
   }  
}


//==============================================================
// PredictCollision
//==============================================================
void box::PredictCollision(int i, int j, vector<DIM, int> pboffset, 
			     double& ctime, int& cpartner, 
			     vector<DIM, int>& cpartnerpboffset)
{
  double ctimej;
  
  if (i!=j)
    {	 
      ctimej = CalculateCollision(i,j,pboffset.Double())+gtime;
      
      if (ctimej < gtime)
	std::cout << "error in find collision ctimej < 0" << std::endl;
      
      if ((ctimej < ctime)&&(ctimej < s[j].nextevent.time))
	{
	  ctime = ctimej;
	  cpartner = j;
	  cpartnerpboffset = pboffset;
	}	
    }
}


//==============================================================
// Find next collision
//==============================================================
event box::FindNextCollision(int i)
{
  collision cc(i, this);
  
  vector<DIM, int> vl, vr;

  for (int k=0; k<DIM; k++)  // check all nearest neighbors
    {
      vl[k] = -1;
      vr[k] = 1;
    }
  
  ForAllNeighbors(i,vl,vr,cc);

  event e;
  if (cc.cpartner == i)  // found no collisions in neighboring cells
    {
      if (cc.ctime != dblINF)
	std::cout << "ctime != dblINF" << std::endl;
      e = event(dblINF,i,INF);  // give check at double INF
    }
  else
    e = event(cc.ctime,i,cc.cpartner,cc.cpartnerpboffset);

  return e;
}


//==============================================================
// Calculates collision time between i and image of j using quadratic formula
//==============================================================
double box::CalculateCollision(int i, int j, vector<DIM> pboffset)
{
  if ((hardwallBC)&&(pboffset.norm_squared() > 1E-12))
    {
      return dblINF;
    }
  else
    {      
      // calculate updated position and velocity of i and j
      vector<DIM> xi = s[i].x + s[i].v*(gtime - s[i].lutime);
      vector<DIM> vi = s[i].v;
      vector<DIM> xj = s[j].x + pboffset*SIZE + s[j].v*(gtime - s[j].lutime);
      vector<DIM> vj = s[j].v;
      
      double r_now_i = s[i].r + gtime*s[i].gr;
      double r_now_j = s[j].r + gtime*s[j].gr;
      
      double A,B,C;
      A = vector<DIM>::norm_squared(vi - vj) - (s[i].gr+s[j].gr)*(s[i].gr+s[j].gr);
      B = vector<DIM>::dot(xi - xj, vi - vj) - (r_now_i+r_now_j)*(s[i].gr+s[j].gr);
      C = vector<DIM>::norm_squared(xi - xj) - (r_now_i+r_now_j)*(r_now_i+r_now_j);

      if (C < -1E-12*(r_now_i+r_now_j))
	{
	  std::cout << "error, " << i << " and " << j << 
	    " are overlapping at time "<< gtime << std::cout;
	  std::cout << "A, B, C = "  << A << " " << " " << B << 
	    " " << " " << C <<  std::endl;
	  if (CheckSphereDiameters()>0)
	    std::cout << "a sphere has grown greater than unit cell" << 
	      std::endl;
	  else
	    std::cout << "unknown error" << std::cout;
	  exit(-1);
	}
      
      return QuadraticFormula(A, B, C);
    }  
}


//==============================================================
// Quadratic Formula ax^2 + bx + c = 0
//==============================================================
 double box::QuadraticFormula(double a, double b, double c)
{
  double x = dblINF;
  double xpos;
  double xneg;
  double det = b*b - a*c;

  if (c <= 0.)
    {
      if(b < 0.) // spheres already overlapping and approaching
	{
	  //std::cout << "spheres overlapping and approaching" << std::endl;
	  //std::cout << "# events= " << neventstot << std::endl;
	  x = 0.;	
	}
    }
  else if (det > -10.*DBL_EPSILON)
    {
      if (det < 0.)  // determinant can be very small for double roots
	det = 0.;    
      if (b < 0.)
	x = c/(-b + sqrt(det));
      else if ((a < 0.)&&(b > 0.))
	x = -(b + sqrt(det))/a;
      else
	x = dblINF;
    }
  return x;
}


//==============================================================
// Returns first event
//==============================================================
void box::ProcessEvent()
{  
  neventstot++;
  // Extract first event from heap
  int i = h.extractmax();   
  event e = s[i].nextevent; // current event
  event f;                  // replacement event

  if ((e.j>=0)&&(e.j<N))  // collision!
    {
      ncollisions++;
      //std::cout << "collision between " << e.i << " and " << e.j << " at time " << e.time << std::endl;
      Collision(e);
      f = FindNextEvent(i);
      s[i].nextevent = f;
      h.downheap(1);
      if (f.time < e.time)
	{
	  std::cout << "error, replacing event with < time" << std::endl;
	  exit(-1);
	}
      
      // make sure collision was symmetric and give j a check
      if ((s[e.j].nextevent.j != i)||(s[e.j].nextevent.time != gtime))
	{
	  std::cout << "error collisions not symmetric" << std::endl;
	  std::cout << "collision between " << e.i << " and " << e.j << " at time " << e.time << std::endl;
	  std::cout << "but " << e.j << " thinks it has " << s[e.j].nextevent.j<< " "  << s[e.j].nextevent.time << std::endl;
	  exit(-1);
	}
      else  // give j a check
	s[e.j].nextevent.j = INF;
    }
  else if (e.j==INF)      // check!  
    {
      nchecks++;
      //std::cout << "check for " << e.i << " at time " << e.time << std::endl;
      f = FindNextEvent(i);
      s[i].nextevent = f;
      h.downheap(1);
    }
  else if (e.j==INF-1)      // sphere outgrowing unit cell, decrease ngrids!
    {
      gtime = e.time;
      Synchronize(false);
      ngrids = ngrids - 1;
      std::cout << "need to reduce ngrids to " << ngrids << std::endl;   
      ChangeNgrids(ngrids);
      h.downheap(1);
    }
  else                    // transfer!
    {
      ntransfers++;
      //std::cout << "transfer for " << e.i << " at time " << e.time << std::endl;
      Transfer(e);
      f = FindNextEvent(i);
      s[i].nextevent = f;
      h.downheap(1);
      //r = FindNextEvent(i, e.j-N-DIM-1);
      //if (f.time <= e.time)
      if (f.time < e.time)
	{
	  std::cout << "error after transfer, replacing new event with < time" << " " << std::endl;
	  std::cout << std::setprecision(16) << "e.time= " << e.time << ", f.time= " << f.time << ", f.i= " << f.i << ", f.j= " << f.j << "e.i= " << e.i << ", e.j= " << e.j << std::endl;
	  std::cout << std::setprecision(16) << "difference= " << e.time - f.time << std::endl;
	  exit(-1);
	}
    }
}


//==============================================================
// Processes a collision
//=============================================================
void box::Collision(event e)
{
  double ctime = e.time;
  int i = e.i;
  int j = e.j;
  vector<DIM,int> v = e.v;  // virtual image
  gtime = ctime;

  // Update positions and cells of i and j to ctime
  s[i].x += s[i].v*(gtime-s[i].lutime);
  s[j].x += s[j].v*(gtime-s[j].lutime);

  // Check to see if a diameter apart
  double r_sum = s[i].r + s[j].r + (s[i].gr+s[j].gr)*gtime;
  double distance = vector<DIM>::norm_squared(s[i].x - s[j].x- v.Double()*SIZE) - r_sum*r_sum;
  if (distance*distance > 10.*DBL_EPSILON)
    std::cout << "overlap " << distance << std::endl;

  s[i].lutime = gtime;
  s[j].lutime = gtime;
  
  vector<DIM,double> vipar;          // parallel comp. vi
  vector<DIM,double> vjpar;          // parallel comp. vj
  vector<DIM,double> viperp;         // perpendicular comp. vi
  vector<DIM,double> vjperp;         // perpendicular comp. vj

  // make unit vector out of displacement vector
  vector<DIM,double> dhat;
  dhat = s[i].x - s[j].x - v.Double()*SIZE;  // using image of j!!
  double dhatmagnitude = sqrt(dhat.norm_squared());
  dhat /= dhatmagnitude;

  vipar = dhat*vector<DIM>::dot(s[i].v, dhat);
  vjpar = dhat*vector<DIM>::dot(s[j].v, dhat);
  viperp = s[i].v - vipar;
  vjperp = s[j].v - vjpar;

  s[i].v = vjpar + dhat*(s[i].gr+s[j].gr)*2 + viperp;
  s[j].v = vipar - dhat*(s[i].gr+s[j].gr)*2 + vjperp;

  // momentum exchange
  double xvelocity;   // exchanged velocity
  xvelocity = vector<DIM>::dot(s[i].v - s[j].v, dhat) - (s[i].gr+s[j].gr);
  xmomentum += xvelocity*dhatmagnitude*s[i].m*s[j].m*2/(s[i].m+s[j].m);
}


//==============================================================
// Transfer, takes care of boundary events too
//=============================================================
void box::Transfer(event e)
{
  gtime = e.time;
  int i = e.i;
  int j = e.j;
  int k=0;           // dimension perpendicular to wall it crosses
 
  // update position and lutime (velocity doesn't change)
  s[i].x += s[i].v*(gtime-s[i].lutime);
  s[i].lutime = gtime;

  vector<DIM,int> celli;  // new cell for i
  celli = s[i].cell;  // this is not redundant
  
  // update cell
  if (j>N+DIM+1)  // right wall
    {
      k = j-N-DIM-2;
      celli[k] = s[i].cell[k] + 1;

      if (hardwallBC)
	{
	  // if in right-most cell, reflect back
	  if (s[i].cell[k] == ngrids - 1)
	    s[i].v[k] = -s[i].v[k] - 4.*s[i].gr;
	  else
	    {
	      if (ngrids>1)
		UpdateCell(i, celli); 
	    }
	}
      else
	{
	  // if in right-most cell, translate x and cell
	  if (s[i].cell[k] == ngrids - 1)
	    {
	      s[i].x[k] -= SIZE;
	      celli[k] -= ngrids;
	    }
	  if (ngrids>1)
	    UpdateCell(i, celli); 
	}
    }
  else if (j<N+DIM+1)  // left wall
    {
      k = -j+N+DIM;
      celli[k] = s[i].cell[k] - 1;

      if (hardwallBC)
	{
	  // if in left-most cell, reflect back
	  if (s[i].cell[k] == 0)
	    s[i].v[k] = -s[i].v[k] + 4.*s[i].gr;
	  else
	    {
	      if (ngrids>1)
		UpdateCell(i, celli); 
	    }
	}
      else
	{
	  // if in left-most cell, translate x and cell
	  if (s[i].cell[k] == 0)
	    {
	      s[i].x[k] += SIZE;
	      celli[k] += ngrids;
	    }
	  if (ngrids>1)
	    UpdateCell(i, celli); 
	}
    }
  else
    std::cout << "error in Transfer" << std::endl;    
  
}


//==============================================================
// Updates cell of a sphere to time
//=============================================================
void box::UpdateCell(int i, vector<DIM,int>& celli)
{
  if (celli == s[i].cell)
    std::cout << "error in update cell..shouldn't be the same" << std::endl;
  
  // delete i from cell array at cell

  if (cells.get(s[i].cell) == i) 
    {
      if (binlist[i] == -1)
	cells.get(s[i].cell) = -1;
      else
	{
	  cells.get(s[i].cell) = binlist[i];
	  binlist[i] = -1;
	}
    }

  else if (cells.get(s[i].cell) == -1)
    {
      std::cout << "error " << i << " not in claimed cell UpdateCell" << std::endl;
      OutputCells();
    }

  else  // if no, find i in binlist
    {  
      int iterater = cells.get(s[i].cell);
      int pointer = iterater;
      while ((iterater != i)&&(iterater != -1))
	{
	  pointer = iterater;
	  iterater = binlist[iterater];
	}
      if (iterater == -1)  // got to end of list without finding i
	{
	  std::cout << "problem " << i << " wasn't in claimed, cell iterater = -1" << std::endl;
	  OutputCells();
	}
      else  // we found i!
	{
	  binlist[pointer] = binlist[i]; 
	  binlist[i] = -1;
	}	  
    } 

  // now add i to cell array at celli
  s[i].cell = celli;
  
  //first check to see if entry at celli
  if (cells.get(celli) == -1) //if yes, add i to cells gridfield
    cells.get(celli) = i;
  else  // if no, add i to right place in binlist
    {
      int iterater = cells.get(celli);  // now iterate through to end and add i
      int pointer = iterater;
      while (iterater != -1)  // find the end of the list
	{
	  pointer = iterater;
	  iterater = binlist[iterater];
	}
      binlist[pointer] = i;
      binlist[i] = -1; // redundant
    }
}


//==============================================================
// Output event heap...purely used for debugging
//==============================================================
void box::OutputEvents()
{
  h.print();
}


//==============================================================
// Output positions of spheres and their cells...purely used for debugging
//==============================================================
void box::OutputCells()
{
  for (int i=0; i<N; i++)
    std::cout << i << " " << s[i].x << " " << s[i].v << " " << s[i].cell << std::endl;
}


//==============================================================
// Update positions...purely for graphical display
//==============================================================
void box::TrackPositions()
{
  for (int i=0; i<N; i++)
    x[i] = s[i].x + s[i].v*(gtime-s[i].lutime);
}


//==============================================================
// Computes the total energy
//==============================================================
double box::Energy()
{
  double E=0;
  for (int i=0; i<N; i++)
    E += 0.5*s[i].m*s[i].v.norm_squared();

  return E/N;
}


//==============================================================
// Calculates the packing fraction
//==============================================================
double box::PackingFraction()
{
  double rfactor = 0.;
  for (int i=0; i<N; i++)
    {
      rfactor += pow(s[i].r + gtime*s[i].gr, DIM);
    }
  double v = (rfactor*pow(sqrt(PI), DIM))/(exp(lgamma(1.+((double)(DIM))/2.)));
  return v/(pow(SIZE, DIM));
}


//==============================================================
// Checks to make sure all sphere diameters are less than dimension
// of unit cell
//==============================================================
int box::CheckSphereDiameters()
{
  int offender = 0;
  for (int i=0; i<N; i++)
    {
      if (s[i].r*2 > 1./ngrids){
	offender = i;
	break;
      }
    }
  return offender;
}


//==============================================================
// Change ngrids
//==============================================================
void box::ChangeNgrids(int newngrids)
{
  cells.set_size(newngrids);
  cells.initialize(-1);      // initialize cells to -1
  for (int i=0; i<N; i++)    // initialize binlist to -1
    binlist[i] = -1;
  AssignCells();
  for (int i=0; i<N; i++)
    s[i].nextevent = event(0., i, INF); 
  Process(N); 
}	 


//==============================================================
// Calculates the optimal ngrids for the initial configuration
// and assumes that ngrids gets updated (reduced) as the packing
// proceeds
//==============================================================
int box::Optimalngrids2(double currentradius)
{
  return (int)(1./(2.*currentradius));
}


//==============================================================
// Calculates the optimal ngrids, assuming ngrids is not updated
// automatically and is very conservative
//==============================================================
int box::Optimalngrids()
{
  double maxr;

  maxr = pow(exp(lgamma(1.+((double)(DIM))/2.))*maxpf/
	     (N*(bidispersityfraction + (1.-bidispersityfraction)*
		 pow(bidispersityratio, DIM))), 
	     1./DIM)/sqrt(PI);

  return (int)(1./(2.*maxr));
}


//==============================================================
// Processes n events
//==============================================================
void box::Process(int n)
{
  double deltat = gtime;
  for (int i=0; i<n; i++)
    {
      ProcessEvent();
    }
  pf = PackingFraction();   // packing fraction
  deltat = gtime - deltat;
  double oldenergy = energy;
  energy = Energy();        // kinetic energy

  energychange = ((oldenergy - energy)/oldenergy)*100; // percent change in energy

  if (deltat != 0.)
    {
      pressure = 1+xmomentum/(2.*energy*N*deltat);
      collisionrate = ((double)(ncollisions))/deltat;
    }  

  // reset to 0
  ncollisions = 0;
  ntransfers = 0;
  nchecks = 0;
  xmomentum = 0.;
  ncycles++;
}


//==============================================================
// Prints statistics for n events
//==============================================================
void box::PrintStatistics()
{
  std::cout << "packing fraction = " << pf << std::endl;
  std::cout << "gtime = " << gtime << std::endl; 
  std::cout << "total time = " << rtime+gtime << std::endl;
  std::cout << "kinetic energy = " << energy << std::endl;
  std::cout << "total # events = " << neventstot << std::endl;
  std::cout << "# events = " << ncollisions+ntransfers+nchecks << ", # collisions = " << ncollisions << ", # transfers = " << ntransfers << ", # checks =" << nchecks << std::endl;
  std::cout << "growthrate = " << growthrate << std::endl;
  std::cout << "collisionrate = " << collisionrate << std::endl;
  std::cout << "reduced pressure = " << pressure << std::endl;
  std::cout << "-----------------" << std::endl;
}


//==============================================================
// Updates spheres to gtime, synchronizes, and can change growth rate
//==============================================================
void box::Synchronize(bool rescale)
{
  double vavg = sqrt(2.*M*energy);

  for (int i=0; i<N; i++)
    {
      s[i].x = s[i].x + s[i].v*(gtime-s[i].lutime);
      s[i].nextevent.time -= gtime;

      if (s[i].nextevent.time < 0.)
	std::cout << "error, event times negative after synchronization" << std::endl;
      if (rescale == true)   // give everyone checks
	{
	  s[i].nextevent = event(0., i, INF); 
	  s[i].v /= vavg;
	}
	  
      s[i].lutime = 0.;
      s[i].r += gtime*s[i].gr;   
    }

  //r += gtime*growthrate;       // r defined at gtime = 0
  rtime += gtime;
  gtime = 0.;
  
  if (rescale == true)
    Process(N);
}


//==============================================================
// Run time
//==============================================================
void box::RunTime()
{
  time(&end);
  std::cout << "run time = " << difftime(end, start) << std::endl;
}


//==============================================================
// Write configuration
//==============================================================
void box::WriteConfiguration(const char* wconfigfile)
{
  int count1;

  if (gtime != 0.)   // synchronize spheres if not currently synchronized
    Synchronize(false);
      
  std::ofstream output(wconfigfile);
  
  count1=0; // Number of spheres of first species
  for (int i=0; i<N; i++) {
   if(s[i].species==1) count1 += 1; }

  // make header
  output << DIM << "\n";
  output << N << " " << 2 << "\n";
  output << count1 << " " << N-count1 << "\n";
  output << std::setprecision(16) << 2*s[1].r << " " << 2*s[1].r*bidispersityratio << "\n";

 // Lattice vectors:
 for (int i=0; i<DIM; i++)
   for (int j=0; j<DIM; j++)
    {
      if (i==j)
	output << 1 << " ";
      else
	output << 0 << " ";
    }
 output << "\n";
 
 // Boundary conditions:
 for (int i=0; i<DIM; i++)   
   output << "T" << " ";  // T = periodic BC
 output << "\n";
  
 for (int i=0; i<N; i++)  // output radius and positions
   {
     for (int k=0; k<DIM; k++)
       output << std::setprecision(16) << s[i].x[k] << " ";
     output << std::setprecision(16) << 2*s[i].r << "";
     //output << std::setprecision(16) << s[i].gr << " ";
     //output << std::setprecision(16) << s[i].m << " ";
     output << "\n";
   }
 
 output.close();
}
```
