## Supplementary material for "Small subpopulations of β-cells do not drive islet oscillatory [Ca^2+^] dynamics via gap junction communication": S1 File: box.h

```
/* 
   Packing of hard spheres via molecular dynamics
   Developed by Monica Skoge, 2006, Princeton University
   Contact: Aleksandar Donev with questions
   This code may be used, modified and distributed freely.
   Please cite:
   
   "Packing Hyperspheres in High-Dimensional Euclidean Spaces"
   	M. Skoge, A. Donev, F. H. Stillinger and S. Torquato, 2006
	
   if you use these codes.	
*/

//-----------------------------------------------------------------------------
// Box maker
//---------------------------------------------------------------------------

#ifndef  BOX_H
#define  BOX_H


#include <vector>
#include <math.h>

#include "vector.h"
#include "grid_field.h"
#include "event.h"
#include "sphere.h"
#include "heap.h"


#define PI     3.141592653589793238462643
#define SIZE   1.0            // size of box
#define VOLUMESPHERE pow(PI,((double)(DIM))/2.)/exp(lgamma(1+((double)(DIM))/2.)) // volume prefactor for sphere
#define DBL_EPSILON  2.2204460492503131e-016 // smallest # such that 1.0+DBL_EPSILON!=1.0
#define M 1.0

//---------------------------------------------------------------------------
// Class neighbor
//---------------------------------------------------------------------------
class neighbor
{
 public:
  int i;
  
  neighbor(int i_i);

 public:
  virtual void Operation(int j, vector<DIM, int>& pboffset) = 0;
};


class box {
  
 public:
  
  // constructor and destructor
  box(int N_i, double r_i, double growthrate_i, double maxpf_i, 
      double bidispersityratio, double bidispersityfraction, 
      double massratio, int hardwallBC);
  ~box();

  // Creating configurations
  int Optimalngrids();
  int Optimalngrids2(double maxr);
  void CreateSpheres(double temp);
  void CreateSphere(int Ncurrent);   
  double Velocity(double temp);
  void VelocityGiver(double temp);  
  void SetInitialEvents();
  void RecreateSpheres(const char* filename, double temp);
  void ReadPositions(const char* filename);
  void AssignCells();
 
  // Predicting next event
  event FindNextEvent(int i);
  void CollisionChecker(event c);
  event FindNextTransfer(int i);
  event FindNextCollision(int i);
  void ForAllNeighbors(int, vector<DIM, int>, vector<DIM,int>, neighbor&);
  void PredictCollision(int i, int j, vector<DIM, int> pboffset, 
			double& ctime, int& cpartner, 
			vector<DIM, int>& cpartnerpboffset);
  double CalculateCollision(int i, int j, vector<DIM>  pboffset);
  double QuadraticFormula(double a, double b, double c);
  
  // Processing an event
  void Process(int n);
  void ProcessEvent();
  void Collision(event e);
  void Transfer(event e);
  void UpdateCell(int i, vector<DIM,int>& celli);
  void Synchronize(bool rescale);
  void ChangeNgrids(int newngrids);

  // Debugging
  void TrackPositions();
  void OutputEvents();
  void OutputCells();
  void GetInfo();  
  int CheckSphereDiameters();

  // Statistics
  double Energy();
  double PackingFraction();  
  void PrintStatistics();
  void RunTime();
  void WriteConfiguration(const char* wconfigfile);
  
  
  //variables

  const int N;                   // number of spheres
  
  int ngrids;                    // number of cells in one direction
  double maxpf;
  double growthrate;             // growth rate of the spheres
  double r;                      // radius, defined at gtime = 0
  double gtime;                  // this is global clock
  double rtime;                  // reset time, total time = rtime + gtime
  double collisionrate;          // average rate of collision between spheres
  double bidispersityratio;      // ratio of sphere radii
  double bidispersityfraction;   // fraction of smaller spheres
  double massratio;              // ratio of sphere masses
  int hardwallBC;                // =0 for periodic BC, =1 for hard wall
  

  // statistics
  double pressure;               // pressure
  double xmomentum;              // exchanged momentum
  double pf;                     // packing fraction
  double energy;                 // kinetic energy
  double energychange;
  int ncollisions;               // number of collisions
  int ntransfers;                // number of transfers
  int nchecks;                   // number of checks
  int neventstot;                // total number of events 
  int ncycles;                   // counts # cycles for output

  time_t start, error, end;      // run time of program

  // arrays
  sphere *s;                      // array of spheres
  grid_field<DIM, int> cells; // array that keeps track of spheres in each cell
  int *binlist;                   // linked-list for cells array
  heap h;                         // event heap
  vector<DIM> *x;                 // positions of spheres.used for graphics
};


//---------------------------------------------------------------------------
// Predicts collisions, inherits neighbor operation
//---------------------------------------------------------------------------
class collision : public neighbor 
{
 public:
  
  box *b; 
  double ctime;
  int cpartner;
  vector<DIM,int> cpartnerpboffset;

 public:
  collision(int i_i, box *b);

  virtual void Operation(int j, vector<DIM, int>& pboffset);
};

#endif 
```
