## Supplementary material for "Small subpopulations of β-cells do not drive islet oscillatory [Ca^2+^] dynamics via gap junction communication": S1 File: ChR2Class.h

```
using namespace std;
#include <cmath>
#include <iostream>
//const double Gd1=0.12;
const double Gd1=0.3257;
const double Gd2=0.11;
//const double Gd2=0.025;
//const double e12=0.0394;
const double e12dark=0.0127;
//const double e21=0.0432;
const double e21dark=0.0218;
const double e1=1.5057;
const double e2=0.645;
const double F0=0.1;
const double dtOld=0;
const double dt=0.1;
const double Gr=9.6694E-4;
//const double Gr=2.56
const double tChR2=1.3;
const double tEnd=10;
const double gO1=.4;
const double gO2=0.04;
const double gammaC=0.1;
const double wavelength=458;
class ChR2Current
{
public:
double O11;
double O21;
double C11;
double C21;

double Ga1;
double Ga2;
double dC1;
double dC2;
double dO1;
double dO2;


void calc_derivs(double,double,double,double,double,double,double,double,double);
};

void ChR2Current::calc_derivs(double O1,double O2,double C1,double C2,double tStart,double tEnd,double t0,double period,double phi)
{
    
   double e12=e12dark+0.005*log(1+(phi/0.024));
   double e21=e21dark+0.004*log(1+(phi/0.024));
   
   double pFlux=(1000*phi/(1E9*(1.98644E-25)/(wavelength)));
   double F0=(1.2E-20)*pFlux/(1000*1.3);
   double dutyCycle=2;
   double t=t0-tStart;
   double tMod=period/dutyCycle - (fmod(t,period));
   //Light transition
   double tCurr=t-(floor(t/period)*period);

   if(t0<tEnd && tMod>=0)
{
 Ga1=e1*F0*(1-exp(-(tCurr)/tChR2));
 Ga2=e2*F0*(1-exp(-(tCurr)/tChR2));


 dO1=Ga1*C1-(Gd1+e12)*O1+e21*O2;
 dO2=Ga2*C2+e12*O1-(Gd2+e21)*O2;
 dC2=Gd2*O2-(Ga2+Gr)*C2;
 dC1=Gr*C2+Gd1*O1-Ga1*C1;


 O11=O1;
 O21=O2;
 C11=C1;
 C21=C2;

}
   //Dark transition
else
{
	
 //Ga1=e1*F0*(exp(-(tCurr-period/dutyCycle)/tChR2)-exp(-(tCurr)/tChR2));
 //Ga2=e2*F0*(exp(-(tCurr-period/dutyCycle)/tChR2)-exp(-(tCurr)/tChR2));
 Ga1=e1*F0*(exp(-(tCurr)/tChR2)-exp(-(tCurr)/tChR2));
 Ga2=e2*F0*(exp(-(tCurr)/tChR2)-exp(-(tCurr)/tChR2));
 if (t0>=tEnd)
 {
	 //tCurr=t-tEnd;
	 //Ga1=e1*F0*(exp(-(tCurr+period/dutyCycle)/tChR2)-exp(-(tCurr)/tChR2));
	 //Ga2=e2*F0*(exp(-(tCurr+period/dutyCycle)/tChR2)-exp(-(tCurr)/tChR2));
	 //Ga1=e1*F0*(exp(-(t0-tEnd)/tChR2));
	 //Ga2=e2*F0*(exp(-(t0-tEnd)/tChR2));
	 Ga1=0;
	 Ga2=0;
 }


 dO1=-(Gd1+e12dark)*O1+e21dark*O2;
 dO2=e12dark*O1-(Gd2+e21dark)*O2;
 dC2=Gd2*O2-(Ga2+Gr)*C2;
 dC1=Gr*C2+Gd1*O1-Ga1*C1;
 
 O11=O1;
 O21=O2;
 C11=C1;
 C21=C2;
        
    
       
}
  
}
```
