## Supplementary material for "Small subpopulations of β-cells do not drive islet oscillatory [Ca^2+^] dynamics via gap junction communication": S1 File: event.C

```
#include "event.h" 

//==============================================================
//==============================================================
//  Class Event 
//==============================================================
//==============================================================


//==============================================================
// Constructor
//==============================================================
event::event(double time_i, int i_i, int j_i, vector<DIM,int> v_i):
  time(time_i),
  i(i_i),
  j(j_i),
  v(v_i)
{
}

event::event(double time_i, int i_i, int j_i):
  time(time_i),
  i(i_i),
  j(j_i)
{
}

event::event(const event& e)
{
  time = e.time;
  i = e.i;
  j = e.j;
  v = e.v;
}

event::event()
{
}

  
//==============================================================
// Destructor
//==============================================================
event::~event() 
{
}

void event::erase()
{
  time = dblINF;
  i = 0;
  j = 0;
}

bool event::operator<(const event& e) const
{
  return e.time < time;
}

bool event::operator>(const event& e) const
{
  return e.time > time;
}

 
```
