## Supplementary material for "Small subpopulations of β-cells do not drive islet oscillatory [Ca^2+^] dynamics via gap junction communication": S1 File: event.h

```
#ifndef  EVENT_H
#define  EVENT_H

#include "vector.h"
#define INF    100000000
#define dblINF 100000000.

class event {

 public:

  // constructor and destructor
  event(double time_i, int i_i, int j_i, vector<DIM,int> v_i);
  event(double time_i, int i_i, int j_i);
  event(const event& e);
  event();
  
  ~event();

  bool operator<(const event&) const;
  bool operator>(const event&) const;
  void erase();

 //variables


  double time;             // time of next collision
  int i;        // collision partner with lower number
  int j;        // collision partner with higher number
  vector<DIM,int> v;        // virtual image

  /* 0<=j<=N                binary collision between i and j
     j=N+DIM+1+x            transfer where x=-(k+1) for left wall
                            and x=k+1 for right wall
     j=INF                 both check after event that did not altered motion of                           i and check after event that altered motion of i, i.e                           rescaling of velocities. I currently don't see need t                           o separate the two

     j=-1                  check after collision

     Virtual identifiers as scalars...I think bad idea, but here's my work
     there will be easily fixed problems if DIM>=10
     -x<=v<=x               x=k+1, image in k direction
     v=xy                   x,y
      =-xy                  -x,y
      =-yx                  x,-y
      =yx                   -x,-y
     v=xyz                  x,y,z
      =-xyz                 -x,y,z
      =-yxz                 x,-y,z
      =-zxy                 x,y,-z
      =zyx                  -x,-y,z
      =xzy                  x,-y,-z
      =yzx                  -x,y,-z
      =-zyx                 -x,-y,-z
  */      

};

#endif 
```
