## Supplementary material for "Small subpopulations of β-cells do not drive islet oscillatory [Ca^2+^] dynamics via gap junction communication": S1 File: GenerateSphere.cpp

```
#include <iostream>
#include <string>
#include <fstream>
#include <ostream>
#include <cstdlib>
#include <stdio.h>
#include <cmath>
#include <string>
using namespace std;


///////////////Set the amount of cells you want here/////////////////
int cellNumber=1000;
struct sphereData
{
	//Loads the initial sphere packing of 4000 Cells
double data[4000][4];
double XYZdata[4000][4];

double dist[4000];
void loadSphereInfo( )
{
	//Ignores some garbage data from the write file
int numCells=4000;
ifstream file("write.dat");
file.ignore(256,'\n'); 
file.ignore(256,'\n'); 
file.ignore(256,'\n'); 
file.ignore(256,'\n'); 
file.ignore(256,'\n'); 
file.ignore(256,'\n'); 
for (int f=0;f<4000;f++)
{
for (int g=0;g<4;g++)
{
file>>data[f][g];
}
}
for (int i=0;i<4000;i++)
{
	//Pulls in the XYZ coordinates of all the cells
double x=data[i][0];
double y=data[i][1];
double z=data[i][2];
   //Calculates the distance between each cell and the center of the islet
dist[i]=sqrt(pow(x-0.5,2)+pow(y-0.5,2)+pow(z-0.5,2));
}
// 
double radius=0.025;
int count=0;
////////////////Starts from the center of the packing and takes only the specified number of cells/////////////////
while (numCells!=cellNumber)
{
count=0;
for (int i=0;i<4000;i++)
{
if (dist[i]<=radius)
{
count++;
}
}
if (count<cellNumber)
{
radius=radius+radius/cellNumber;
}
if (count>cellNumber)
{
radius=radius-radius/cellNumber;
}
numCells=count;
//cout<<numCells<<endl;
}

count=0;
for (int i=0;i<4000;i++)
{
if(dist[i]<=radius)
{
XYZdata[count][0]=data[i][0];
XYZdata[count][1]=data[i][1];
XYZdata[count][2]=data[i][2];
XYZdata[count][3]=data[i][3];
count++;
}
}
}
/////////////////////////////////////////////////////////////////////////////////////////////////////////
};

// Structure to calculate the Nearest Neihbors of each cell
struct sphereCalc
{
public:
const int numCols;
double NN[4000][15];
sphereCalc(const int numCols) : numCols(numCols) {}
void sphereNN(double XYZ[4000][4])
{
double x;
double y;
double z;
double x1;
double y1;
double z1;
double dist;
double minDist=1;
int jMin=0;
//Sets up a blank row for each cell
for (int i=0;i<numCols;i++)
{
for (int j=0;j<15;j++)
{
NN[i][j]=-1;
}

}
//Run through each cell and calculate distance between every other cell. If this distance is less than a specified critera (related to the radius), call it a neighboor
for (int i=0;i<numCols;i++)
{
x=XYZ[i][0];
y=XYZ[i][1];
z=XYZ[i][2];
int count=0;
for (int j=0;j<numCols;j++)
{
x1=XYZ[j][0];
y1=XYZ[j][1];
z1=XYZ[j][2];
if (j!=i)
{
dist=sqrt(pow((x-x1),2)+pow((y-y1),2)+pow((z-z1),2));
if (dist<minDist)
{
minDist=dist;
jMin=j;
}
//Nearest Neighboor distance criteria.
if (dist<0.072)
{
NN[i][count]=j;
count=count+1;
}
}
}
if (count==0)
{
NN[i][0]=jMin;
}
}


}

};


// Calling the main file to 
int main(int argc, char** argv)
{

string NNname;
string cNum;
char *pEnd;
ifstream f(argv[1]);
if (f.is_open())
getline(f,NNname);
getline(f,cNum);
char *fileName = (char*)NNname.c_str();
cellNumber=strtod(cNum.c_str(),&pEnd);
cout<<NNname<<endl;
sphereData myData;
myData.loadSphereInfo();
remove(fileName);
sphereCalc myCalc(cellNumber);
remove("XYZpos.txt");
myCalc.sphereNN(myData.XYZdata);
ofstream file;
file.open("XYZpos.txt",ios::app);
cout<<cellNumber<<endl;
for (int i=0;i<cellNumber;i++)
{
for (int j=0;j<3;j++)
{
file<<myData.XYZdata[i][j]<<' ';
}
file<<endl;
}

file.close();
std::ofstream outfile;
outfile.open(fileName,ios::app);
for (int i=0;i<cellNumber;i++)
{
for (int j=0;j<15;j++)
{
cout<<myCalc.NN[i][j]<<' ';
outfile<<myCalc.NN[i][j]<<' ';
}
cout<<endl;
outfile<<endl;
}
outfile.close();
return 0;
}
```
