## Supplementary material for "Small subpopulations of β-cells do not drive islet oscillatory [Ca^2+^] dynamics via gap junction communication": S1 File: grid_field.h

```
/* 
   Packing of hard spheres via molecular dynamics
   Developed by Monica Skoge, 2006, Princeton University
   Contact: Aleksandar Donev with questions
   This code may be used, modified and distributed freely.
   Please cite:
   
   "Packing Hyperspheres in High-Dimensional Euclidean Spaces"
   	M. Skoge, A. Donev, F. H. Stillinger and S. Torquato, 2006
	
   if you use these codes.	
*/


#ifndef GRID_FIELD_H
#define GRID_FIELD_H

#include "vector.h"

// ======================================================================
// grid_field 
// ======================================================================

// A field of V-vectors on a D dimensional manifold

template<int D, class T>
class grid_field {
  
 public:
  int elements;

 private:
  T* f;
  vector<D, int> size;           // number of grid points for each dimension
  vector<D, int> offset;
 
 public:

  grid_field();
  grid_field(const vector<D, int>&);
  ~grid_field();

  T& get(const vector<D, int>&);

  vector<D, int> get_size() const;
  void set_size(const vector<D, int>&);
  void set_size(const int);
  void initialize(const int i);
};


// grid_field
// ~~~~~~~~~~~~
template<int D, class T>
grid_field<D, T>::grid_field()
  : f(0), elements(0)
{
}


// grid_field
// ~~~~~~~~~~~~
template<int D, class T>
grid_field<D, T>::grid_field(const vector<D, int>& s)
  : f(0)
{
  set_size(s);
}


// ~grid_field
// ~~~~~~~~~~~~~
template <int D, class T>
grid_field<D, T>::~grid_field()
{
  if(f != 0)
    delete[] f;
}


// get_size
// ~~~~~~~~
template<int D, class T>
inline vector<D, int> grid_field<D, T>::get_size() const
{
  return size;
}


// set_size
// ~~~~~~~~
template<int D, class T>
void grid_field<D, T>::set_size(const vector<D, int>& s)
{
  if(f != 0)
    delete[] f;

  size = s;

  elements = 1;
  for(int i=0; i<D; i++) {
    offset[i] = elements;
    elements *= size.x[i];
  }

  f = new T[elements];
}


// set_size
// ~~~~~~~~
template<int D, class T>
void grid_field<D, T>::set_size(const int s)
{
  vector<D, int> square;

  for(int k=0; k<D; k++)
    square[k] = s;

  set_size(square);
}


// get
// ~~~
template<int D, class T>
inline T& grid_field<D, T>::get(const vector<D, int>& pos)
{
  int p=0;
  for(int i=0; i<D; i++)
    p += pos.x[i]*offset[i];

  return f[p];
}


// initialize
// ~~~
template<int D, class T>
void grid_field<D, T>::initialize(const int value)
{
  for(int i=0; i<elements; i++)
    f[i] = value;
}


#endif
```
