## Supplementary material for "Small subpopulations of β-cells do not drive islet oscillatory [Ca^2+^] dynamics via gap junction communication": S1 File: heap.C

```
#include "heap.h"
#include "event.h"
#include <iostream>
#include "box.h"


//==============================================================
//==============================================================
//  Class heap: Event heap used in hard spheres calculation
//==============================================================
//==============================================================


//==============================================================
// Constructor
//==============================================================
heap::heap(int maxsize_i):
  maxsize(maxsize_i)
{
  a = new int[maxsize];
  index = new int[maxsize];
  
  N = 0;   // current number of events in heap
}


//==============================================================
// Constructor
//==============================================================
heap::heap(const heap &h)
{
  maxsize = h.maxsize;
  a = h.a;
  index = h.index;
  N = h.N;                // current number of events in heap
  s = h.s;
}


//==============================================================
// Destructor
//==============================================================
heap::~heap()
{
  delete[] a;
  delete[] index;
}

//==============================================================
// Upheap
//==============================================================
void heap::upheap(int k)
{
  int i = a[k];

  while ((k>1)&&(s[a[k/2]].nextevent.time > s[i].nextevent.time))
    {
      a[k] = a[k/2];
      index[a[k/2]] = k;
      k = k/2;
    }
  a[k] = i;
  index[i] = k;
}

//==============================================================
// Downheap
//==============================================================
void heap::downheap(int k)
{
  int j;
  int i = a[k];
  
  while(k <= N/2)
    {
      j = k+k;
      if ((j < N)&&(s[a[j]].nextevent.time > s[a[j+1]].nextevent.time))
	j++;
      if (s[i].nextevent.time <= s[a[j]].nextevent.time)
	break;
      a[k] = a[j];
      index[a[j]] = k;
      k = j;
    }
  a[k] = i;
  index[i] = k;
}

//==============================================================
// Insert
//==============================================================
void heap::insert(int i)
{
  if (N >= maxsize)
    std::cout << "error, N >= maxsize, cannot insert another event" << std::endl;
  else
    {
      N++;
      a[N] = i;
      index[i] = N;
      upheap(N);
    }  
}


//==============================================================
// Extract max
//==============================================================
int heap::extractmax()
{
  return a[1];
}

/*
//==============================================================
// Replace
//==============================================================
void heap::replace(int i)
{
  a[1] = i;
  s[i].nextevent = t;

  if (!(e.time > s[a[1]].nextevent.time))
    {
      if (!(s[a[1]].nextevent.j == INF))// must be check i'm changing to coll. at same time
	std::cout << "error replaced non check with earlier time" << std::endl;
      a[1] = i;
      index[i] = 1;
    }
  else
    {    
      a[0] = i;
      downheap(0);
    }
}
*/

//==============================================================
// Search
//==============================================================
int heap::search(int j)
{
  for (int k=1; k<=N; k++)
    {
      if (a[k] == j)
	return k;
    }
  return -1;
}

/*
//==============================================================
// Change
//==============================================================
void heap::change(int i)
{
  int iindex = index[i];
  
  if (s[i].nextevent.time == s[a[iindex]].nextevent.time)  
    std::cout << "error changing an event to an equal time" << std::endl;
  else if (s[i].nextevent.time > s[a[iindex]].nextevent.time)
    std::cout << "error changing an event to a greater time" << std::endl;

  a[iindex] = i;
  upheap(iindex);
}
*/

//==============================================================
// Print
//==============================================================
void heap::print()
{
  for (int k=1; k<=N; k++)
    std::cout << k << " " << a[k] << " " << s[a[k]].nextevent.j << " " << s[a[k]].nextevent.time << std::endl;
}


//==============================================================
// Check index array
//==============================================================
void heap::checkindex()
{
  for (int k=1; k<=N; k++)
    std::cout << k << " " << a[k] << " " << index[a[k]] << std::endl;
}
```
