## Supplementary material for "Small subpopulations of β-cells do not drive islet oscillatory [Ca^2+^] dynamics via gap junction communication": S1 File: heap.h

```
//---------------------------------------------------------------------------
// Event heap maker
//---------------------------------------------------------------------------

#ifndef  HEAP_H
#define  HEAP_H

#include "event.h"
#include "sphere.h"

class heap {

 public:

  // constructor and destructor
  heap(int maxsize);
  heap(const heap &h);
  ~heap();

  // variables
  int maxsize;   // max allowed number of events
  int N;         // current number of events
  int *a;
  sphere *s;
  int *index;     // array of indices for each sphere
  //event minevent;


  // functions which operate on a binary heap
  
  void upheap(int k);
  void downheap(int k);
  void insert(int i);
  void replace(int i);
  int search(int j);
  void change(int i); 
  int extractmax();
  void print();
  void checkindex();
  
};
#endif 
```
