## Supplementary material for "Small subpopulations of β-cells do not drive islet oscillatory [Ca^2+^] dynamics via gap junction communication": S1 File: InCircle.h

```
#ifndef INCIRCLE_H
#define INCIRCLE_H

#endif // INCIRCLE_H


struct CircleCoords
{
  int InCircleBool[cellNumber];
double X;
double Y;
double Z;

  for (int i=0;i<1000;i++)
  {
  X=cellPos[i][0];
  Y=cellPos[i][1];
  Z=cellPos[i][2];


  }
};
```
