## Supplementary material for "Small subpopulations of β-cells do not drive islet oscillatory [Ca^2+^] dynamics via gap junction communication": S1 File: MainFile.cpp

```
#include <stdio.h>
#include <iostream>
#include <boost/numeric/odeint.hpp>
#include <boost/multi_array.hpp>
#include <boost/numeric/ublas/matrix.hpp>
#include <boost/numeric/ublas/io.hpp>
#include <cassert>
#include <cmath>
#include <ostream>
#include <fstream>
#include <vector>
#include <boost/random/detail/config.hpp>
#include <boost/random.hpp>
#include <boost/random/normal_distribution.hpp>
#include <boost/fusion/container/vector.hpp>
#include <boost/fusion/include/vector.hpp>
#include <boost/fusion/container/vector/vector_fwd.hpp>
#include <boost/fusion/include/vector_fwd.hpp>
#include <omp.h>
#include <string>
#include <algorithm>
#include <boost/math/distributions/skew_normal.hpp>
#include "BetaCell.h"

int main( int argc , char **argv )
{

//while(ChR2CurrentPulse(0,period1,1000,Frequency)>ChR2CurrentPulse(0,delt,1000,Frequency))
//{
//delt=delt+0.001;
//}


const int lengthNum=cellNumber*30;


  // Get the number of processors this job is using:
int varNumber;

  // Get the rank of the processor this thread is running on.  (Each
  // processor has a unique rank.)


vector_type x1(cellNumber*30);
vector_type dxdt(cellNumber*30);
double* varAr=new double[lengthNum];
ifstream myFile ("vars5exo.txt");
if (myFile.is_open())
{
for (int i=0;i<30*cellNumber;i++)
{
myFile>>varAr[i];
}

myFile.close();
}


ifstream infoFile(argv[1]);

string NNfileLine;

getline(infoFile,NNfileLine);

char *NNfileName=(char*)NNfileLine.c_str();

ifstream NNfile(NNfileName);
for (int i=0;i<cellNumber;i++)
{
for (int j=0;j<15;j++)
{
NNfile>>NN[i][j];
}
}

ifstream cFile("XYZpos.txt");
if(cFile.is_open())
{
for (int i=0; i<cellNumber;i++)
{
for (int j=0;j<3;j++)
{
cFile>>cellPos[i][j];
//cout<<cellPos[i][j];
}
//cout<<endl;
}
}

string potentialNameLine;
string timeInfoLine;
string calciumInfoLine;
string mutantLine;
string sodiumInfoLine;
string potassiumInfoLine;
string caerInfoLine;
string atpInfoLine;
string adpInfoLine;
string ikatpInfoLine;
string IRPInfoLine;
string PPInfoLine;
string DPInfoLine;
string FIPInfoLine;
string RIPInfoLine;
string CapInfoLine;
string NoiseInfoLine;
    
getline(infoFile,potentialNameLine);
getline(infoFile,sodiumInfoLine);
getline(infoFile,potassiumInfoLine);
getline(infoFile,calciumInfoLine);
getline(infoFile,caerInfoLine);
getline(infoFile,atpInfoLine);
getline(infoFile,adpInfoLine);
getline(infoFile,IRPInfoLine);
getline(infoFile,PPInfoLine);
getline(infoFile,DPInfoLine);
getline(infoFile,FIPInfoLine);
getline(infoFile,RIPInfoLine);
getline(infoFile,CapInfoLine);
getline(infoFile,timeInfoLine);
getline(infoFile,NoiseInfoLine);
getline(infoFile,mutantLine);

char* pEnd;


pMutant=strtod(mutantLine.c_str(),&pEnd);
potentialname=(char*)potentialNameLine.c_str();
calciuminfo=(char*)calciumInfoLine.c_str();
sodiuminfo=(char*)sodiumInfoLine.c_str();
potassiuminfo=(char*)potassiumInfoLine.c_str();
timename=(char*)timeInfoLine.c_str();
caerinfo=(char*)caerInfoLine.c_str();
atpinfo=(char*)atpInfoLine.c_str();
adpinfo=(char*)adpInfoLine.c_str();
IRPinfo=(char*)IRPInfoLine.c_str();
PPinfo=(char*)PPInfoLine.c_str();
DPinfo=(char*)DPInfoLine.c_str();
FIPinfo=(char*)FIPInfoLine.c_str();
RIPinfo=(char*)RIPInfoLine.c_str();
Capinfo=(char*)CapInfoLine.c_str();
Noiseinfo=(char*)NoiseInfoLine.c_str();
    
remove("O1.txt");
remove("O2.txt");
remove("C1.txt");
remove("C2.txt");


O1info="O1.txt";
O2info="O2.txt";
C1info="C1.txt";
C2info="C2.txt";


if( remove(ikatpinfo) !=0)
	perror("Error Delting");
	else
		puts("Success");

if( remove(potentialname ) != 0 )
    perror( "Error deleting file" );
  else
    puts( "File successfully deleted" );

 if( remove( timename) != 0 )
    perror( "Error deleting file" );
  else
    puts( "File successfully deleted" );

if( remove( cellinfo ) != 0 )
    perror( "Error deleting file" );
  else
    puts( "File successfully deleted" );

if (remove(calciuminfo)!=0)
	perror("Error Deleting File");
else
	puts("File Deleted correctly");

if(remove(sodiuminfo)!=0)
perror("Error Deleting File");
else
puts("Deleted Correctly");

if(remove(potassiuminfo)!=0)
perror("Error Deleting File");
else
puts("Deleted Correctly");

if(remove(caerinfo)!=0)
perror("Error Deleting File");
else
puts("Deleted Correctly");

if(remove(atpinfo)!=0)
perror("Error Deleting File");
else
puts("Deleted Correctly");

if(remove(adpinfo)!=0)
perror("Error Deleting File");
else
puts("Deleted Correctly");
    
    if(remove(PPinfo)!=0)
        perror("Error Deleting File");
    else
        puts("Deleted Correctly");
    
    if(remove(IRPinfo)!=0)
        perror("Error Deleting File");
    else
        puts("Deleted Correctly");
    
    if(remove(DPinfo)!=0)
        perror("Error Deleting File");
    else
        puts("Deleted Correctly");
    
    if(remove(FIPinfo)!=0)
        perror("Error Deleting File");
    else
        puts("Deleted Correctly");
    
    if(remove(RIPinfo)!=0)
        perror("Error Deleting File");
    else
        puts("Deleted Correctly");
    
    if(remove(Capinfo)!=0)
        perror("Error Deleting File");
    else
        puts("Deleted Correctly");
    if(remove(Noiseinfo)!=0)
        perror("Error Deleting File");
    else
        puts("Deleted Correctly");

int count=0;
for ( int i=0;i<cellNumber*30;i++)
{
x1[i]=varAr[i];
}


ofstream excitationFile;
remove("ChR2Cells.txt");
excitationFile.open("ChR2Cells.txt");

ifstream randomVar("RandomVars.txt");
double RandomVarMat[cellNumber][12];
for(int i=0;i<cellNumber;i++)
{
for(int j=0;j<12;j++)
{
randomVar>>RandomVarMat[i][j];
//cout<<RandomVarMat[i][j]<<' ';
}
//cout<<' '<<endl;
}

//Setting cellular properties
for(int i=0;i<cellNumber;i++)
{
gKATPar[i]=RandomVarMat[i][0];
gCoup[i]=RandomVarMat[i][1];
gKtoar[i]=RandomVarMat[i][2];
PCaERar[i]=RandomVarMat[i][3];
gKCaBKar[i]=RandomVarMat[i][4];
PNACAar[i]=RandomVarMat[i][5];
Prelar[i]=RandomVarMat[i][6];
Popar[i]=RandomVarMat[i][7];
ATPar[i]=RandomVarMat[i][8];
KRev[i]=RandomVarMat[i][9];
gChR2[i]=RandomVarMat[i][10];
RandomSeed[i]=RandomVarMat[i][11];
}


//integrate_adaptive(make_controlled(1.0,1.0,stepper_type()),stiff_system(mynode),x1,0.0,100000.00,0.1);
//doIntegrate(mynode,x1,0.1,1000.0,1.
tMax=900000.00;
totalnodes=1;
//omp_set_num_threads(1);


/*#pragma omp parallel shared(totalnodes)
{
const int mynode=omp_get_thread_num();

integrate_const(euler<vector_type>(),stiff_system(mynode),x1,0.0,tMax,0.2);

}
*/


BetaSolver(x1,dxdt,0.1);


    return 0;
}
```
