## Supplementary material for "Small subpopulations of β-cells do not drive islet oscillatory [Ca^2+^] dynamics via gap junction communication": S1 File: my_vector.h

```
#ifndef VECTOR_H
#define VECTOR_H

#include <iostream>
#include <fstream>

#define DIM 3

template <int D, typename T=double>
class vector {
  
  public:
  T x[D];
  
  public:
  
  vector();
  vector(const T[D]);           
  vector(const vector&);
  ~vector();

  vector<D, T>& operator+=(const vector<D, T>&);
  vector<D, T>& operator-=(const vector<D, T>&);
  vector<D, T>& operator*=(const T);
  vector<D, T>& operator/=(const T);
  vector<D, T> operator+(const vector<D, T>&) const;
  vector<D, T> operator-(const vector<D, T>&) const;
  vector<D, T> operator*(const T) const;
  vector<D, T> operator/(const T) const;
  vector<D, T> operator%(const T) const;
  bool operator==(const vector<D, T> &a) const;

  vector<D, int> integer() const;
  vector<D, double> Double() const;
  static vector<D, int> integer(const vector<D, T>&); 
  static vector<D, double> Double(const vector<D, T>&); 
  
  T& operator[](const unsigned int);
 
  double dot(const vector<D, T>&) const;
  static double dot(const vector<D, T>&, const vector<D, T>&);
  
  double norm_squared() const;
  static double norm_squared(const vector<D, T>&);

  void read(std::ifstream&);
  void write(std::ofstream&) const;
};


template <int D, typename T>
std::ostream& operator<<(std::ostream&, const vector<D, T>&);


// constructor
// ~~~~~~~~~~~
template <int D, typename T>
vector<D, T>::vector()
{
  for(int k=0; k<D; k++)
    x[k] = 0;
}

template <int D, typename T>
vector<D, T>::vector(const T x_i[D])
{
  for(int k=0; k<D; k++)
    x[k] = x_i[k];
}

template <int D, typename T>
vector<D, T>::vector(const vector<D, T> &v)
{
  for(int k=0; k<D; k++)
    x[k] = v.x[k];
}


// destructor
// ~~~~~~~~~~
template <int D, typename T>
vector<D, T>::~vector()
{
}


// +=
// ~~
template <int D, typename T>
inline vector<D, T>& vector<D, T>::operator+=(const vector<D, T> &v)
{
  for(int k=0; k<D; k++)
    x[k] += v.x[k];

  return *this;
}


// -=
// ~~
template <int D, typename T>
inline vector<D, T>& vector<D, T>::operator-=(const vector<D, T> &v)
{
  for(int k=0; k<D; k++)
    x[k] -= v.x[k];

  return *this;
}


// *=
// ~~
template <int D, typename T>
inline vector<D, T>& vector<D, T>::operator*=(const T s)
{
  for(int k=0; k<D; k++)
    x[k] *= s;

  return *this;
}

// /=
// ~~
template <int D, typename T>
inline vector<D, T>& vector<D, T>::operator/=(const T s)
{
  for(int k=0; k<D; k++)
    x[k] /= s;

  return *this;
}


// +
// ~ 
template <int D, typename T>
inline vector<D, T> vector<D, T>::operator+(const vector<D, T> &a) const
{
  vector<D, T> c;
  
  for(int k=0; k<D; k++)
    c.x[k] = x[k] + a.x[k];

  return c;
}


// -
// ~
template <int D, typename T>
inline vector<D, T> vector<D, T>::operator-(const vector<D, T> &a) const
{
  vector<D, T> c;

  for(int k=0; k<D; k++)
    c[k] = x[k] - a.x[k];

  return c;
}


// *
// ~
template <int D, typename T>
inline vector<D, T> vector<D, T>::operator*(const T s) const
{
  vector<D, T> c;
  
  for(int k=0; k<D; k++)
    c[k] = x[k] * s;

  return c;
}


// /
// ~
template <int D, typename T>
inline vector<D, T> vector<D, T>::operator/(const T s) const
{
  vector<D, T> c;

  for(int k=0; k<D; k++)
    c[k] = x[k] / s;

  return c;
}


// ==
// ~
template <int D, typename T>
inline bool vector<D, T>::operator==(const vector<D, T> &a) const
{
  for(int k=0; k<D; k++)
    {
      if (!(x[k]==a.x[k]))
	return false;
    }
  return true;
}


// %
// ~
template <int D, typename T>
inline vector<D, T> vector<D, T>::operator%(const T s) const
{
  vector<D, T> c;

  for(int k=0; k<D; k++)
    c[k] = x[k] % s;

  return c;
}


// integer
// ~~~~~~~
template <int D, typename T>
inline vector<D, int> vector<D, T>::integer() const
{
  vector<D, int> c;

  for(int k=0; k<D; k++)
    c[k] = (int)x[k];

  return c;
}

template <int D, typename T>
inline vector<D, int> vector<D, T>::integer(const vector<D, T>& v)
{
  return v.integer();
}


// double
// ~~~~~~~
template <int D, typename T>
inline vector<D, double> vector<D, T>::Double() const
{
  vector<D, double> c;

  for(int k=0; k<D; k++)
    c[k] = (double)x[k];

  return c;
}

template <int D, typename T>
inline vector<D, double> vector<D, T>::Double(const vector<D, T>& v)
{
  return v.Double();
}


// []
// ~~
template <int D, typename T>
inline T& vector<D, T>::operator[](const unsigned int i)
{
  return x[i];
}


// Dot
// ~~~
template <int D, typename T>
inline double vector<D, T>::dot(const vector<D, T> &a) const
{
  double d=0;

  for(int k=0; k<D; k++)
    d += x[k] * a.x[k];

  return d;
}

template <int D, typename T>
inline double vector<D, T>::dot(const vector<D, T> &a, const vector<D, T> &b)
{
  return a.dot(b);
}


// NormSquared
// ~~~~~~~~~~~
template <int D, typename T>
inline double vector<D, T>::norm_squared() const
{
  return dot(*this, *this);
}

template <int D, typename T>
inline double vector<D, T>::norm_squared(const vector<D, T>& v)
{
  return v.norm_squared();
}


// read
// ~~~~
template <int D, typename T>
void vector<D, T>::read(std::ifstream& in)
{
  in.read((char*)x, sizeof(T)*D);
}

// write
// ~~~~~
template <int D, typename T>
void vector<D, T>::write(std::ofstream& out) const
{
  out.write((const char*)x, sizeof(T)*D);
}


// Insertion
// ~~~~~~~~~
template <int D, typename T>
std::ostream& operator<<(std::ostream& os, const vector<D, T>& v)
{
  os << "(";

  for(int k=0; k<D-1; k++)
    os << v.x[k] << ", ";

  os << v.x[D-1] << ")";

  return os;
}


// ======================================================================
// vector_field 
// ======================================================================

// A field of V-vectors on a D dimensional manifold

template<int V, int D, typename T=double>
class vector_field {
  
 public:
  int elements;

 private:
  vector<V, T>* f;
  vector<D, int> size;           // number of grid points for each dimension
  vector<D, int> offset;
 
 public:

  vector_field();
  vector_field(const vector<D, int>&);
  ~vector_field();

  vector<D, int> get_size() const;
  void set_size(const vector<D, int>&);

  vector<V, T>& get(const vector<D, int>&);

  void read(std::ifstream&);
  void write(std::ofstream&) const;

  static void swap(vector_field<V, D, T>&, vector_field<V, D, T>&);
};


// vector_field
// ~~~~~~~~~~~~
template<int V, int D, typename T>
vector_field<V, D, T>::vector_field()
  : f(0), elements(0)
{
}


// vector_field
// ~~~~~~~~~~~~
template<int V, int D, typename T>
vector_field<V, D, T>::vector_field(const vector<D, int>& s)
  : f(0)
{
  set_size(s);
}

// ~vector_field
// ~~~~~~~~~~~~~
template <int V, int D, typename T>
vector_field<V, D, T>::~vector_field()
{
  if(f != 0)
    delete[] f;
}

// get_size
// ~~~~~~~~
template<int V, int D, typename T>
inline vector<D, int> vector_field<V, D, T>::get_size() const
{
  return size;
}

// set_size
// ~~~~~~~~
template<int V, int D, typename T>
void vector_field<V, D, T>::set_size(const vector<D, int>& s)
{
  if(f != 0)
    delete[] f;

  size = s;

  elements = 1;
  for(int i=0; i<D; i++) {
    offset[i] = elements;
    elements *= size.x[i];
  }

  f = new vector<V, T>[elements];
}

// get
// ~~~
template<int V, int D, typename T>
inline vector<V, T>& vector_field<V, D, T>::get(const vector<D, int>& pos)
{
  int p=0;
  for(int i=0; i<D; i++)
    p += pos.x[i]*offset[i];

  return f[p];
}


// read
// ~~~~
template<int V, int D, typename T>
void vector_field<V, D, T>::read(std::ifstream& in)
{
  in.read((char*)f, elements*sizeof(T)*V);
}


// write
// ~~~~~
template<int V, int D, typename T>
void vector_field<V, D, T>::write(std::ofstream& out) const
{
  out.write((const char*)f, elements*sizeof(T)*V);
}

// swap
// ~~~~
template<int V, int D, typename T>
void vector_field<V, D, T>::swap(vector_field<V, D, T>& v1,
				 vector_field<V, D, T>& v2)
{
  vector<V, T>* f;

  f = v1.f;
  v1.f = v2.f;
  v2.f = f;
}


#endif 
```
