## Supplementary material for "Small subpopulations of β-cells do not drive islet oscillatory [Ca^2+^] dynamics via gap junction communication": S1 File: neighbor.C

```
#include "vector.h"
#include <iostream>
#include "box.h"


//==============================================================
//==============================================================
//  Class neighbor
//==============================================================
//==============================================================

neighbor::neighbor(int i_i)
 : i(i_i)
{
}


//==============================================================
//==============================================================
//  Class collision
//==============================================================
//==============================================================

collision::collision(int i_i, box *b_i)
  : neighbor(i_i),
    b(b_i)
{
  ctime = dblINF;
  cpartner = i;
}


//==============================================================
// Operation is finding the next collision from a given cell
//==============================================================
void collision::Operation(int j, vector<DIM, int>& pboffset)
{
  b->PredictCollision(i, j, pboffset, ctime, cpartner, cpartnerpboffset);
}
```
