## Supplementary material for "Small subpopulations of β-cells do not drive islet oscillatory [Ca^2+^] dynamics via gap junction communication": S1 File: read me.docx

Follows is a description for how we run the islet model. This model is run on the University of Colorado JANUS cluster where *BetaCell0.sh* contains the following shell script used. The model requires the boost C++ library (<http://www.boost.org/>) and sphere packing algorithm (Skoge et al., 2006)

**Set up the model in the JANUS environment:**

```
#!/bin/bash
#Setting the name of the job
#PBS -N Beta_Cell_Cluster
#Setting a walltime for the job
#PBS -l walltime=1:00:00
#Selecting processors
#PBS -l nodes=1:ppn=12
#SBATCH --reservation=janus-serial
cd $PBS_O_WORKDIR
```

This sets up the JANUS cluster environment by setting the number of processors, runtime, resource usage and job name.

**Build the islet:**

g++ neighbor.C spheres.C box.C sphere.C event.C heap.C read_input.C -o SpherePacking

./SpherePacking sphereInput

g++ GenerateSphere -o IsletGenerate

./IsletGenerate isletInput

First the code compiles the sphere packing code and sets up the islet using files in the *SpherePacking* directory. First the sphere packing files (*neighbor.c, spheres.c, box.c, sphere.c, event.c, heap.c, read_input.c*) are compiled using g++, to create a random sphere packing routine where the number of hard spheres in set in the sphereInput file. The code then compiles the islet generator (from *GenerateSpheres.cpp*) to create an islet of a custom size and corresponding connectivity matrix, where isletInput sets the number of cells in the islet. Other files in the *SpherePacking* directory (*stats.dat, write.dat, box.h, event.h, grid_field.h, heap.h, InCircle.h, my_vector.h, read_input.h, sphere.h, vector,h, NN10A.txt, XYZpos.txt*) are needed for running the sphere routines and are called in the above files, where *spheres* is generated upon compiling.

Note: This process uses a random number generator to create a new islet upon each time of running. Therefore it was only used when setting up a new islet in order to examine variability in simulating a set of conditions.

**Set heterogeneity for the islet:**

g++ -std=c++0x -I /directory_to_boost -fopenmp RandomVars.cpp -o RandomGenerator

./RandomGenerator

The *RandomVars.cpp* is compiled using g++ to create the executable file *RandomGenerator*. This is run which creates the file *RandomVars.txt* file which contains the set of heterogeneous parameters for each cell in the islet.

**Generate results files, construct islet and set initial cell parameters:**

g++ -I /directory_to_boost -fopenmp MainFile.cpp -o Beta

*MainFile.cpp* loads the initial conditions (*vars5exo.txt*) and sets up the general model ODE solver (*Beta*). This is called in *BetaCell.h* where the main islet model code is located.

**Run the model:**

./Beta 0B >> time_output0.txt

This code runs the model (*Beta*) and outputs the time step. The designated input file (*0B*) names the various output files that contain the output variables (e.g. membrane potential, [Ca^2+^]).

Other files include *ChR2Class.h* and *ChR2Class.cpp* which contain model code for an optional Channelrhodopsin class that was not utilized in the islet model for this study.
