## Supplementary material for "Small subpopulations of β-cells do not drive islet oscillatory [Ca^2+^] dynamics via gap junction communication": S1 File: read_input.C

```
#include <iomanip>
#include <iostream>
#include <stdio.h>
#include <fstream>

#include "read_input.h"

//================================================================
//
// Source File for input
//
//================================================================
int read_input::read(int argc, char * argv[])
{
  int error = 0;
  if (argc != 2) 
    {
    std::cout << "Syntax: spheres input" << std::endl;
    error = 1;
    } 
  else 
    {
    std::ifstream infile;
    infile.open(argv[1]);
    if(!infile)
      {
	std::cout << "Can't open " << argv[1] << " for input." << std::endl;
	error = 2;
	return error;
      } 
    else 
      {
	std::cout << "Reading input from file " << argv[1] << std::endl;
      }
    char buf[100],c;
    infile.get(buf,100,'='); infile.get(c); infile >> eventspercycle;
    infile.get(buf,100,'='); infile.get(c); infile >> N;
    infile.get(buf,100,'='); infile.get(c); infile >> initialpf;
    infile.get(buf,100,'='); infile.get(c); infile >> maxpf;
    infile.get(buf,100,'='); infile.get(c); infile >> temp;
    infile.get(buf,100,'='); infile.get(c); infile >> growthrate;
    infile.get(buf,100,'='); infile.get(c); infile >> maxpressure;
    infile.get(buf,100,'='); infile.get(c); infile >> maxcollisionrate;
    infile.get(buf,100,'='); infile.get(c); infile >> bidispersityratio;
    infile.get(buf,100,'='); infile.get(c); infile >> bidispersityfraction;
    infile.get(buf,100,'='); infile.get(c); infile >> massratio;
    infile.get(buf,100,'='); infile.get(c); infile >> hardwallBC;
    infile.get(buf,100,'='); infile.get(c); 
    infile.width(NAME_LEN-1); infile >> readfile;
    infile.get(buf,100,'='); infile.get(c); 
    infile.width(NAME_LEN-1); infile >> writefile;
    infile.get(buf,100,'='); infile.get(c); 
    infile.width(NAME_LEN-1); infile >> datafile;

    if(infile.eof()) 
      {
	std::cout << "Error reading input file " << argv[1] << std::endl;
	error = 3;
      }
    std::cout << "   eventspercycle : " << eventspercycle << std::endl;
    std::cout << "   N : " << N << std::endl;
    std::cout << "   initialpf : " << initialpf << std::endl;
    std::cout << "   maxpf : " << maxpf << std::endl;
    std::cout << "   temp : " << temp << std::endl;
    std::cout << "   growthrate : " << growthrate << std::endl;
    std::cout << "   maxpressure : " << maxpressure << std::endl;
    std::cout << "   maxcollisionrate : " << maxcollisionrate << std::endl;
    std::cout << "   bidispersityratio : " << bidispersityratio << std::endl;
    std::cout << "   bidispersityfraction : " << bidispersityfraction << std::endl;
    std::cout << "   massratio : " << massratio << std::endl;
    std::cout << "   hardwallBC : " << hardwallBC << std::endl;
    std::cout << "   readfile : " << readfile << std::endl;
    std::cout << "   writefile : " << writefile << std::endl;
    std::cout << "   datafile : " << datafile << std::endl;
    }
  return error;
}
```
