## Supplementary material for "Small subpopulations of β-cells do not drive islet oscillatory [Ca^2+^] dynamics via gap junction communication": S1 File: read_input.h

```
#ifndef READ_INPUT_H
#define READ_INPUT_H

#define NAME_LEN 256


class read_input {

public:

  int eventspercycle;          // # events per particle per cycle 
  int N;                        // # spheres
  double initialpf;               // initial packing fraction
  double maxpf;            // maximum packing fraction
  double temp;                      // initial temperature (temp=0 means v=0)
  double growthrate;               
  double maxpressure;         
  double maxcollisionrate;    
  double bidispersityratio;         // ratio of sphere radii for bidisperse 
  double bidispersityfraction;      // fraction of larger spheres
  double massratio;                 // ratio of sphere masses
  int hardwallBC;                   // =0 for periodic, =1 for hard walls
  char readfile[NAME_LEN];    // file with configuration; if new, creates new
  char writefile[NAME_LEN];    // file to write configuration
  char datafile[NAME_LEN];       // file to write statistics

  int read(int argc, char* argv[]);
 
};


#endif
```
