## Supplementary material for "Small subpopulations of β-cells do not drive islet oscillatory [Ca^2+^] dynamics via gap junction communication": S1 File: RunSimulation.sh

```
#!/usr/bin/env bash
#BSUB -J BetaCellModel
#BSUB -n 4
#BSUB -R "select[mem>10] rusage[mem=10] span[hosts=1]"
#BSUB -o stdout_%J.out
#BSUB -e stderr_%J.err

#Execute

rm potential.txt sodium.txt potassium.txt calcium.txt caer.txt atp.txt adp.txt IRP.txt PP.txt DP.txt FIP.txt RIP.txt Cap.txt time.txt noise.txt
rm std*
#rm spheres Islet RandomGenerator Beta

#compile sphere and generate sphere input file
g++ neighbor.C spheres.C box.C sphere.C event.C heap.C read_input.C -o spheres 
./spheres sphereInput

#create islet
g++ -I /vol2/home/dwuletj/boost/boost_1_58_0/ -fopenmp GenerateSphere.cpp -o Islet
./Islet isletInput

#compile and general random numbers
g++ -std=c++0x -I /vol2/home/dwuletj/boost/boost_1_58_0/ -fopenmp RandomVars.cpp -o RandomGenerator
./RandomGenerator

#run main code for solving differential equations
g++ -I /vol2/home/dwuletj/boost/boost_1_58_0/ -fopenmp MainFile.cpp -o Beta
./Beta 0B 
```
