## Supplementary material for "Small subpopulations of β-cells do not drive islet oscillatory [Ca^2+^] dynamics via gap junction communication": S1 File: sphere.C

```
#include "box.h"
#include <iostream>
#include <math.h>
#include <stdlib.h>
#include <time.h>

#include "vector.h"

//==============================================================
//==============================================================
//  Class Sphere: 
//==============================================================
//==============================================================


//==============================================================
// Constructor
//==============================================================
sphere::sphere()
{
}


//==============================================================
// Constructor
//==============================================================
sphere::sphere(const sphere& s)
{
  i = s.i;
  x = s.x;
  v = s.v;
  cell = s.cell;
  lutime = s.lutime;
  nextevent = s.nextevent;
  nextcollision = s.nextcollision;
  r = s.r;
  gr = s.gr;
  m = s.m;
  species=s.species;
}

//==============================================================
// Constructor
//==============================================================
sphere::sphere(int i_i, vector<DIM> x_i, vector<DIM, int> cell_i, 
	       double lutime_i, double r_i, double gr_i, double m_i, int species_i):
  i(i_i),
  x(x_i),
  cell(cell_i),
  lutime(lutime_i),
  r(r_i),
  gr(gr_i),
  m(m_i),
  species(species_i)
{
}

//==============================================================
// Destructor
//==============================================================
sphere::~sphere() 
{

}

 
```
