## Supplementary material for "Small subpopulations of β-cells do not drive islet oscillatory [Ca^2+^] dynamics via gap junction communication": S1 File: sphere.h

```
#ifndef  SPHERE_H
#define  SPHERE_H


#include "vector.h"


class sphere {

 public:

  // constructor and destructor

  sphere();
  sphere(const sphere& s);
  sphere(int i_i, vector<DIM> x, vector<DIM, int> cell_i, double lutime_i, 
	 double r_i, double gr_i, double m_i, int species_i);
  ~sphere();

 //variables
  
  int i;                          // sphere ID

  // impending event
  event nextevent;                // next event...can be collision or transfer
  event nextcollision;            // next collision if next event is transfer
  // maybe nextnext event
  
  // past information
  double lutime;                  // last update time
  vector<DIM, int> cell;          // cell that it belongs to
  vector<DIM, double> x;          // position
  vector<DIM, double> v;          // velocity
  double r;                       // sphere radius
  double gr;                      // sphere growth rate
  double m;                       // sphere mass
  int species;                    // species number (not used during the MD)
  // make sure efficent in memory

 

};

#endif 
```
