## Supplementary material for "Small subpopulations of β-cells do not drive islet oscillatory [Ca^2+^] dynamics via gap junction communication": S1 File: spheres.C

```
//===========================================================
//===========================================================
//===========================================================
//
//  Molecular dynamics simulation of hardspheres
//
//===========================================================
//===========================================================
//===========================================================

#include <iostream>
#include <math.h>
#include <fstream>
#include <vector>
#include <time.h>
#include <string.h>
#include <cstdlib>
#include "box.h"
#include "sphere.h"
#include "event.h"
#include "heap.h"
#include "read_input.h"


int main(int argc, char **argv)
{
  read_input input;
  int error = input.read(argc, argv);
  if (error) return error;

  double d, r;   // initial diameter and radius of spheres

  if(strcasecmp(input.readfile, "new")==0)
    input.readfile[0]=0;

  if (input.readfile[0]) // read in existing configuration
    {
      // read the header
      std::ifstream infile(input.readfile);
      if (!infile)
	{
	  std::cout << "error, can't open " << input.readfile  << std::endl;
	  exit(-1);
	}
      else
	{
	  int dim;
	  infile >> dim; infile.ignore(256, '\n');
	  if (dim != DIM)  // quit if dimensions don't match
	    {
	      std::cout << "error, dimensions don't match" << std::endl;
	      exit(-1);
	    }
	  infile.ignore(256, '\n');  // ignore the N 1 line
	  infile >> input.N; infile.ignore(256, '\n');
	  std::cout << "N = " << input.N << std::endl;
	  infile >> d; infile.ignore(256, '\n');
	  std::cout << "d = " << d << std::endl;
	  r = d/2.;
	  std::cout << "r = " << r << std::endl;
	}
    }
  else // create a new configuration
    {
      r = pow(input.initialpf*pow(SIZE, DIM)/(input.N*VOLUMESPHERE), 1.0/((double)(DIM)));
    }

  box b(input.N, r, input.growthrate, input.maxpf, input.bidispersityratio, 
	input.bidispersityfraction, input.massratio, input.hardwallBC);
  
  std::cout << "ngrids = " << b.ngrids << std::endl;
  std::cout << "DIM = " << DIM << std::endl;

  if(input.readfile[0])
    {
      std::cout << "Reading in positions of spheres" << std::endl;
      b.RecreateSpheres(input.readfile, input.temp);
    }
  else 
    {
      std::cout << "Creating new positions of spheres" << std::endl;
      b.CreateSpheres(input.temp);
    } 
  
  std::ofstream output(input.datafile);
  output.precision(16);  

  
  while ((b.collisionrate < input.maxcollisionrate) && (b.pf < input.maxpf) && (b.pressure < input.maxpressure)) 
    {
      b.Process(input.eventspercycle*input.N);
      output << b.pf << " " << b.pressure << " " << 
	b.collisionrate << " " << b.neventstot << " " << std::endl;

      b.Synchronize(true);
    }
  
  output.close();

  b.WriteConfiguration(input.writefile);
  std::cout << "b.pf = " << b.pf << std::endl;
  std::cout << "b.pressure = " << b.pressure << std::endl;
  std::cout << "b.collisionrate = " << b.collisionrate << std::endl;
  
  
  
  return 0;
}
```
