## Supplementary material for "Small subpopulations of β-cells do not drive islet oscillatory [Ca^2+^] dynamics via gap junction communication": S1 File: RandomVars.cpp

```
#include "BetaCell.h"
#include "boost/random/random_device.hpp"
#include <random>
#include <ctime>
#include <boost/random/mersenne_twister.hpp>
#include <boost/random/uniform_real_distribution.hpp>
#include <boost/random/variate_generator.hpp>
#include <cmath>
//Randomizes cellular properties and saves to file
int main()
{
	srand (4);

//Initialize random variable matrix
double randomVars[cellNumber][12];
double cSum=0;

int sortgCoup = 0; //set to 1 to sort Coup array
int sortgKATP = 0; //set to 1 to sort gKATP
int sortkGlyc = 0; //set to 1 to sort kGlyc

double gCoupVarSort[cellNumber];
double gKATPVarSort[cellNumber];
double kGlycVarSort[cellNumber];

int numofhubs = (int) round(cellNumber * hubMutant);
boost::mt19937 gen(4);
boost::random::uniform_real_distribution<> dis(0.25, 1.5);
//boost::mt19937 rng; // 

//most of these distributions are 10% variation but we change them for various conditions to 25% or 50% variation
//I used 25% varation in Glyc for subpopulation simulations 
//I also used 50% variation in gKATP and GLyc for subpopulation simulations JD

boost::normal_distribution<> gKATPv_hub(2.35, 0.0235); //normal 25%
boost::normal_distribution<> gKATPv(2.29, 0.0229); //normal 25%
//boost::normal_distribution<> gKATPv(2.31,1.155); //subpopulation 50% variation

//boost::gamma_distribution<>gCoupS(4,4); //Set coupling strengths here
//boost::gamma_distribution<>gCoupS(4,4); //Set coupling strengths here
boost::gamma_distribution<>gCoupS(4,4); //Set coupling strengths here

boost::normal_distribution<> gKtoarv_hub(2.125, 0.02125);
boost::normal_distribution<> gKtoarv(2.128, 0.02128);

//boost::normal_distribution<> PCaERarv(0.096, .0096);
boost::normal_distribution<> PCaERarv_hub(0.097, 0.00097);
boost::normal_distribution<> PCaERarv(0.096, 0.00096);

//boost::normal_distribution<> pKATPv(1.0,0.000);

boost::normal_distribution<> gGKCaBKv_hub(2.134,0.02134);
boost::normal_distribution<> gGKCaBKv(2.125,0.02125);

boost::normal_distribution<> PNACAv_hub(205,2);
boost::normal_distribution<> PNACAv(204,2);

boost::normal_distribution<> Prelv_hub(0.45,0.0045);
boost::normal_distribution<> Prelv(0.46,0.0046);

boost::normal_distribution<> Popv_hub(0.000497,0.00000497);
boost::normal_distribution<> Popv(0.000500,0.00000500);
 
boost::normal_distribution<> ATPv_hub(3.998,0.03998);
boost::normal_distribution<> ATPv(4.003,0.04003);
//boost::normal_distribution<> KBOXv(0.0000063,0.0000006);
 boost::normal_distribution<> KBOXv(0.0000063,0.0000006);

//boost::normal_distribution<> GLYCv(0.000126,0.000063); //50% variation for subpopulations
//boost::normal_distribution<> GLYCv(0.000126,0.0000315); //25% Variation used for the subpopulation stuff
//boost::normal_distribution<> GLYCv(0.000126,0.0000126); %normal 10% variation
boost::normal_distribution<> GLYCv_hub(0.000106,0.00000106);//simulate hubs to have same top 10% mean and followers to have bottom 90% mean with small variation.
boost::normal_distribution<> GLYCv_fol(0.000129,0.00000129); //sigma is sigma/4

for (int i = 0; i<cellNumber; i++)
{
	//Calling the random number generators from BetaCell.h
	if (i < numofhubs) { 
		randomVars[i][0] = gKATPv_hub(gen);
		while (randomVars[i][0]<0)
		{
			randomVars[i][0] = gKATPv_hub(gen);
		}
		randomVars[i][1] = gCoupS(gen);
		while (randomVars[i][1]<0)
		{
			randomVars[i][1] = gCoupS(gen);
		}
//		cSum = cSum + randomVars[i][1];
		randomVars[i][2] = gKtoarv_hub(gen);
		while (randomVars[i][2]<0)
		{
			randomVars[i][2] = gKtoarv_hub(gen);
		}
		randomVars[i][3] = PCaERarv_hub(gen);
		while (randomVars[i][3]<0)
		{
			randomVars[i][3] = PCaERarv_hub(gen);
		}
		randomVars[i][4] = gGKCaBKv_hub(gen);
		while (randomVars[i][4]<0)
		{
			randomVars[i][4] = gGKCaBKv_hub(gen);
		}
		randomVars[i][5] = PNACAv_hub(gen);
		while (randomVars[i][5]<0)
		{
			randomVars[i][5] = PNACAv_hub(gen);
		}
		randomVars[i][6] = Prelv_hub(gen);
		while (randomVars[i][6]<0)
		{
			randomVars[i][6] = Prelv_hub(gen);
		}
		randomVars[i][7] = Popv_hub(gen);
		while (randomVars[i][7]<0)
		{
			randomVars[i][7] = Popv_hub(gen);
		}
		randomVars[i][8] = ATPv_hub(gen);
		while (randomVars[i][8]<0)
		{
			randomVars[i][8] = ATPv_hub(gen);
		}
	} else {
		randomVars[i][0] = gKATPv(gen);
		while (randomVars[i][0]<0)
		{
			randomVars[i][0] = gKATPv(gen);
		}
		randomVars[i][1] = gCoupS(gen);
		while (randomVars[i][1]<0)
		{
			randomVars[i][1] = gCoupS(gen);
		}
//		cSum = cSum + randomVars[i][1];
		randomVars[i][2] = gKtoarv(gen);
		while (randomVars[i][2]<0)
		{
			randomVars[i][2] = gKtoarv(gen);
		}
		randomVars[i][3] = PCaERarv(gen);
		while (randomVars[i][3]<0)
		{
			randomVars[i][3] = PCaERarv(gen);
		}
		randomVars[i][4] = gGKCaBKv(gen);
		while (randomVars[i][4]<0)
		{
			randomVars[i][4] = gGKCaBKv(gen);
		}
		randomVars[i][5] = PNACAv(gen);
		while (randomVars[i][5]<0)
		{
		randomVars[i][5] = PNACAv(gen);
		}
		randomVars[i][6] = Prelv(gen);
		while (randomVars[i][6]<0)
		{
			randomVars[i][6] = Prelv(gen);
		}
		randomVars[i][7] = Popv(gen);
		while (randomVars[i][7]<0)
		{
			randomVars[i][7] = Popv(gen);
		}
		randomVars[i][8] = ATPv(gen);
		while (randomVars[i][8]<0)
		{
			randomVars[i][8] = ATPv(gen);
		}
}

/*	randomVars[i][9] = GLYCv(gen);
	while (randomVars[i][9]<0)
	{
		randomVars[i][9] = GLYCv(gen);
	}*/
	if (i < numofhubs){
		randomVars[i][9] = GLYCv_hub(gen);
		while (randomVars[i][9]<0)
		{
			randomVars[i][9] = GLYCv_hub(gen);
		}
	} else {
		randomVars[i][9] = GLYCv_fol(gen);
		while (randomVars[i][9]<0)
		{
			randomVars[i][9] = GLYCv_fol(gen);
		}
	}

	//Unique random number generator to help determine coupling conductance.
	randomVars[i][10] = rand() % 1000000;
	randomVars[i][11] = dis(gen);

	gKATPVarSort[i] = randomVars[i][0];
	gCoupVarSort[i] = randomVars[i][1];
	kGlycVarSort[i] = randomVars[i][9];
}


if (std::abs(sortgKATP)==1)
{
	std::sort(gKATPVarSort, gKATPVarSort+cellNumber);
	if (sortgKATP < 0)
	{
		std::reverse(gKATPVarSort, gKATPVarSort+cellNumber);
	//want this is lowest to highest order because low gKATP is better for cells
	}
}

if (std::abs(sortgCoup)==1)
{
	std::sort(gCoupVarSort, gCoupVarSort+cellNumber);
	if (sortgCoup==1)
	{
		std::reverse(gCoupVarSort, gCoupVarSort+cellNumber);
	}
}
if (std::abs(sortkGlyc)==1)
{
	std::sort(kGlycVarSort, kGlycVarSort+cellNumber);
	if (sortkGlyc==1)
	{
		std::reverse(kGlycVarSort, kGlycVarSort+cellNumber);
	}	
}

// first hub cells simulations used this...
for(int i=0;i<cellNumber;i++)
{
	randomVars[i][0]=gKATPVarSort[i];
	randomVars[i][1]=gCoupVarSort[i];
	randomVars[i][9]=kGlycVarSort[i];

	if (i < numofhubs){
		//hub values
		randomVars[i][1]=gCoupVarSort[i];
		//randomVars[i][9] = kGlycVarSort[i]/1.25*5.5*2.0; //just move hub values to be higher
	} else {
		randomVars[i][1]=gCoupVarSort[i];
		//randomVars[i][9] = kGlycVarSort[i]/1.25; //make follower values less
	}
	cSum = cSum + randomVars[i][1];
}

std::shuffle(std::begin(randomVars),&randomVars[numofhubs], gen);
std::shuffle(&randomVars[numofhubs], std::end(randomVars), gen);//shuffle follower cells only

double cMean=cSum/cellNumber;

////////////////Below sets the average coupling conductance///////////////
for(int i=0;i<cellNumber;i++)
{
randomVars[i][1]=randomVars[i][1]*0.12/cMean;
}
///////////////////////////////////////////////////////////////////
remove("RandomVars.txt");

//Write variables to file RandomVars.txt
ofstream outFile;
outFile.open("RandomVars.txt");
for(int i=0;i<cellNumber;i++)
{
for(int j=0;j<=11;j++)
{
outFile<<randomVars[i][j]<<' ';
}
outFile<<' '<<endl;
}

return 0;
}
```
