## Supplementary material for "Small subpopulations of β-cells do not drive islet oscillatory [Ca^2+^] dynamics via gap junction communication": S1 Table

| Parameter | Description of parameter | Mean 'low phase' cell population | Mean 'non-low phase' cell population, | units |
|---|---|---|---|---|
| $g_{KATP}$ | Max conductance of $K_{ATP}$ channel current | 2.3517 | 2.2930 | pA $mV^{-1}$ |
| $g_{KTO}$ | Conductance of $I_{KCa(BK)}$ (voltage and $Ca^{2+}$) dependent transient outward $K^{+}$ current | 2.12521 | 2.12793 | pA $mV^{-1}$ |
| $P_{SERCA}$ | Maximum rate of pumping $Ca^{2+}$ into ER | 0.09666 | 0.09586 | amole $ms^{-1}$ |
| $P_{NaCa}$ | Maximum amplitude of $I_{NaCa}$, $Na^{+}/Ca^{2+}$ exchanger | 204.98 | 203.84 | pA |
| $P_{rel}$ | Converting factor for $Ca^{2+}$ release from ER | 0.4548 | 0.4609 | fl $ms^{-1}$ |
| $P_{op}$ | Maximum rate of ATP production from oxphos | 0.00049742 | 0.00049957 | $ms^{-1}$ |
| $[ATP_{tot}]$ | Total amount of ATP species | 3.99813 | 4.00341 | mM |
| $k_{glc}$ | Rate constant of glycolysys | 0.0001064 | 0.0001286 | $ms^{-1}$ |
